# Unavoidable Disasters: the Hsp70 Chaperone Can Only Prevent Aggregation Over a Limited Time Span

**DOI:** 10.64898/2026.11.09.765239

**Authors:** Miranda F. Mecha, Matías J. Dahl, Jasmine K. Machhi, Samuel C. Carey, Hanming Yang, Xi Chen, Silvia Cavagnero

## Abstract

Cells must attain and maintain proper protein folding and avoid aggregation: this is a pervasive challenge across all domains of life. Molecular chaperones are central to this process, acting through mechanisms shaped by the free-energy landscapes of their client proteins. The impact of chaperones on client-specific kinetic and thermodynamic characteristics, however, remains poorly characterized. Here, we develop the CHAMPION70 computational model to perform kinetic simulations on the *E. Coli* Hsp70 chaperone system in the presence of client-protein folding and aggregation. Experimentally determined folding/unfolding/aggregation rate constants are taken into account. In the case of weakly aggregation-prone clients, CHAMPION70 predicts that Hsp70 prevents aggregation only over timescales approaching the average human lifespan (79 years). For strongly aggregation-prone clients, however, aggregation inevitably occurs starting from a few minutes, even at higher-than-physiological Hsp70 concentrations. Thus, the Hsp70 chaperone system merely delays, yet surprisingly does not eliminate, the onset of aggregation. Further, CHAMPION70 predicts that Hsp70 most effectively retards the aggregation of slow-folding, thermodynamically stable clients. In contrast, thermodynamically unstable clients remain engaged with Hsp70 for prolonged periods even when aggregation propensity is low, thereby consuming chaperone capacity and reducing the extent of productive folding. This outcome suggests that condensate formation by proteins containing disordered regions may serve to mitigate unnecessary Hsp70 engagement. In all, our results provide a quantitative framework for understanding how client-specific kinetic and thermodynamic properties shape chaperone activity and show that the Hsp70 chaperone system does not altogether prevent aggregation: it merely retards it.

## INTRODUCTION

Molecular chaperones are essential for maintaining proteostasis in all organisms. They promote *de novo* protein folding, prevent misfolding and aggregation, and assist in the recovery of stress-damaged proteins.^1–4^ As highly conserved components of the proteostasis network, chaperones are indispensable for sustaining a functional proteome across all forms of life.^5^ For instance, evolutionary studies suggest that even the last universal-common-ancestor (LUCA) organism included at least one molecular chaperone (Hsp60, or a GroEL-like chaperonin).^6^ Further, even the most highly simplified version of a bacterium, developed by Craig Venter’s team in 2016 and featuring a minimal genome, includes genetic encoding for the DnaK, DnaJ and GrpE chaperones.^7^ This trio of molecular helpers is collectively known as the Hsp70 chaperone system.

Among the many molecular chaperones across bacteria and eukaryotes, the 70-kDa heat shock protein family (Hsp70) is particularly important. For instance, in *E. coli*, simultaneous deletion of the genes encoding for the Hsp70 and trigger factor chaperones leads to cell death, at temperatures higher than 20 °C.^8–10^ Loss of DnaK leads to growth defects, temperature sensitivity, and to the accumulation of misfolded and aggregated proteins,^11,12^ underscoring the essential role of Hsp70 even under non-stress conditions. Other chaperone pathways contribute to folding quality control. Yet, none of them fully compensates for the absence of DnaK.^12^ As a result, DnaK plays a uniquely important role in maintaining a functional bacterial proteome.

Hsp70 is ubiquitous, and genes encoding Hsp70 chaperones are highly conserved across organisms.^13–15^ Hsp70 chaperones contain two domains that are allosterically regulated via nucleotide-binding (Fig. 1A, B). DnaK, i.e., the Hsp70 chaperone from *E. coli*, consists of two domains comprising a 45-kDa nucleotide-binding domain (NBD) and a 25-kDa substrate-binding domain (SBD).^16^ The NBD mediates allosteric conformational changes controlled by the nucleotide state. These effects dramatically influence the affinity of the chaperone for its client-protein.^17^ The SBD of Hsp70 preferentially interacts with 4-5 nonpolar client-protein residues, often aliphatic (Val, Leu, Ile) and sometimes aromatic (Phe, Tyr), typically flanked by short (ca. 4 amino acids) stretches of positively charged residues.^18,19^

**Fig. 1.**
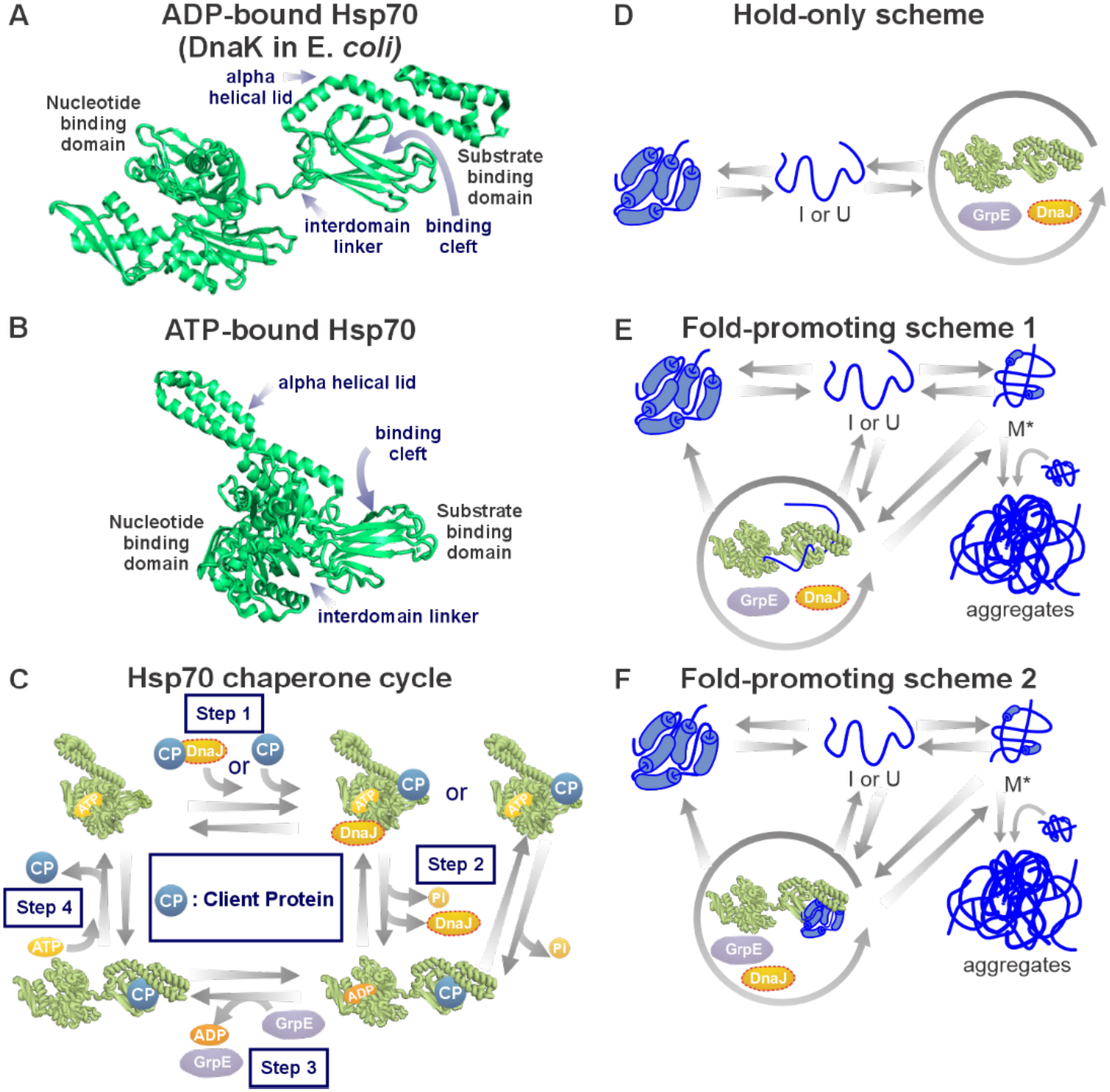
Hsp70 chaperone structure, functional cycle, and client-specific folding outcomes. **A.** ADP-bound *E. coli* Hsp70 (DnaK), PDB: 2KHO (Bertelsen *et al.*, 2019). The structure includes the nucleotide-binding domain, substrate-binding domain, interdomain linker, and α-helical lid. The ADP-bound conformation resembles the nucleotide-free state. **B.** ATP-bound *E. coli* Hsp70 (DnaK), PDB: 4B9Q (Kityk *et al.*, 2012). The lid adopts an open conformation, exposing the substrate-binding cleft. **C.** Schematic of the *E. coli* Hsp70 chaperone cycle showing sequential steps of client binding and release regulated by cochaperones DnaJ and GrpE. ATP hydrolysis and nucleotide exchange drive transitions between low and high affinity states. **D.** According to a well-known non-fold-promoting scheme (hold-only), the client-protein binds Hsp70 and is released in a similar conformation to the one in which it initially binds. By transiently sequestering unfolded or intermediate states, Hsp70 lowers the free concentration of aggregation-prone species in solution, thereby reducing aggregation. **E.** According to a possible fold-promoting model (scheme 1), the client-protein is released from Hsp70 in a folding-competent state that can readily reach its native conformation after release. **F.** According to a possible fold-promoting model (scheme 2), the client-protein folds to its native-state while bound to Hsp70, and is subsequently released in solution in its native conformation.

Hsp70 chaperone activity involves a functional cycle that includes the Hsp40 (a J-protein) and nucleotide exchange factor (NEF) cochaperones.^20–22^ The Hsp70 (i.e., DnaK in *E. coli*) cycle proceeds via four main steps, as schematically illustrated in Fig. 1C. First, the nucleotide-free chaperone binds ATP resulting in a state with low affinity and fast exchange rates for client-proteins.^23^ During the second step, client-proteins either directly bind Hsp70 or are delivered to the chaperone via a prior interaction with a J-domain protein (i.e., DnaJ in *E. coli*).^23–25^ Then, ATP hydrolysis takes place, leading to the conversion of ATP to ADP and to the effective closure of the Hsp70 α-helical lid over the client-protein. The ADP-bound state of Hsp70 has a high affinity and slow exchange rates for the client-protein. During the third step, a NEF (GrpE in *E. coli*) causes the dissociation of ADP and the generation of a nucleotide-free state of client-protein-bound Hsp70. As part of the fourth and final step, ATP binding causes client-protein release via opening of the α-helical lid, thus enabling the cycle to start anew.^23^

A central question to understand the Hsp70 functional cycle is how client-proteins interact with the chaperone and how this interaction influences folding outcomes. Experimental and theoretical studies indicate that Hsp70 can operate through distinct modes of action that depend on the properties of the client-protein. For some clients, Hsp70 binds the protein and subsequently releases it in the same conformational state that the protein originally assumed upon interaction initiation. According to this non-fold-promoting model, Hsp70 effectively merely “holds” unfolded or misfolded proteins, reducing their free concentration in solution and therefore slowing down aggregation (Fig. 1D).^26–28, 29–35^ Consistent with this mechanism, ADP-or nucleotide-free-Hsp70 kinetics stabilizes unfolded and misfolded client-protein conformations.^33,36–38^ Experimental studies on the non-aggregation-prone (NAP) client-protein RNase H^D^ show an Hsp70 behavior consistent with this hold-only mechanism, according to which Hsp70 interaction slows the acquisition of native structure, relative to chaperone-free conditions.^39^

In contrast, for misfolding and aggregation-prone client-proteins, Hsp70 can actively promote folding of native structure through mechanisms consistent with the fold-promoting schemes shown in Fig. 1E and 1F. In these cases, Hsp70 facilitates the unfolding of misfolded states and accelerates conversion to the native conformation. For instance, Imamoglu *et al.* showed that both the yield and rate of bioactive native-state formation of firefly luciferase increase in the presence of Hsp70.^40^ At low luciferase concentrations (1 nM), where aggregation is negligible but ∼40% misfolded monomers get generated *in situ*, Hsp70 accelerates acquisition of the native-state by actively unfolding bound misfolded species, thereby removing intermediates that would otherwise promote aggregation at higher concentrations.^40^

To date, a few kinetic models were developed to simulate client-protein folding within cell-like environments in the presence of a variety of chaperones.^41–43^ A computational tool for predicting protein folding timecourse mediated specifically by the *E. coli* Hsp70 chaperone was developed by Tomita and coworkers.^44^ This model assumes that client-proteins cannot fold independently and that they are unable to bind nucleotide-free Hsp70.^45^ In addition, this algorithm does not take into account the fact that K/J/E complexes can exist both in the absence and presence of client-proteins.^20,45–47^ Another predictive tool developed by Powers and coworkers, FoldEco,^48^ takes the cellular protein-folding machinery into account more comprehensively than earlier models. FoldEco predicts increased aggregation in the absence of chaperones and makes insightful predictions about slow-folding and misfolding-prone proteins. This model, which incorporates several major *E. coli* chaperone systems, was validated against experimental data on heterologous firefly luciferase (fluc) expression and accurately reproduces *in vivo* folding outcomes.^48^ FoldEco is therefore a powerful tool for predicting proteostasis behavior in the presence of multiple chaperones. On the other hand, its representation of the Hsp70 (K/J/E) system is necessarily simplified and does not capture how Hsp70 engages different client-proteins through distinct kinetic and thermodynamic interaction modes, including whether a given client undergoes fold-promoting behavior or predominantly hold-only interactions. In addition, Sekhar *et al.* developed CHAMP70, a more detailed Hsp70-centric kinetic model tailored to predict the folding of NAP proteins in the absence and presence of Hsp70.^39^ The CHAMP70 kinetic model addresses several of the simplifying assumptions of previous models and, wherever possible, employs experimentally determined rate constants for specific steps of the Hsp70 chaperone cycle. CHAMP70 was validated against experimental data on the refolding of a non-aggregating client-protein (RNase H^D^)^49^ in the absence and presence of Hsp70.^39^ Unfortunately, none of the above models explains the experimental observations, which show that Hsp70 has opposite effects on folding kinetics depending on client aggregation propensities.^40,44,50^ Hsp70 slows down native-state formation of NAP proteins,^50^ consistent with a hold-only mechanism, but accelerates native-state acquisition of aggregation-prone clients through unfolding/refolding steps.^40^

In this work, we develop CHAMPION70, a computational model able to perform Chaperone-Mediated Protein folding kinetic Simulations involving Hsp70. CHAMPION70 has been validated against experimental data on NAP and aggregation-prone client-proteins with different kinetic and thermodynamic characteristics. We performed simulations in the presence of client-proteins with either zero, weak, or strong aggregation propensities. The CHAMPION70 simulations indicate that Hsp70 can delay, but not indefinitely prevent, aggregation for proteins with even modest misfolding tendencies, highlighting intrinsic limits to the protective capacity of the *E. coli* Hsp70 system. In addition, the model reveals that Hsp70 most effectively prevents aggregation for clients that fold slowly but are thermodynamically stable, while unstable clients remain persistently bound and cannot be efficiently rescued. Together, these results define the conditions under which Hsp70 succeeds or fails at maintaining protein solubility and clarify how client-dependent features influence overall folding and aggregation outcomes.

## Results and Discussion

### Kinetic modeling of folding and aggregation of two fundamentally distinct Hsp70 clients

The primary objective of this work is to develop a detailed kinetic model capable of predicting how the Hsp70 chaperone system modulates protein folding and aggregation. As a first step, we established a computational description of the intrinsic folding, misfolding, and aggregation behavior of two model client proteins in the absence of chaperones. The goal was to ensure that our aggregation framework accurately reproduces known experimental behaviors, before introducing the Hsp70 system. Given that Hsp70 interacts with clients spanning a wide range of aggregation propensities, we selected two well-characterized slow-folding and thermodynamically stable Hsp70 clients with experimentally distinct aggregation behaviors. Specifically, we examined the *E. Coli* Ribonuclease H* I53D variant (RNase H^D^),^49–51^ a small protein (Fig. 2A) that does not aggregate on a 2-3 min timescale at room temperature, and firefly luciferase (fluc), a larger and more complex client (Fig. 2F) that misfolds and aggregates under refolding conditions in the absence of chaperones.^40^ The above clients were chosen because extensive experimental data are available for both and, importantly, because Hsp70 affects the folding kinetics of these clients in fundamentally different ways. Namely, the Hsp70 chaperone (K/J/E) system experimentally slows down RNase H^D^ native-state formation. Conversely, K/J/E accelerates acquisition of the native-state in the case of fluc.

**Fig. 2.**
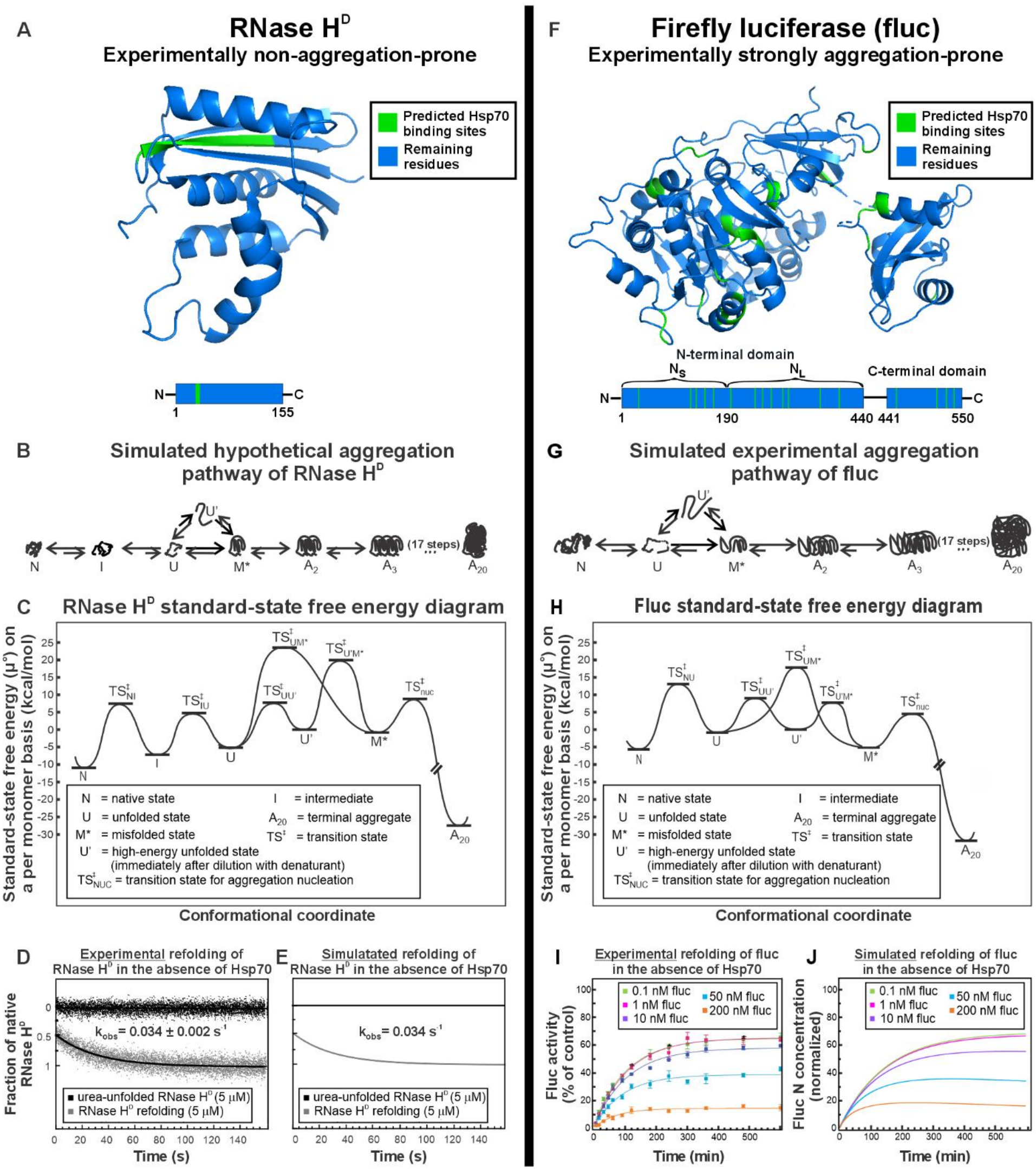
RNase H^D^ and firefly luciferase (fluc) folding in the absence of chaperones. **A.** Three-dimensional structure of *E. Coli* RNase H^D^ (PDB 1F21). The green region indicates the predicted Hsp70 binding site identified with the sequence-based algorithm by Rüdiger *et al.* (1997). **B.** Experimental stopped-flow refolding kinetics of RNase H^D^ (gray, 5 μM) and reference trace of the urea-unfolded state (black, 5 μM) according to the literature (Sekhar *et al.*, 2012). Data were collected by far-UV circular dichroism (222 nm) at 30° C. **C.** Simulation of experimental results of panel B according to the kinetic scheme in panel E, starting from a 100% population of U’ (5 μM). **D.** Kinetic scheme overview used for RNase H^D^ simulations. **E.** Standard-state chemical potential diagram (per-monomer) of the reaction scheme used to simulate the folding and hypothetical aggregation of RNase H^D^. **F.** Three-dimensional structure of *E. Coli* firefly luciferase (fluc) (PDB 1LCI). Green regions indicate the multiple predicted Hsp70 binding sites identified using the Rüdiger *et al.* sequence-based algorithm. **G.** Concentration-dependent refolding activity of fluc (0.1-200 nM) in the absence of chaperones, reproduced from Imamoglu *et al.* (2020). **H.** Simulation of the experimental results in panel G (0.1-200 nM). Fluc was refolded from the transiently expanded U’ state. **I.** Kinetic scheme overview used for fluc simulations. **J.** Free-energy diagram (per-monomer) of the reactions used for simulations in panel H.

Next, we generated folding/misfolding/aggregation kinetic schemes for RNase H^D^ (Fig. 2B) and fluc (Fig. 2G) according to the following criteria. First, we assumed that, immediately after dilution from denaturant, proteins occupy a conformationally unrelaxed (typically fairly extended)^52^ unfolded state denoted as U’. This state can either convert into a conformationally relaxed more stable unfolded state U or misfold to generate the aggregation-competent state M*. Second, we considered that the U state can then either fold to the native state N (e.g. fluc), convert to a folding intermediate I (e.g. RNase H^D^ burst-phase intermediate^50^), or misfold into M*. Third, we assumed that the M* state undergoes aggregation via primary nucleation forming a dimeric intermediate, followed by elongation via sequential addition of misfolded M* monomers. Folding/unfolding rate constants, as well as kinetic pathways of folding intermediates, were taken from experimental data. Given that RNase H^D^ and fluc differ in their intrinsic aggregation propensities, we introduced a general framework for describing aggregation. Specifically, we defined three aggregation regimes: a non-aggregation-prone (NAP) regime, a weakly aggregation-prone (WAP) regime, and a strongly aggregation-prone (SAP) regime.

The above aggregation regimes were implemented by modifying the rate constants of transitions impacting the buildup of the misfolded state M* (U⇌M*, U’⇌M*, U⇌U’). The NAP regime lacks transitions leading to M*, thus preventing M* formation. The WAP regime retains the full folding, misfolding, and aggregation network, but imposes high kinetic barriers for U⇌M* and U’⇌M*, limiting access to M*. The SAP regime lowers these barriers, deepening the M* basin and rendering the standard-state free-energy of M* similar to that of the native-state, thereby enabling rapid misfolding and aggregation. Kinetic rate constants for the NAP, WAP, and SAP regimes were chosen to satisfy the above criteria (Supplementary Table S4), and corresponding energy-landscape representations of these regimes are shown in Supplementary Fig. S12. Experimental solubility measurements in the *E. coli* cytosolic proteome support the existence of the above three classes, with approximately 20% of proteins behaving as NAP, 53% as WAP, and 26% as SAP (Dang *et al.*, in preparation).

Next, standard-state chemical-potential landscapes of RNase H^D^ and fluc were created, as shown in Figs. 2C and 2H, from experimentally determined folding/unfolding rate constants and from appropriate kinetic parameters for the aggregation regimes described above (see details in Supporting Materials). Kinetic modeling was implemented with the COPASI^53^ software, using the refolding/aggregation reaction schemes in Fig. 2B and 2G for RNase H^D^ (WAP aggregation conditions) and fluc (SAP aggregation conditions), respectively. Note that, in the case of RNase H^D^, we modeled the protein as refolding on short timescales (as experimentally observed^50^) and aggregating on long timescales.

Finally, using experimentally determined folding and unfolding rate constants together with WAP (RNase H^D^) and SAP (fluc) aggregation-regime parameters, we simulated the refolding timecourses of these two proteins in the absence of chaperones. As shown in Figs. 2D and 2E, the resulting timecourses accurately reproduce the experimentally observed refolding kinetics of RNase H^D^.^50^ Excellent agreement between simulations and experiments was also found for the concentration-dependent refolding behavior of fluc^40^ (Fig. 2I, 2J). These simulations also enabled following the misfolded-(M*) and aggregated-(A_20_) state timecourses under experimental fluc refolding conditions (Supplementary Fig. S1). Therefore, as an added advantage, these simulations also provide access to experimentally unassessed M* and A_20_ time-dependent concentrations.

The above results are important because they ensure that our kinetic modeling correctly captures the intrinsic folding and misfolding behavior of both clients in the absence of the Hsp70 chaperone system.

### Development of the CHAMPION70 kinetic model

Next, we extended the kinetic folding/aggregation framework developed in the previous section to include client-protein interactions with the Hsp70 chaperone system. The previously published CHAMP70 model^39^ (Supplementary Table S5-S6) describes only non-fold-promoting (“hold-only”) interactions and does not incorporate client-protein aggregation and chaperone interactions with misfolded intermediates (M*). To address these limitations, we integrated the client-protein folding and aggregation pathways described in the previous section into a new Hsp70 model that encompasses Hsp70 interactions with both unfolded and misfolded species.

We first implemented an early iteration of the program based solely upon a hold-only kinetic model that does not involve any active unfolding of bound clients. In this model, both unfolded (U) and misfolded (M*) client-proteins bind to the Hsp70 system and are released in the same conformation as the one in which they originally bind. Using the kinetic scheme in Supplementary Fig. S3 and the rate constants in Supplementary Table S9, the hold-only kinetic scheme closely reproduces RNase H^D^ refolding kinetics across multiple combinations of DnaK, DnaJ, and GrpE (Fig. 3A). When GrpE is added to complete the full K/J/E system, the model underestimates the experimental native-state formation rate. Crucially, however, it successfully reproduces the overarching trend: full K/J/E chaperone cycling slows native-state acquisition, with the model simply being slightly more effective at retarding folding than seen experimentally. These results confirm that RNase H^D^ engages the Hsp70 system through non-fold-promoting (hold-only) interactions.

**Fig. 3.**
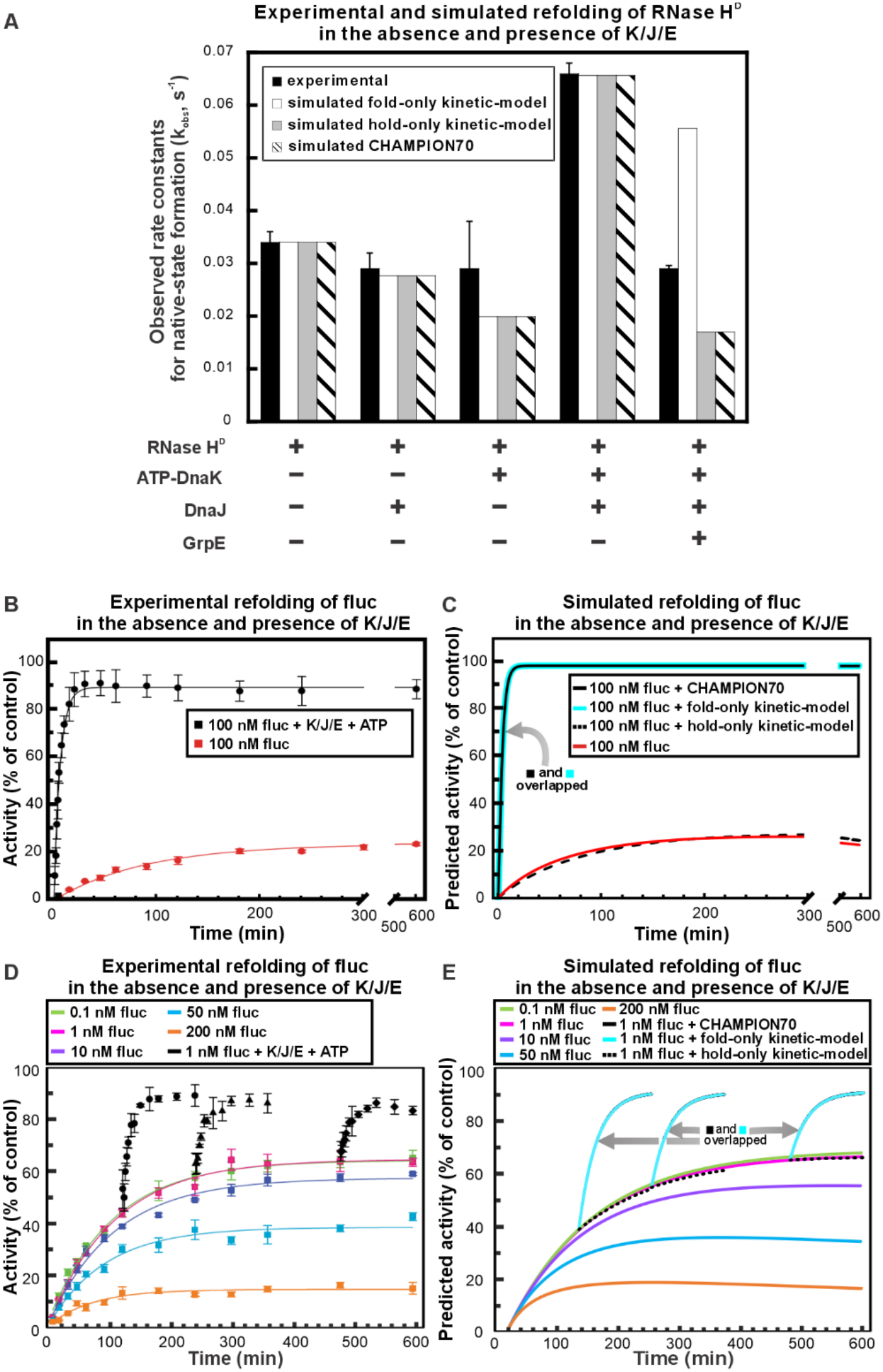
Experimental and simulated refolding of RNase H^D^ and firefly luciferase in the presence of Hsp70. **A.** Experimental RNase H^D^ refolding data obtained in the presence of different combinations of K/J/E chaperones were compared with simulations from the hold-only kinetic model, the fold-only kinetic model, and the unified CHAMPION70 model. All simulations used starting concentrations matching the experimental conditions: 5 μM of client-protein (U’),15 μM DnaK, 3 μM DnaJ, 6 μM GrpE, and 1 mM ATP (Sekhar *et al.*, 2012). **B.** Refolding of 100 nM fluc in the absence or presence of *E. coli* Hsp70 (3 μM DnaK, 1 μM DnaJ, 1.5μM GrpE, and 5mM ATP), reproduced from Imamoglu *et al.* (2020). **C.** Simulation of the data in panel B using four conditions: the no-chaperone folding model validated in Fig. 2, the hold-only kinetic model, the fold-only kinetic model, and the full CHAMPION70 model. All simulations use the same chaperone and nucleotide concentrations as in panel B. **D.** Concentration-dependent refolding activity of fluc (0.1-200 nM) in the absence and presence of chaperones (3 μM DnaK, 1 μM DnaJ, 1.5μM GrpE, and 5mM ATP), reproduced from Imamoglu *et al.* Black circles, triangles and diamonds indicate folding rescue of 1 nM fluc by addition of DnaK, DnaJ, GrpE and ATP. **E.** Simulation of the data in panel D using the same four conditions as in panel C: the non-chaperone folding model, the hold-only kinetic model, fold-only kinetic model, and the full CHAMPION70 model. All simulations use the same chaperone and nucleotide concentrations as in panel D.

Next, we applied the same hold-only framework to the strongly aggregation-prone protein (in the absence of chaperones) fluc. In this case, the hold-only (i.e., non-fold-promoting) model produced a pronounced mismatch with experimental observations (Fig. 3C, 3E). Given that misfolded fluc is released from Hsp70 in the same misfolded conformation, it remains highly susceptible to aggregation. As a result, simulated folding kinetics and native-state yields closely resemble those observed in the absence of chaperones and fail to capture the experimentally observed enhancement of fluc refolding by the Hsp70 system. The above findings demonstrate that a purely non-fold-promoting mechanism cannot account for Hsp70-mediated refolding of aggregation-prone clients.

Importantly, we also evaluated an unfold-only kinetic model on fluc refolding, where Hsp70 releases bound clients exclusively in the unfolded state (Supplementary Fig. S5 and S6). Although this unfold-only model increases the pool of unfolded fluc available for productive folding, the resulting kinetics remain similar to refolding in the absence of chaperones and fail to reproduce the experimentally observed enhancement of fluc refolding. Hence, merely unfolding a client protein is insufficient to explain Hsp70-mediated rescue of aggregation-prone proteins. This result is surprising and highly significant to understand Hsp70’s mechanism of action.

We then evaluated whether a purely fold-promoting interaction scheme could account for Hsp70-mediated refolding behavior. According to our fold-only kinetic model (Supplementary Fig. S4), unfolded (U) and misfolded (M*) client-proteins bind to the Hsp70 system and are released directly into the native-state (N). Use of this scheme for fluc refolding in the presence of the K/J/E chaperone system produced very good agreement between kinetic simulations and experimental refolding kinetics and native-state yields (Fig. 3C, 3E). The latter result shows that fold-promoting interactions are needed to rescue aggregation-prone clients. In contrast, however, the fold-only scheme did not reproduce RNase H^D^’s refolding behavior (Fig. 3A). Given that the fold-only kinetic model forces every chaperone cycling event to yield N, complete K/J/E cycling accelerates native-state acquisition. This acceleration directly contradicts experimental observations for RNase H^D^, where full K/J/E depresses the rate of native-state formation.^50^ Together, these combined RNase H^D^ and fluc validations demonstrate that neither a purely hold-only or fold-promoting kinetic model can universally describe all Hsp70 clients.

In order to capture both hold-only and fold-promoting behaviors within a single kinetic model, we combined the two interaction modes into the CHAMPION70 (**Cha**perone-**M**ediated **P**rotein folding kinetic Simulat**ion**) scheme, according to which unfolded (U) clients follow a non-fold-promoting pathway and misfolded (M*) clients follow a fold-promoting pathway (Fig. 4). The CHAMPION70 model is in excellent agreement with most of the experimental refolding data for both RNase H^D^ (Fig. 3A) and fluc (Fig. 3C, 3E), capturing the non-fold-promoting behavior of RNase H^D^ and the fold-promoting enhancement observed for fluc. Remarkably, this result was achieved with no need to introduce any client-specific chaperone-related kinetic parameters.

**Fig. 4.**
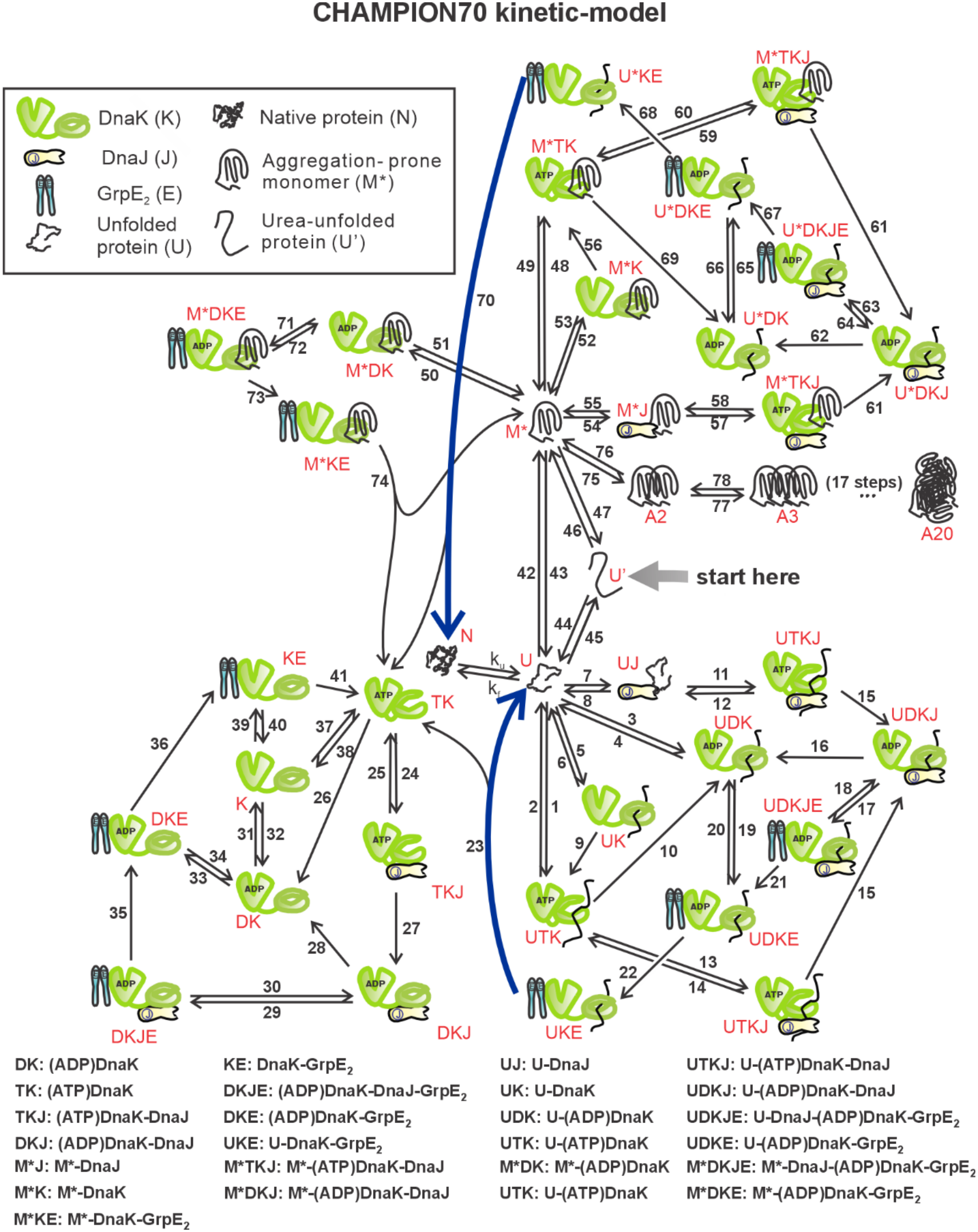
CHAMPION70 kinetic scheme. The CHAMPION70 kinetic model begins from the urea-unfolded state U’, which collapses into a conformationally relaxed unfolded state U or a misfolded state M* upon transfer into an excess of aqueous buffer within a hypothetical protein refolding process that either lacks or includes chaperones. U either folds to the native state N or interacts with Hsp70 (i.e., K/J/4E) via steps that involve no conformational changes. Conversely, misfolded species M* interacts with Hsp70 through steps that lead to an unfolded-like conformation denoted here as U*. The unfolded-like species U* is subsequently released as native state N. Alternatively, M* can also self-associate to generate progressively larger species that ultimately lead to the final aggregate, which is modeled here as a 20-mer denoted as A_20_. Independent nucleotides involved in some of the steps are not explicitly shown, for simplicity. See Supplementary Table S7 for a complete list of each of the steps, including all reactants and products as well as all the pertinent rate constants.

### Even clients with multiple binding sites rarely bind more than a single Hsp70 chaperone molecule at a time

Sequence-based predictors identify a single Hsp70 binding site in RNase H^D^ and multiple sites in fluc (Supplementary Fig. S9)^18^, raising the question of whether simultaneous binding by multiple chaperones onto a single client chain must be modeled. However, experimental proteome-wide DnaK (i.e., *E. coli* Hsp70) binding measurements show that steady-state client occupancy is exceptionally low across the *E. Coli* proteome.^54^ This occupancy is described by the relative enrichment factor denoted as REF. This parameter is defined as the fraction of client-protein population bound to DnaK at steady-state relative to the total client-protein population in the cell lysate.^54^ Even for client-proteins displaying the highest measured binding levels in the dataset, the maximal observed REF value is ∼0.05. This result shows that over 95% of the cellular protein population is completely unbound, at any given time.^54^

Probabilistic modeling of independent site occupancy suggests that over 97% of chaperone-bound client molecules carry only a single DnaK chaperone at a time. This outcome renders simultaneous multi-chaperone engagement on an individual client chain, unlikely under physiological conditions (see Supplementary Materials for a full statistical derivation and additional considerations).

While simplified simulations under non-aggregating conditions show that additional chaperone binding sites delay early folding without altering final native yields (Supplementary Fig. S10D), CHAMPION70 robustly reproduces the experimental refolding behavior of fluc using a single binding site scheme (Fig. 3C, 3E). Consequently, parameterizing CHAMPION70 with a single client binding site is sufficient to capture the predominant kinetic features of the Hsp70 chaperone system while, at the same time, avoiding unnecessary model over-parameterization.

### Hsp70-chaperone-mediated protection from aggregation only applies over finite timescales and eventually fails

After validating CHAMPION70 against experimental client-refolding data, we employed this model to predict how client-protein’s intrinsic folding rates, thermodynamic stability and aggregation propensities shape interactions with Hsp70 as well as aggregation-*vs*-folding outcomes. Towards this end, we employed four representative client proteins belonging to the kinetic and thermodynamic regimes defined by regions I through IV in Fig. 5A. These representative clients span the slow-to-fast folding-kinetics regimes and range from being thermodynamically unstable to thermodynamically stable (Fig. 5A; Supplementary Table S11). This client-protein classification is consistent with a previous study where Hsp70 action was formally analyzed in the absence of client-protein aggregation.^55^ Each of the 4 representative client proteins contains a single predicted DnaK-binding site.^39^ DnaK interactions have also been experimentally observed^37,50,56^ except for the formin binding protein (FBP), which represents fast-folding and thermodynamically unstable proteins. These features enable us to assess the effect of Hsp70 on client-protein folding and aggregation in the absence of complicating factors arising from the presence of multiple binding sites.

**Fig. 5.**
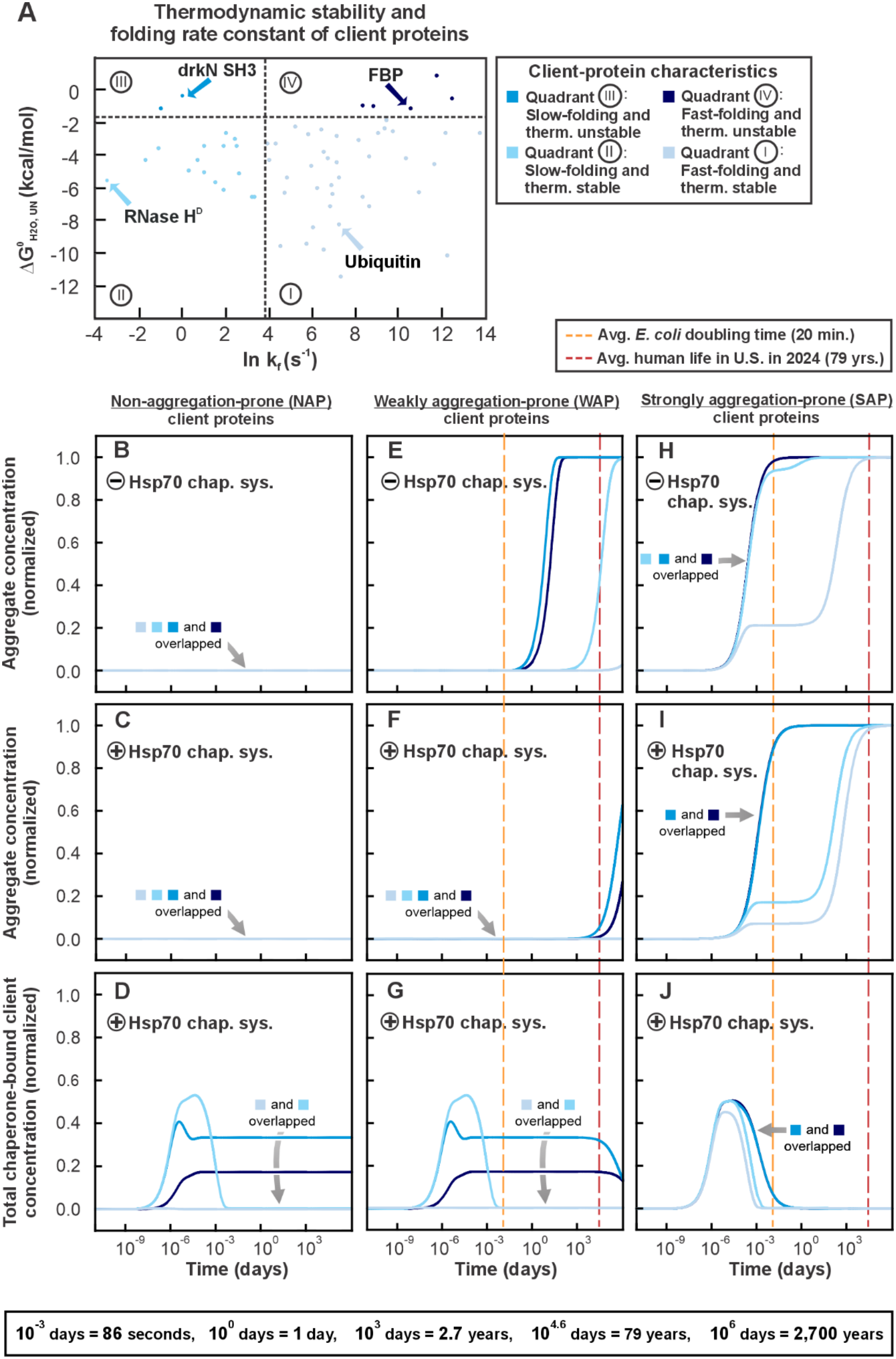
Hsp70 cannot indefinitely prevent client-protein aggregation. **A.** Overview of the folding rate constants and thermodynamic stabilities of 61 client proteins from KineticDB (Bogatyreva *et al.*, 2009). Four proteins were selected (blue, various shades) as representative client proteins from each quadrant. All simulations began with 15 μM client-protein in the U’ state. Simulations in the presence of chaperones contained 25 μM DnaK, 5 μM DnaJ, 10 μM GrpE, 3.5 mM ATP, 0.12 mM ADP, and 5 mM P_i_. **B, E, H.** Simulated formation of the fully aggregated A_20_ species in the absence of chaperones for clients with three aggregation propensities: non-aggregation-prone (NAP) (**B**), weakly aggregation-prone (WAP) (**E**), and strongly aggregation-prone (SAP) (**H**), shown for each of the four representative folding profiles. **C, F, I.** Simulated formation of A_20_ in the presence of chaperones for non-aggregation-prone (NAP) (**C**), weakly aggregation-prone (WAP) (**F**), and strongly aggregation-prone (SAP) (**I**) client-proteins. **D, G, J.** Simulated association of client species with the Hsp70 cycle for non-aggregation-prone (NAP) (**D**), weakly aggregation-prone (WAP) (**G**), and strongly aggregation-prone (SAP) (**J**) client-proteins.

First, when client-protein aggregation is absent (NAP conditions), we find that thermodynamically stable clients fold too rapidly to significantly interact with the chaperone. This concept is quantitatively illustrated by the CHAMPION70 simulation results of Fig. 5D.

In contrast, thermodynamically unstable client proteins are predicted to remain persistently bound to Hsp70 regardless of their folding rates (Fig. 5D). Thus, client-protein thermodynamic instability is expected to impose a heavy chaperone burden on the cell machinery. This prediction has important evolutionary implications. For instance, intrinsically disordered proteins (IDPs), which are far more abundant in eukaryotes than in bacteria,^6^ belong to this category of thermodynamically unstable proteins. Importantly, IDPs minimally interact with Hsp70 and instead often partition into biomolecular condensates.^57,58^ Proteome-wide analyses show that intrinsic disorder negatively correlates with chaperone binding, as IDPs’ low hydrophobicity leads to them carrying fewer nonpolar Hsp70-binding motifs than foldable proteins.^59^ On the other hand, IDPs still carry a number of Hsp70 binding sites.^59^ Now, it is possible that IDPs evolved a tendency to phase-separate as condensates to escape the unstable proteins’ thermodynamic tendency to persistently bind Hsp70, consistent with Fig. 5D. In this way, IDPs, which are particularly abundant in eukaryotes but are also found in prokaryotes, do not monopolize chaperone capacity, and preserve Hsp70’s availability to assist folding-prone (i.e., non-IDP) client proteins, which benefits the most from chaperone engagement.

When aggregation propensity is weak (WAP conditions), Hsp70 is hugely beneficial and maintains client-protein solubility for decades, substantially extending the time before aggregation takes over (Fig. 5E, F). Fast-folding and thermodynamically stable proteins (class I) fold efficiently on their own and therefore do not engage with Hsp70. Conversely, slow-folding and stable proteins (class II) transiently and effectively bind Hsp70, resulting in the longest persistence in soluble-client lifetime (Fig. 5F, G). After ∼4-6 years, unstable proteins (classes III and IV) begin to aggregate, though they remain largely soluble over decade-long timescales. Just as it was observed under NAP conditions, these thermodynamically unstable proteins bind Hsp70 for the entirety of the simulation (i.e., ∼3,170 yrs, Fig. 5G), with a modest decline in binding coinciding with the onset of aggregation. The above results show that, under modest client-protein aggregation propensity (WAP conditions), Hsp70 supports client solubility for extended periods of time (>79 yrs). This protection timescale clearly exceeds the lifespan of many organisms, including *E. coli*, which can therefore experience myriads of aggregation-free doubling events (each occurring every ca. 20 min) in the presence of Hsp70.

When client-protein aggregation propensity is strong (SAP conditions), not surprisingly, all types of clients are predicted to undergo aggregation at a faster rate. Within this regime, the M* basin is lowered, placing M* close in standard-state free energy to the native-state (Supplementary Fig. S12). This energetic proximity suggests that M* adopts a compact, partially native-like conformation, consistent with experimental φ-value studies, which show that misfolded intermediates and their transition states can retain substantial native-like structure.^60^ In the case of the PDZ-domain protein, for instance, the misfolded intermediate retains substantial native structure, reducing solvent exposure of nonpolar segments that are typically recognized by Hsp70.^60^ These compact misfolded states are therefore poor Hsp70 clients, and provide a structural explanation for the increased M* population and accelerated aggregation observed for the clients falling under the SAP regime (Fig. 5H, I, J). Importantly, the simulations demonstrate that Hsp70 provides only a finite protection from client-protein aggregation. Thermodynamically stable proteins experience substantial aggregation delays in the presence of Hsp70, with the slow-folding stable client gaining many decades of additional solubility (Fig. 5H, I). Unstable proteins, however, aggregate rapidly despite prolonged chaperone binding.

In summary, the CHAMPION70 simulations show that SAP clients are not optimally served by the Hsp70 chaperone system, and aggregation eventually overcomes chaperone-mediated rescue. At its core, this phenomenon is related to the intrinsic higher thermodynamic stability of aggregates relative to folded states and complexes with the Hsp70 chaperone system. As a result, long-term Hsp70 association is replaced by aggregate formation (Fig. 5I, J), and the chaperone system is therefore ultimately unescapably destined to fail. Intriguingly, all SAP clients are predicted to reach complete aggregation within ca. 79 years, which is the average human lifespan in the U.S. in 2024, released by the US National Center for Health Statistics in 2026.^61^ While this multi-decade timescale far exceeds *E. Coli* doubling time, it highlights the intrinsic limits of chaperone capacity under severe aggregation pressure, independent of post-translational modification of clients or other proteostasis-related events.

In all, the above outcome demonstrates that ultimate long-term thermodynamically driven sinking into aggregated states imposes a fundamental boundary on proteostasis. This constraint applies even in the absence of aging-related decreases in chaperone concentration, which are often invoked to explain aging-related proteostasis failures. Thus, the computational predictions shown in this work carry important implications for the health of *E. coli’s* colonies over time. In addition, the general concept of long-term thermodynamically driven chaperone failure also has key implications for long-lived eukaryotic organisms, including humans.

### The presence of secondary nucleation renders Hsp70 less effective at delaying aggregation

Incorporation of monomer-dependent secondary nucleation into the simulations further underscores the kinetic limits discussed in the previous section (Supplementary Fig. S16). Up to now, simulations modeled aggregation exclusively via rate-limiting primary nucleation, requiring two misfolded monomers (M*) to collide and form a dimeric intermediate prior to elongation. In contrast, the presence of secondary nucleation bypasses the rate-limiting primary nucleation step and accelerates aggregate buildup by catalyzing new nucleation events on existing aggregates’ surfaces.^62^ Under WAP conditions, thermodynamically stable proteins are predicted to remain resistant to aggregation in the presence of Hsp70. On the other hand, unstable clients are projected to exhibit moderately increased aggregation (Supplementary Fig. S16J).

The most dramatic effects are observed under SAP conditions. In the presence of Hsp70, the presence of monomer-dependent secondary nucleation compresses the timescale for complete aggregation from ca. 79 years to ca. 3 years (Supplementary Fig. S16M, S16N). Thus, Hsp70’s ability to prevent inevitable long-term aggregation is expected to be further challenged in the presence of secondary nucleation.

### Hsp70 sequestration of thermodynamically unstable proteins reduces native-state acquisition

Native-and misfolded-state timecourses, shown in Fig. 6, clarify why thermodynamically unstable proteins are expected to impose such a persistent burden on the cell machinery. Under NAP conditions, thermodynamically stable clients reach full native occupancy both in the absence and presence of Hsp70 (Fig. 6B, C), while thermodynamically unstable clients retain substantial unfolded populations. Hsp70 binding further shifts unstable clients toward unfolded states, reducing their native-state population, a direct consequence of their extended association time with the chaperone.

**Fig. 6.**
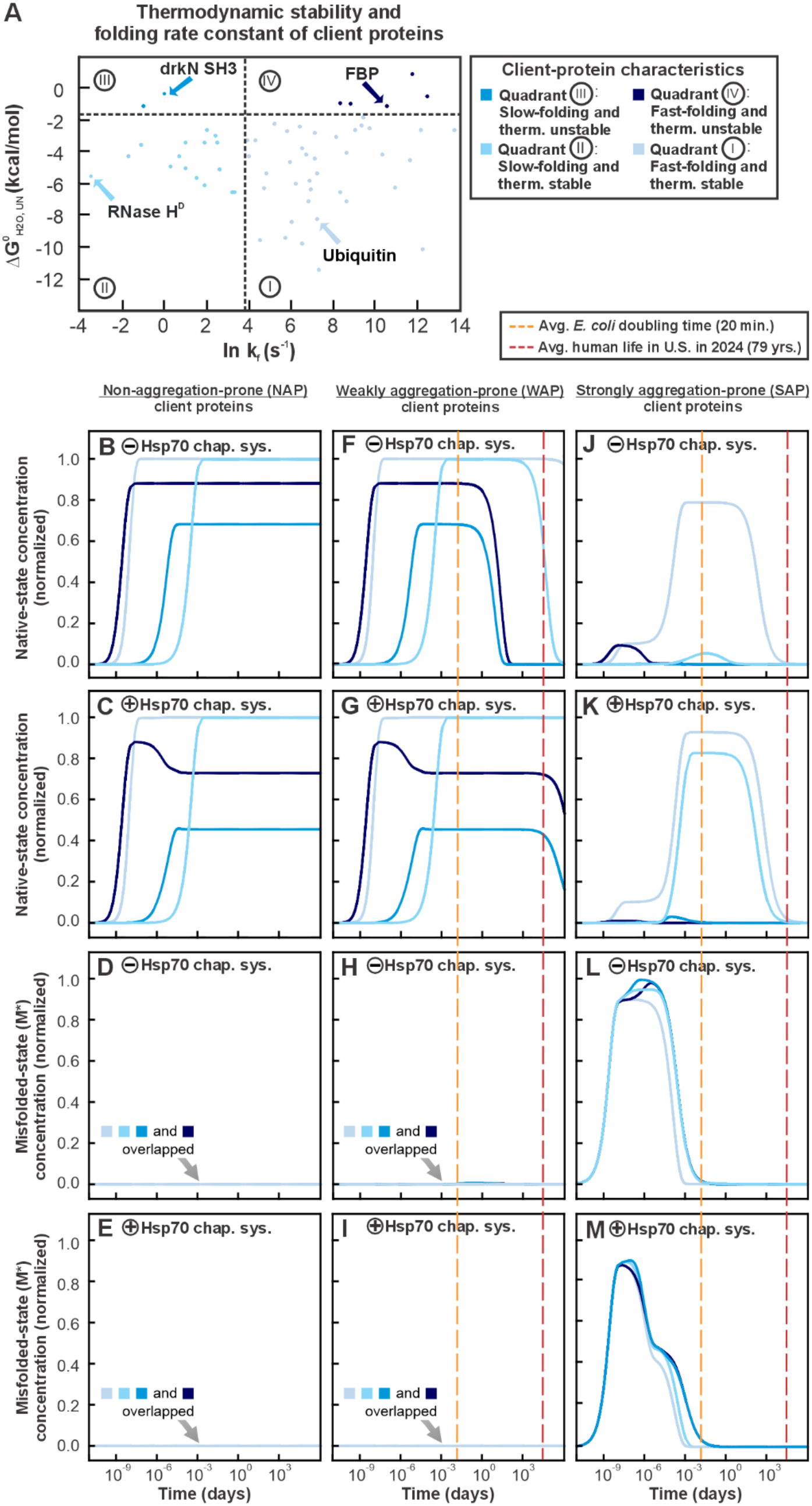
Hsp70 binding of thermodynamically unstable clients reduces native-state acquisition. All simulations start from 15 μM of client-protein in the U’ state. Simulations in the presence of chaperones include 25 μM DnaK, 5 μM DnaJ, 10 μM GrpE, 3.5 mM ATP, 0.12 mM ADP, and 5 mM P_i_. **A.** Overview of the folding rate constants and thermodynamic stabilities of 61 client proteins from KineticDB (Bogatyreva *et al.*, 2009). **B, F, J.** Simulated formation of the native-state in the absence of chaperones for clients with three aggregation propensities: non-aggregation-prone (NAP) (**B**), weakly aggregation-prone (WAP) (**F**), and strongly aggregation-prone (SAP) (**J**), shown for each of the four representative folding profiles. **C, G, K.** Simulated formation of the native-state in the presence of chaperones for non-aggregation-prone (NAP) (**C**), weakly aggregation-prone (WAP) (**G**), and strongly aggregation-prone (SAP) (**K**) client-proteins. **D, H, L.** Simulated formation of the misfolded state in the absence of chaperones for non-aggregation-prone (NAP) (**D**), weakly aggregation-prone (WAP) (**H**), and strongly aggregation-prone (SAP) (**L**) client-proteins. **E, I, M.** Simulated formation of the misfolded state in the presence of chaperones for non-aggregation-prone (NAP) (**E**), weakly aggregation-prone (WAP) (**I**), and strongly aggregation-prone (SAP) (**M**) client-proteins.

Under weak aggregation propensities (WAP conditions), stable proteins again reach fully native populations (Fig. 6F, G), while unstable proteins maintain a balance between native and unfolded state (Fig. 6F, G). The M* populations remain low (Fig. 6H, I), though even small, persistent M* populations are ultimately expected to funnel proteins to aggregation, on long timescales. In essence, chaperone action delays accumulation of M* and slows down aggregation by partitioning unfolded clients away from misfolded states. This scenario is consistent with the multi-year delays in aggregation shown in Fig. 5F and 5G.

Under strong aggregation propensity (SAP conditions), native-state formation is initially enhanced for stable proteins in the presence of Hsp70 (Fig. 6K), but this recovery is only temporary. As M* accumulates rapidly (Fig. 6L, M), native-state populations decline for all clients. Unstable proteins fail to sustain any native structure despite extensive chaperone engagement, given that Hsp70 repeatedly returns them to unfolded states that readily repopulate M*.

Together, the above data show that unstable proteins both monopolize Hsp70 currency and yet remain poorly folding clients, explaining why they aggregate rapidly even when chaperones are abundant, while stable proteins can be partially, though not indefinitely, rescued.

### Higher-than-physiological Hsp70 concentrations are unable to prevent aggregation

Next, we evaluated whether increased chaperone availability may overcome the unavoidable long-term aggregation observed under WAP and SAP conditions. Towards this end, we simulated the four representative client-proteins across a range of total Hsp70 concentrations spanning lower-than physiological, physiologically relevant, and higher-than-physiological-chaperone levels. The results are shown in Fig. 7.

**Fig. 7.**
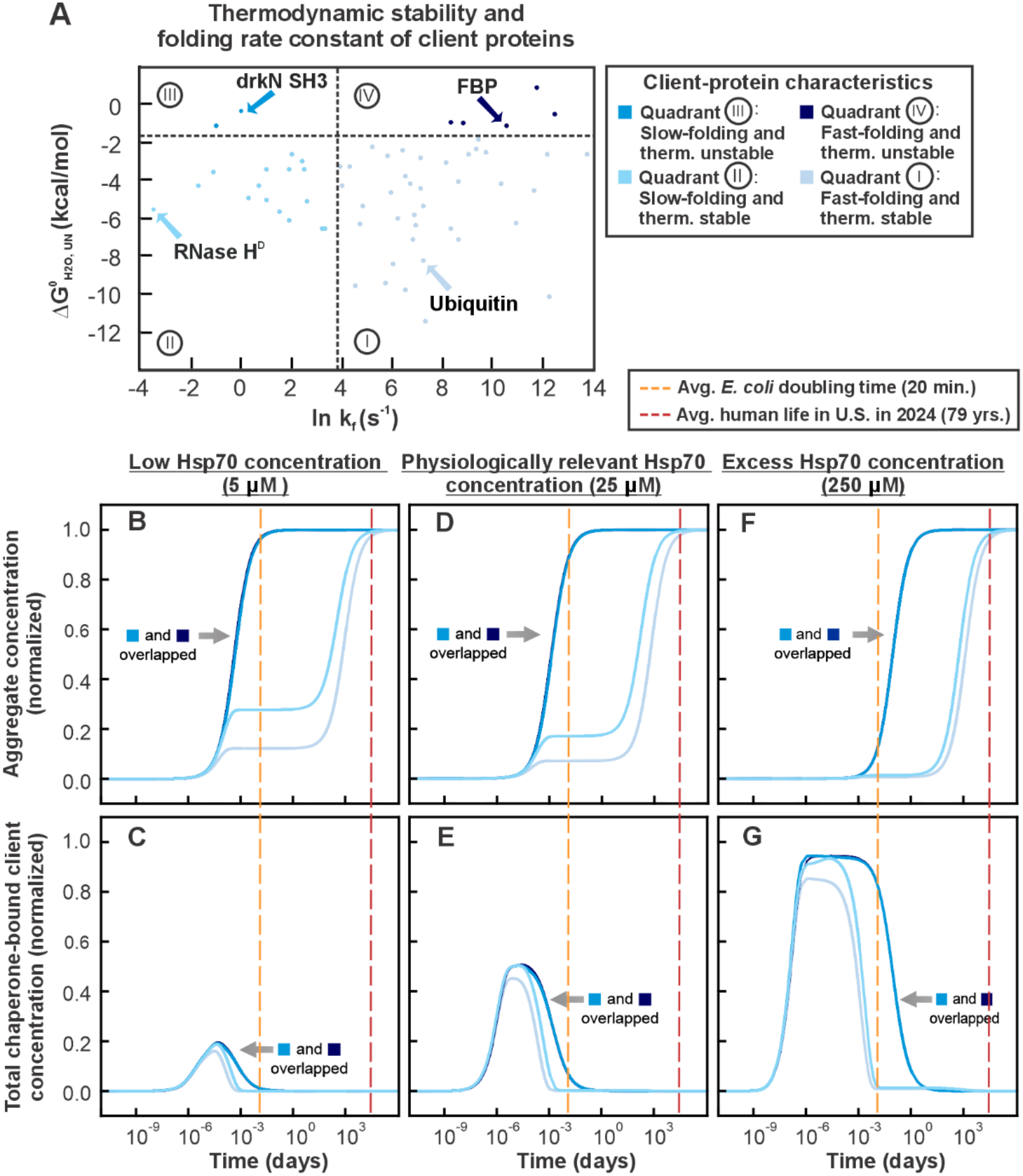
Elevated chaperone concentrations increase client-chaperone association without preventing long-term aggregation. All simulations start from 15 μM client-protein in the U’ state and contained 5 μM DnaJ, 10 μM GrpE, 3.5 mM ATP, 0.12 mM ADP, and 5 mM P_i_. The concentration of DnaK was varied across simulations (5 μM, 25 μM, and 250 μM). **A.** Overview of the folding rate constants and thermodynamic stabilities of 61 client proteins from KineticDB (Bogatyreva *et al.*, 2009). **B, D, F.** Simulated formation of the fully aggregated A_20_ species under strongly aggregation-prone (SAP) conditions at 5 μM DnaK (**B**), 25 μM DnaK (**D**), and 250 μM DnaK (**F**). **C, E, G.** Simulated total chaperone-bound client concentration under the same strongly aggregation-prone (SAP) conditions at 5 μM DnaK (**C**), 25 μM DnaK (**E**), and 250 μM DnaK (**G**).

Higher Hsp70 levels promote more rapid client association with Hsp70 and prolongs the duration of chaperone engagement, resulting in sustained interaction throughout the early stages of the simulations (Fig. 7G). Experimental studies show that Hsp70 exists in a monomer-oligomer equilibrium even under basal conditions, and that higher temperatures, ATP depletion, and increased Hsp70 concentration all shift more of the chaperone into oligomers that do not bind client-proteins.^63,64^ Given that our simulations assume that all Hsp70 remains available for client binding, they likely overestimate the effective chaperone pool *in vivo*. Despite the above effects, elevated Hsp70 concentrations do not prevent long-term aggregation. Thermodynamically unstable proteins aggregate rapidly across all Hsp70 levels. Thermodynamically stable proteins exhibit overall slower aggregation. Increasing Hsp70, however, primarily decreases the extent of intermediate aggregation rather than altering the timing of progression to full aggregation. The onset of aggregation occurs approximately at the same time, across all Hsp70 concentrations. This results shows that enhanced chaperone availability reduces the extent of early misfolding but does not shift the ultimate aggregation fate of the protein population.

In short, the simulations of Figure 7 demonstrate that, while increasing Hsp70 capacity can delay aggregation and transiently stabilize vulnerable clients, it cannot eliminate the intrinsic tendency of proteins to misfold and aggregate over long timescales.

### The Hsp70 chaperone is far from perfect

Chaperones provide an essential service to cellular life by protecting client-proteins that are vulnerable to misfolding and aggregation, starting from the early stages of protein life.^10,65–67^ Our simulations show that, while the Hsp70 system delays aggregation and promotes refolding for these clients, its protective capacity is limited. Fig. 8 summarizes these limitations (Fig. 8A-8L). Over short timescales, Hsp70 effectively stabilizes unfolded clients and promotes native-state formation, proving to be particularly effective in the case of thermodynamically stable proteins. Over long timescales, however, these early advantages get progressively attenuated. Misfolded and aggregated species accumulate, and the chaperone system becomes increasingly unable to rescue unstable clients or prevent the eventual loss of native structure. This behavior reflects a fundamental inherent thermodynamic constraint of protein landscapes. Although the native state is a local, metastable minimum, the aggregated state represents a deeper and more stable free-energy basin. As a result, Hsp70’s ability to suppress aggregation is inherently time-limited, even under ideal conditions. Over sufficiently long timescales, even thermodynamically stable proteins eventually aggregate despite the presence of fully bioactive molecular chaperones in the medium. This constraint is expected to be particularly problematic for long-lived organisms.

**Fig. 8.**
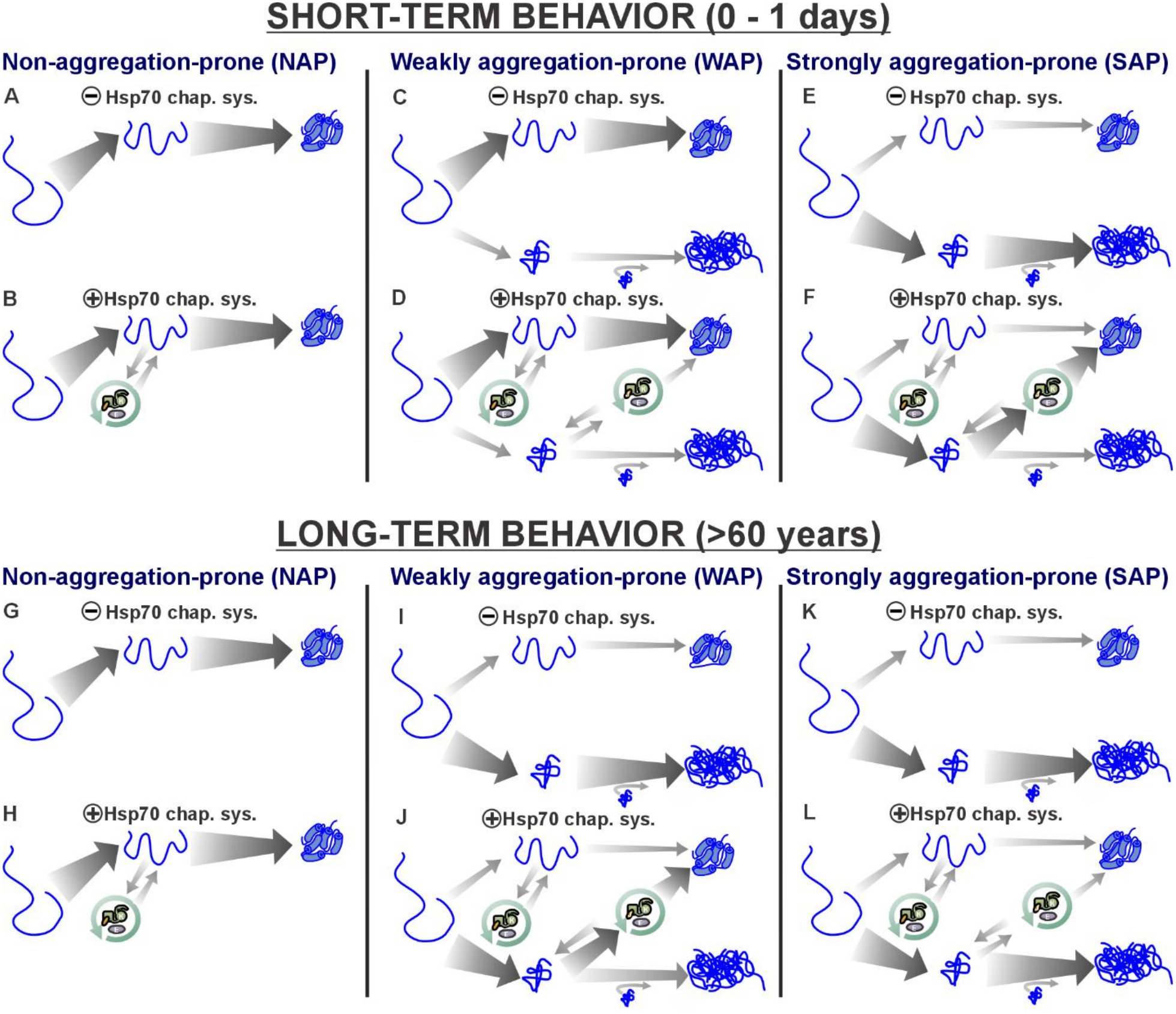
Overview of the influence of the Hsp70 chaperone system on client-protein folding and aggregation. Overview of the dominant folding outcomes for representative client-proteins with three aggregation propensities: non-aggregation-prone, weakly aggregation-prone, and strongly aggregation-prone. Each panel illustrates the prevailing fate of the client population under conditions without or with the Hsp70 chaperone system, evaluated at short-term and long-term time points. **A, C, E.** Short-term outcomes without chaperones for non-aggregation-prone (NAP) (**A**), weakly aggregation-prone (WAP) (**C**), and strongly aggregation-prone (SAP) (**E**) clients. Non-and weakly aggregation-prone (NAP and SAP) proteins favor the native-state while strongly aggregation-prone (SAP) proteins tend toward aggregation. **B, D, F.** Short-term outcomes with chaperones for non-aggregation-prone (NAP) (**B**), weakly aggregation-prone (WAP) (**D**), and strongly aggregation-prone (SAP) (**F**) clients. Chaperones shift all three classes toward folding. **G, I, K.** Long-term outcomes without chaperones for non-aggregation-prone (NAP) (**G**), weakly aggregation-prone (WAP) (**I**), and strongly aggregation-prone (SAP) (**K**) clients. Weakly and strongly aggregation-prone (WAP and SAP) clients accumulate in the fully aggregated A_20_ state. **H, J, L.** Long-term outcomes with chaperones for non-aggregation-prone (NAP) (**H**), weakly aggregation-prone (WAP) (**J**), and strongly aggregation-prone (SAP) (**L**) clients. Weakly aggregation-prone (WAP) clients show substantial native recovery but still form some aggregated species. Strongly aggregation-prone (SAP) clients exhibit limited rescue by chaperones and predominantly accumulate in the fully aggregated state.

Our simulations also show that increasing the concentration of the Hsp70 chaperone system can further delay late aggregation but cannot prevent it indefinitely. This result highlights a broader limitation of hold-only and fold-promoting chaperone mechanisms. Chaperones like Hsp90,^68,69^ which primarily stabilizes and remodels late-stage folding intermediates, and GroEL/GroES,^69,70^ which promotes productive folding within a protected cavity, also rely on cycles of binding, stabilization, and refolding. These mechanisms can transiently improve solubility either by holding misfolded species or by accelerating folding. However, they do not eliminate misfolded or aggregation-competent clients. As a result, even specialized hold-only or fold-promoting chaperones ultimately face the same long-term limitation of Hsp70.

Cells, therefore, must rely on additional protein quality-control mechanisms that address the intrinsic limitations of the chaperone machinery as a whole. Misfolded and aggregated proteins can be removed through dedicated degradation pathways, including proteasomal degradation in eukaryotes and Lon/Clp proteases in bacteria.^71,72^ Eukaryotic cells can also eliminate larger aggregates through selective autophagy, delivering aggregated proteins to the lysosome for breakdown. In addition to these degradation routes, many quality-control systems promote disaggregation,^73–75^ resolubilizing aggregates so proteins can either refold into a functional state or be directed to degradation machinery. The above complementary pathways collectively display protective capabilities that Hsp70 alone, or any chaperone limited to hold-only and/or fold-promoting behavior, cannot sustain. Further studies are needed to explore whether and how these activities integrate with the Hsp70 hold-only and fold-promoting behaviors to maintain long-term proteostasis with no excessive free-energy penalties.

### Implications for prokaryotic and eukaryotic organisms

To date, most experimental kinetic studies have been performed with the *E. Coli* Hsp70 system (DnaK, DnaJ, GrpE). Although CHAMPION70 employs rate constants that have been experimentally assessed for the bacterial Hsp70 chaperone system, two important considerations are in hand. First, the Hsp70 ATPase cycle is highly conserved across a wide variety of organisms.^13–15^ Second, experimental measurements on the human stress-inducible Hsp70A1 chaperone show that ATP binding, ATP hydrolysis, and ATP-induced conformational transitions occur with rate constants of the same order of magnitude as those used in CHAMPION70 (Supplementary Table S16).^76^ Therefore, the above similarities suggest that the kinetic regime explored here may be broadly relevant to both bacterial and human Hsp70. Therefore, while the CHAMPION70 kinetic model developed here is grounded in *E. coli* biochemistry, its mechanistic predictions may well capture features of Hsp70 behavior that extend across different organisms.

*E. coli* has a cell-division time of ∼20 min, and proteins subject to weak aggregation conditions (WAP) are not at risk of aggregating, up to individual cell-doubling timescales. In fact, even in the absence of Hsp70, most WAP proteins remain soluble over a single cell cycle, and in the presence of Hsp70 even longer-term aggregation is effectively prevented (Fig. 5E, F). Under strong aggregation-prone (SAP) conditions, however, the scenario is more nuanced. Hsp70 is essential to protect slow-folding thermodynamically stable proteins across bacterial single-doubling timescales. On the other hand, thermodynamically unstable proteins aggregate rapidly even in the presence of the chaperone (Fig. 5H, I). Interestingly, over timescales corresponding to the end of typical stationary phases under physiologically relevant conditions (ca. 2 weeks)^77^, the Hsp70 chaperone system is predicted to start failing at its aggregation-protection capability (Fig. 5H, I). Remarkably, the latter timescale (ca. 14 days) corresponds to the known experimental onset of the bacterial cell death phase.^77^ Hence, this prediction suggests that *E. coli*’s death may be significantly contributed by the intrinsic failure of the Hsp70 chaperone system to protect all classes of proteins beyond ∼2 weeks.

Note that, even if cell division in bacteria is known to dilute aggregates from WAP and SAP proteins via asymmetric partitioning upon cell doubling,^78^ new aggregation events reinstate the aggregated status within seconds. Therefore, WAP and SAP proteins, which are known to exist in bacteria (Dang *et al.*, in preparation; and ^79^ and ^6^) pose a real threat to long-term bacterial doublings. Bacteria manage this threat via a relatively simple proteostasis network that relies on chaperones including Hsp70 (e.g., in the cytoplasm, mainly Trigger Factor, the K/J/E Hsp70 system, the GroEL/GroES Hsp60 system, the ClpB Hsp100 disaggregase, and a small number of small heat-shock proteins a.k.a. sHsps). Hsp70, together with the other chaperones, applies sufficient aggregation protection across typical bacterial growth and stationary phases of ca. 2 weeks.^80–82^ In summary, bacteria are subject to an inescapable trend according to which the Hsp70 chaperone system eventually fails to prevent weak and strong aggregation trends beyond a few weeks. Remarkably, this trend does not require any classical proteostasis-imbalance events (e.g., chaperone downregulation), which are typically associated with multiple bacterial doublings and bacterial senescence.^83^

Juxtaposed to the case of bacteria, eukaryotic cells, especially those belonging to long-lived multicellular organisms, face even more severe challenges. Proteins must remain soluble for years or decades, and even low levels of misfolded species can progressively turn into soluble and insoluble aggregates over extended timescales. This burden is especially severe in terminally differentiated, post-mitotic cells (e.g. cortical neurons and cardiomyocytes) which do not divide, requiring individual cells to maintain proteostasis across the organism’s entire lifespan.^84^ Moreover, eukaryotic proteomes contain a much higher proportion of aggregation-prone proteins than bacterial proteomes,^6^ further increasing the burden on Hsp70 chaperones. As a result, eukaryotes rely on an expanded collection of chaperones, including specialized cochaperones, ATP-independent and-dependent disaggregases, ubiquitin-proteasome and autophagy pathways, which buffer misfolding and aggregation over the extended eukaryotic lifespans.^6,85^ Despite this complex protective network, it is remarkable that the Hsp70 chaperone system is predicted to intrinsically fail at protecting both WAP and SAP proteins against aggregation over a timescale comparable to the average human lifetime in the U.S. (∼79 yrs; Fig. 5E,H,F,I). This failure represents a fundamental thermodynamic boundary that occurs even in the absence of any transcriptionally regulated proteostasis imbalances leading to decreases in Hsp70 concentration.

## Conclusions

In this work, we presented CHAMPION70, a novel kinetic model that integrates protein folding, misfolding, aggregation, and Hsp70 chaperone interactions within a single reaction network. By varying only client-specific folding and unfolding rate constants, the model captures both hold-only and fold-promoting chaperone behaviors, demonstrating that differences in Hsp70-mediated folding outcomes arise from intrinsic client-protein properties, particularly thermodynamic stability and folding rate.

By explicitly incorporating aggregation as a competing thermodynamic endpoint, the model reveals an inherent limitation of the Hsp70 system: it can delay, but not ultimately prevent, the accumulation of aggregated species. This limitation persists even under idealized conditions of constant chaperone availability, indicating that long-term aggregation is not solely a consequence of external stress, molecular damage, or age-related proteostasis decline.^86–88^ Instead, it reflects fundamental features of protein folding landscapes that chaperones can counteract only for finite periods of time.

## Methods

### General aspects of computer simulations

All computational modeling was performed with the COPASI (version4.44, build 295) software package.^53^ COPASI integrates ordinary differential equations using Livermore Solver of Ordinary Differential Equations Algorithm (LSODA).^89^ Rate laws for all elementary steps were implemented using COPASI’s standard procedures. COPASI assumes that reaction order matches molecularity for each elementary reaction.

All symbols, rate constants, and kinetic parameters are provided in Supplementary Tables S1 − S10. Rate constants for the Hsp70 chaperone-cycle steps, both in the absence and presence of client-protein, were selected to match experimentally determined values whenever possible, similarly to previously published simulations.^39^

Unless otherwise stated, simulations involving chaperones are preequilibrated *in silico* for 300 seconds in the absence of client-protein. The resulting concentrations of free chaperones, cochaperones, and their complexes are then used as initial conditions for simulations including client-protein. For simulations mimicking experimental time courses, total ATP is set to the experimentally used concentration before pre-equilibration. The resulting ATP, ADP and P_i_ concentrations after pre-equilibration are used as initial values for the client-containing simulations. In all other simulations, ATP, ADP, and P_i_ concentration are held constant after pre-equilibration to prevent depletion of the nucleotide pool.

Unless otherwise stated, simulations containing chaperones use 25 μM DnaK, 5 μM DnaJ, and 10 μM GrpE, concentrations that fall within physiologically relevant ranges for *E. Coli*.

### Protein folding simulations in the absence of chaperones

Supplementary Table S2. lists the 23 reversible reactions and associated rate constants used to model protein folding and aggregation in the absence of chaperones. These parameters were used for RNase H^D^, firefly luciferase (fluc), and all other client-proteins examined in this work. Folding rate constants for each protein are provided in Supplementary Table S3.

Aggregation is modeled with a reversible nucleation-elongation scheme according to which two aggregation-prone misfolded species (M*) form a dimer (A_2_) via a rate-determining nucleation step. Aggregate elongation follows via 18 equivalent reversible steps eventually leading to the final aggregate (A_20_). Aggregation elongation steps are steeply downhill (Supplementary Table S2). Supplementary Table S4 summarizes the kinetic modifications that are employed to generate NAP, WAP, and SAP conditions. These modifications alter the forward and reverse U’ to M* rate constants through changes in transition-state barrier, with no alteration in standard-state free-energies of U’ or M*.

Unless otherwise stated, simulations begin from a 15 μM client-protein in the U’ state, which represents a hypothesized denaturant-unfolded species immediately after dilution into refolding buffer. From U’, client-proteins proceed along two parallel routes: a folding pathway (via the conformationally relaxed unfolded state U) and an aggregation pathway (via M*).

### Updates to the CHAMP70 kinetic model

The original CHAMP70 model was implemented in Gepasi,^90^ which is no longer supported. COPASI reproduces Gepasi’s behavior but uses a different approach to identify conserved moieties (Householder reflections^53,91,92^ rather than moiety-conserved cycles^93,94^). The original 15 reversible and 13 irreversible steps and rate constants of the CHAMP70 kinetic model are listed in Supplementary Table S5.

CHAMP70 was updated to ensure that the system reflects a biologically relevant steady state. This was done by adjusting ATP, ADP, and P_i_ at experimentally determined concentrations within an *in vivo* environment (see Supplementary Materials for additional details).^95^ In addition, rate constants corresponding to ATP hydrolysis in the presence of DnaJ, in both the absence and presence of a client-protein (reaction steps 15, 27), were modified to better reproduce experimental ATP hydrolysis data.^96^

### CHAMPION70 kinetic model

The CHAMPION70 model expands upon CHAMP70 by including 44 reversible and 24 irreversible steps (Supplementary Table S7) so that both folding/unfolding and aggregation of client-protein are taken into account. Supplementary Table S4 summarizes the kinetic variations used to generate NAP, WAP and SAP conditions. For convenience, the differences between CHAMPION70, hold-only, fold-only, and unfold-only kinetic models are listed in Supplementary Table S9.

### Kinetic and thermodynamic parameters for client-protein folding

Thermodynamic stabilities and folding rate constants of the 61 small, two-state, single-domain proteins employed in this work were obtained from the KineticDB database (Supplementary Table S11).^55^ Four representative proteins were selected from each of the four quadrants based upon thermodynamic stabilities and folding rates (Fig. 5A) to provide representative overviews of client-protein behavior as a function of the client’s kinetic and thermodynamic characteristics.

## Acknowledgments

We thank Matt Dalphin and Natalie Feider for helpful discussions and technical assistance. We are also grateful to Devin Schultz and Gordon L.W. Winkler for their contributions to the early stages of the computational analysis. M.F.M. was the recipient of a Molecular Biophysics T32 Training Fellowship by the National Institutes of Health (5T32GM008293-24). This work was funded by the National Science Foundation (NSF) grants MCB 2124672 and MCB-2547879 to S.C.

## Availability of data and materials

Simulation data and/or CHAMPION70 program used in the current study available from the corresponding author upon reasonable request.

## Author contributions

M.F.M., M.J.D., and S.C. designed the research; M.F.M., M.J.D., J.K.M., H.Y., and X.C. performed the computations; M.F.M., M.J.D., S.C.C. and S.C. prepared the figures and wrote the manuscript.

