## Supplementary Materials for "Unavoidable Disasters: the Hsp70 Chaperone Can Only Prevent Aggregation Over a Limited Time Span"

### Supplementary Methods

**General description of Supplementary Tables.** Supplementary Table S1 lists all abbreviations used in simulations. When letters from these abbreviations appear together (i.e. with no intervening spaces), they denote a molecular complex with at least one chaperone or cochaperone (i.e., T + K → TK indicates formation of the ATP-DnaK complex). Supplementary Tables S2-S10 contain the full set of reactions and rate constants used throughout the simulations. Supplementary Table S11 comprises folding/unfolding rates and thermodynamic stabilities for 61 small, single domain proteins. Supplementary Table S12 provides simulation durations and time intervals for each simulation performed. Supplementary Table S13 contains the concentrations of client-proteins and chaperones used in all simulations. Supplementary Table S14 and S15 show comparisons between CHAMPION70 computational predictions and experimental data. Supplementary Table S16 compares *E. Coli* Hsp70 (DnaK) rate constants used in CHAMPION70 with experimentally determined Hsp70 rate constants in humans.

**Computational assessment of standard-state chemical potentials for the generation of landscapes.** Standard-state chemical potentials for monomeric species were obtained from their forward and reverse rate constants. For a reversible transition between two monomeric states, the standard-state free-energy difference is

$$\Delta\mu^{o} = -T * R * \ln\left(\frac{k_{forward}}{k_{reverse}}\right) \quad , \tag{1}$$

where T is temperature (Kelvin), R is Boltzmann's constant, $k_{forward}$ and $k_{reverse}$ are the corresponding rate constants.

For visualization, we set the high energy unfolded state immediately after dilution with denaturant (U') as the reference state with $\mu^{o} = 0$ kcal/mol. For example, using the folding and unfolding rates of firefly luciferase (fluc), the standard-state chemical potential of the native state is

$$\Delta\mu^{o} = -\,298.15\,K * 0.001986\frac{kcal}{mol*K} * \ln\left(\frac{0.12\,sec^{-1}}{0.000035\,sec^{-1}}\right) = -4.82\frac{kcal}{mol} \ . \tag{2}$$

Transition-state barrier heights were computed using a relation derived from Eyring's equation. For a reaction with rate constant k

$$k = k_{max} * e^{\frac{-\Delta\mu^{o\ddagger}}{RT}} \quad , \tag{3}$$

where $k_{max}$ is the diffusion-limited maximum rate constant. Rearranging relation (3) enables determining $\Delta\mu^{o\ddagger}$

$$\Delta\mu^{o\ddagger} = -T * R * \ln\left(\frac{k}{k_{max}}\right) \quad . \tag{4}$$

For instance, the transition-state barrier height for the conversion of the unfolded state to native fluc is

$$\Delta\mu^{o\ddagger} = -298.15K * 0.001986 \frac{kcal}{mol*K} * \ln\left(\frac{0.12\ sec^{-1}}{2*10^{9}\ sec^{-1}}\right) = 13.94 \frac{kcal}{mol} \ . \tag{5}$$

For nucleated polymerization, the standard-state free energy of an aggregate of size n is constructed from a nucleation step followed by elongation steps. For example, trimer formation requires one nucleation event and two elongation events

$$M^* + M^* = A_2 \tag{6}$$

$$\Delta\, G^0 nucleation = G^0_{A2} - 2G^0_{M^*} \tag{7}$$

$$A_2 + M^* = A_3 \tag{8}$$

$$\Delta G^0_{A3} = \Delta G^0 nucleation + 1 * \Delta G^0 elongation \tag{9}$$

$$A_3 + M^* = A_4 \tag{10}$$

$$\Delta\, G^0_{A4} = \Delta G^0 nucleation + 2 * \Delta G^0 elongation \tag{11}$$

$$\dots$$

$$A_{\,n-1} + M^* = A_n \tag{12}$$

$$\Delta G^0_{An} = \Delta G^0 nucleation + [n-2] * \Delta G^0 elongation \tag{13}$$

$$\Delta G^0_{nucl,pmb} = \frac{\Delta G^0 nucleation}{2} \tag{14}$$

$$\Delta G^0_{An,pmb} = \frac{\Delta G^0 nucleation + [n-2]*\Delta G^0 elongation}{n} \quad , \tag{15}$$

where the pmb subscript denotes quantities on a per-monomer basis. For sufficiently large n, the total $\Delta G^0$ approaches the per-monomer value $\Delta G^0_{An,pmb}$.

**Updates to the CHAMP70 model.** Here, two refinements to the CHAMP70 program[1] were introduced to improve the treatment of ATP hydrolysis-driven kinetics and to better reproduce experimentally measured ATPase behavior. These refinements are also implemented in the CHAMPION70 model, which expands upon CHAMP70 program to include aggregation pathways and fold-promoting interactions with misfolded client-proteins.

The first update pertains to the treatment of nucleotide and phosphate concentrations. Our previously published CHAMP70 kinetic model only kept ATP concentration constant. We now also fix the concentrations of ADP and inorganic phosphate ($P_i$). This approach is based on experimental evidence showing that cells maintain stable concentrations of these species, with a stable ATP:ADP ratio of approximately 30:1 and a relatively constant $P_i$ concentration.[2] This treatment ensures a continuous energy-driven flux through the system, representing the biological function of the Hsp70 chaperone system.[3]

The second update involves the ATP hydrolysis rate constant in the presence of DnaJ. In CHAMP70, this rate constant had previously been optimized to reproduce experimentally measured DnaJ dependent stimulation of DnaK ATPase activity obtained from quenched-flow experiments performed in the absence of client-protein.[4] When the model was transitioned from Gepasi to COPASI, the numerical behavior changed so that the previously optimized value no longer reproduced the experimentally observed DnaJ dependent hydrolysis rates. For this reason, the ATP hydrolysis rate constant in the presence of DnaJ was re-optimized using the same experimental measurements (Fig. S8), and the resulting value was applied to reaction steps corresponding to ATP hydrolysis in the presence of DnaJ, with and without client-protein (reaction steps 15 and 27).

**Statistical justification for a single occupied DnaK binding-site per client protein in live cells and for CHAMPION70 binding-site parameterization.** In order to evaluate whether client-proteins with multiple predicted binding sites routinely engage multiple chaperones simultaneously, we analyzed the proteome-wide DnaK binding data from Calloni *et al.*[5] To quantify actual steady-state DnaK engagement across the *E. Coli* proteome, Calloni *et al.*[5] used SILAC pulldowns to measure the DnaK-bound client abundance relative to the total client-protein abundance in soluble lysate, defining this ratio as the relative enrichment factor, according to

$$REF = \frac{[Client_{bound}]}{[Client_{total}]} \ . \tag{16}$$

Calloni *et al.*[5] reported experimentally determined average REF values of ca. 0.05 for DnaK-enriched clients (i.e., clients displaying the highest measured chaperone binding), 0.01 for medium-enriched clients, and < 0.001 for less-enriched clients.[5] While they did not directly measure the number of DnaK molecules bound to an individual chain, their experimental REF values enabled us to calculate the probability of simultaneous multi-DnaK binding to a single protein chain.

In order to model the degree of single-chain occupancy from the above population measurements, we defined the number of bound DnaK chaperones per client as $X$, a Poisson random variable.[6] In our statistical derivation, the entire protein pool yields a fixed REF value, while an individual chaperone-occupied client carrying either a single DnaK chaperone or multiple DnaKs, varies randomly across individual protein chains. According to this model, the probability that a client molecule is bound to at least one DnaK chaperone, denoted as $P(X \geq 1)$, corresponds to the measured REF value, according to the Poisson probability distribution

$$P(X \geq 1) = 1 - P(X = 0) = 1 - e^{-\lambda} = REF \ , \tag{17}$$

where $\lambda$ is the single Poisson parameter. For DnaK-enriched clients displaying maximal binding (REF = 0.05), solving equation (17) yields a Poisson parameter $\lambda \approx 0.0513$. This value denotes the average number of bound DnaK chaperones per client chain across the total protein pool.

To evaluate multi-chaperone binding within the chaperone-bound population, we also calculated the conditional probability that a bound client carries two or more DnaK molecules, according to

$$P(X \geq 2 \mid X \geq 1) = \frac{P(X\geq 2)}{P(X\geq 1)} = \frac{1-e^{-\lambda}(1+\lambda)}{1-e^{-\lambda}}\,. \qquad (18)$$

Substituting $\lambda \approx 0.0513$ into equation (18) gives $P(X \geq 2 \mid X \geq 1) \approx 0.026$. This result means that more than 97% of bound client molecules are predicted to carry only a single DnaK chaperone at any given instant in live *E. coli* cells. Modeling DnaK binding as a Poisson process assumes that chaperone engagement at distinct binding sites occurs independently. If initial DnaK binding were to sterically block access to adjacent sites (negative cooperativity), the true percentage of simultaneous binding would be even lower than 2.6%. Conversely, if initial binding were to cause local unfolding that exposes additional sites or recruits further chaperones (positive cooperativity), the population could shift toward a bimodal distribution where client chains are either completely unbound or carry multiple chaperones.

We also evaluated whether the experimental conditions in Calloni *et al.* reflect physiological conditions in living cells. While *E. coli* cells were grown at a physiological temperature of 37 °C, apyrase was added during cell lysis to rapidly deplete free ATP.[5] Depleting ATP locks DnaK in its high-affinity ADP-bound state and prevents substrate release, effectively reducing the dissociation rate ($k_{off}$) toward zero and trapping pre-existing DnaK-client complexes. In living cells with active ATP cycling, GrpE-mediated nucleotide exchange accelerates client dissociation by increasing $k_{off}$, shortening steady-state residence times and making simultaneous multi-DnaK binding even less likely in true physiological conditions than in ATP-depleted lysates. In addition, the lysate contained lower relative levels of DnaJ (DnaK:GrpE:DnaJ at ~30:20:1)[5] than the physiological concentrations modeled in CHAMPION70 (5:2:1), which could moderately decrease the association rate ($k_{on}$). Preventing client dissociation during lysis, however, ensures that the measured REF value serves as a conservative upper bound value for the steady-state DnaK occupancy in the cell.

**General principles of detailed balance.** The principle of detailed balance, or microscopic reversibility, applies to kinetic schemes at thermodynamic equilibrium. Consider a closed reaction loop:

$$A \overset{K_1}{\rightleftarrows} B \ ,$$

$$B \overset{K_2}{\rightleftarrows} C \ ,$$

$$C \overset{K_3}{\rightleftarrows} A \ .$$

The principle of detailed balance requires that ΔG° for a complete loop $A \rightleftarrows A$ must be 0.[7] Since

$$\Delta G^\circ = -RT \ln K \tag{19}$$

where R is Boltzmann's constant and T is temperature, therefore,

$$0 = \Delta G_1{}^\circ + \Delta G_2{}^\circ + \Delta G_3{}^\circ = -RT\left(\ln K_1 + \ln K_2 + \ln K_3\right) = -RT \ln\left(K_1 K_2 K_3\right) \ . \tag{20}$$

Thus

$$K_1 K_2 K_3 = 1 \ . \tag{21}$$

Equation (21) is known as the Wegscheider's condition.[8,9] In general, if there are n reactions in a reaction loop, the equilibrium constant for the reaction system is $K_n$, and Wegscheider's condition is equivalent to[8,9]

$$\prod_{i=1}^{n} K_i = 1 \tag{22}$$

In many proposed kinetic models, some reactions may be treated as effectively irreversible. Even in these cases, one can still assess whether a proposed reaction network is compatible with detailed balance by examining its reaction loops.

An irreversible step can be treated as having an infinite equilibrium constant. If it is still possible to satisfy Wegscheider's condition under this assumption, the loop is considered legal and consistent with detailed balance. Otherwise, the reaction loop is illegal and inconsistent with detailed balance. Examples of legal and illegal loops are shown in Supplementary Fig. S2 and a more detailed review of legal and illegal loops is provided by Stanbury and Hoffman.[7]

**Principles of steady-state kinetics.** A steady-state is defined as a condition according to which the concentrations of all chemical species ($c_i$) remain constant with time ( $\frac{dc_i}{dt} = 0$), while the system maintains a non-zero net flux ($J^{net} \neq 0$) with a non-zero chemical potential difference driving the system ($\Delta\mu \neq 0$).

In contrast, at thermodynamic equilibrium all net fluxes vanish ($J^{net} = 0$) and all chemical potential differences are zero.[10,11]

To illustrate the principles of steady-state kinetics, consider the simple reversible reaction $A \rightleftarrows B$, with forward and reverse rate constants $k_1$ and $k_{-1}$, in an open system where external forces maintain the concentrations $c_A$ and $c_B$ at fixed values:

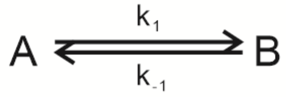


Given these external forces hold the concentrations constant, the system is at steady-state. Since the system is not at equilibrium, the net flux: $J^{net} = J_+ - J_- \neq 0$, where $J_+ = k_1 c_A$ and $J_- = k_{-1} c_B$, are individually non-zero. The chemical potentials $\mu_A = \mu_B^0 + k_B T \ln c_A$ and $\mu_B = \mu_B^0 + k_B T \ln c_A$ yield a chemical potential difference

$$\Delta\mu = \mu_B - \mu_A = \mu_B^0 - \mu_A^0 + k_B T \ln \frac{c_B}{c_A} \tag{23}$$

Using relation: $\frac{k_1}{k_{-1}} = e^{\frac{\mu_B^0 - \mu_A^0}{k_B T}}$, equation (23) becomes

$$\Delta\mu = \mu_B^0 - \mu_A^0 + k_B T \ln\frac{c_B}{c_A} = k_B T \ln\frac{k_1}{k_{-1}} + k_B T \ln\frac{c_B}{c_A} = k_B T \ln\frac{k_1 c_B}{k_{-1} c_A} = k_B T \ln\left(\frac{J_-}{J_+}\right) \quad (24)$$

Substituting relation (6) into the entropy-production expression yields

$$-J^{net}\Delta\mu = k_B T(J_+ - J_-)\ln\left(\frac{J_+}{J_-}\right) \geq 0 \quad . \quad (25)$$

For any reversible reaction step, the chemical potential is therefore tied to the ratio of forward and reverse fluxes. A non-zero net flux requires $J_+ \neq J_-$, which in turn implies $\Delta\mu \neq 0$. Thus, the equality in the entropy-production inequality is not satisfied, and the system is maintained in a nonequilibrium steady-state.[10,11] Note that the far left of equation (25) is the entropy-production rate, which is strictly positive in an open system away from equilibrium. The equality of the inequality holds in closed equilibrium systems where the zero-entropy condition is satisfied.

Importantly, given that steady-state conditions enforce non-zero fluxes and non-zero chemical potential differences, detailed balance does not hold.[10,11] Detailed balance requires $J_+ = J_-$ for every reaction, which is only possible at thermodynamic equilibrium. Only when all driving forces are removed and the system is closed, can it relax to equilibrium, where the detailed-balance and Wegscheider's condition apply.[10,11]

**Steady-state conditions in CHAMPION70.** CHAMPION70 operates as a nonequilibrium steady-state (NESS) driven by the hydrolysis of ATP to ADP + inorganic phosphate ($P_i$), under constant temperature and pressure. According to this model, the concentrations of ATP, ADP, and $P_i$ are held fixed, which in turn fixes the corresponding chemical potentials and establishes a constant, non-zero chemical potential difference for ATP hydrolysis. This chemical potential difference provides the thermodynamic driving force that sustains non-zero net fluxes throughout the relevant reaction network.

ATP hydrolysis is strongly exergonic ($\Delta\mu^o < 0$), and the reaction step directly coupled to ATP hydrolysis therefore carries a positive net flux. Although the reaction network is open and exchanges energy with its surroundings, the combined system of network plus surroundings is closed, and must obey the second law of thermodynamics. Under steady-state conditions, the entropy production rate satisfies

$$-J^{ss}\Delta\mu \geq 0 \tag{26}$$

with the equality applying only at thermodynamic equilibrium.[10,11] Given that ATP hydrolysis maintains a non-zero chemical potential difference and a non-zero net flux, the system cannot satisfy the above equality and therefore cannot be at equilibrium. As long as ATP, ADP, and $P_i$ concentrations are held constant, the network remains under a nonequilibrium steady-state.

In the sections that follow, we will examine how the nonequilibrium driving force supplied by ATP hydrolysis affects reaction loops within a reaction network like CHAMPION70. First, we show that any loop containing an ATP hydrolysis step cannot satisfy detailed balance. Then, we prove that loops that do not contain ATP hydrolysis but share a reaction edge with the original loop cannot, likewise, satisfy detailed balance because the nonequilibrium flux propagates through the shared edge. Finally, we show that even loops that neither contain ATP hydrolysis nor share an edge with a driven loop are maintained in a nonequilibrium steady-state through their coupling to the larger network. Together, the above results demonstrate that no loop in CHAMPION70 is required to satisfy detailed balance, regardless of whether the loop contains an ATP hydrolysis step.

**Detailed balance does not apply to kinetic loops that include a reaction step operating under steady state conditions.** Consider the following elementary loop composed of three species A, B, and C, where the reaction $A \rightleftarrows B$ is coupled to ATP hydrolysis according to

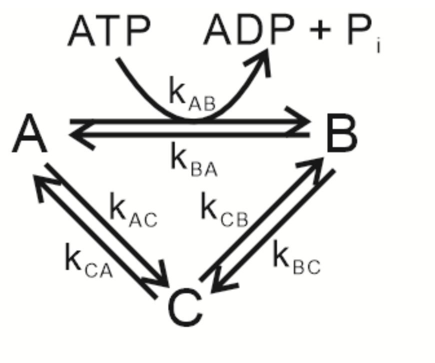


Given that ATP hydrolysis is strongly exergonic and that the concentrations of ATP, ADP, and $P_i$ are held fixed, the coupled reaction carries a positive, non-zero flux: $J_{AB}^{net} = k_{AB}P_A^{SS} - k_{BA}P_B^{SS} > 0$. Given that ATP, ADP, and $P_i$ are maintained at constant concentrations, all nucleotide-dependent reaction steps can be written using effective rate constants whose numerical values include the fixed nucleotide concentrations. This is why ATP, ADP, and $P_i$ do not appear explicitly in net-flux rate equations and steady-state probability rate equations. Below, we write the steady-state probability rate equations, where each steady state probability ($P_i^{SS}$) denotes the normalized concentration of state i. At steady state, the $P_i^{SS}$ are time-independent because the total incoming probability flux balances the outgoing flux, even though nonzero fluxes can circulate through the network. The steady-state probability rate equations of species A, B, and C are

$$\frac{dP_A(t)}{dt} = -(k_{AB} + k_{AC})P_A^{SS} + k_{BA}P_B^{SS} + k_{CA}P_C^{SS} = 0 \quad , \tag{27}$$

$$\frac{dP_B(t)}{dt} = k_{AB}P_A^{SS} - (k_{BA} + k_{BC})P_B^{SS} + k_{CB}P_C^{SS} = 0 \quad , \tag{28}$$

$$\frac{dP_C(t)}{dt} = k_{AC}P_A^{SS} + k_{BC}P_B^{SS} - (k_{CA} + k_{CB})P_C^{SS} = 0 \quad . \tag{29}$$

Upon defining the net fluxes through each reaction as

$$J_{AB}^{net} = k_{AB}P_A^{SS} - k_{BA}P_B^{SS} \quad , \tag{30}$$

$$J_{BC}^{net} = k_{BC}P_B^{SS} - k_{CB}P_C^{SS} \quad , \tag{31}$$

$$J_{CA}^{net} = k_{CA}P_C^{SS} - k_{AC}P_A^{SS} \quad , \tag{32}$$

we can rearrange the steady-state rate equations and express them in terms of the net fluxes. For species A

$$\frac{dP_A(t)}{dt} = -\left(k_{AB}P_A^{SS} - k_{BA}P_B^{SS}\right) + \left(k_{CA}P_C^{SS} - k_{AC}P_A^{SS}\right) = -J_{AB}^{net} + J_{CA}^{net} = 0 \ . \tag{33}$$

The above implies:

$$J_{AB}^{net} = J_{CA}^{net} \ . \tag{34}$$

Repeating this process for species B and C yields

$$J_{AB}^{net} = J_{BC}^{net} \ , \tag{35}$$

$$J_{BC}^{net} = J_{CA}^{net} \ . \tag{36}$$

Since the ATP-hydrolysis-coupled reaction ($J_{AB}^{net}$) has a positive net flux under steady-state conditions, each step in the loop (e.g. $A \rightleftarrows B$) must also have a positive net flux. Detailed balance requires zero net flux throughout any reaction loop, which is incompatible with an overall positive net flux. Therefore, any loop containing an ATP-hydrolysis-coupled reaction under steady-state (or pseudo-steady-state conditions) cannot satisfy detailed balance. Thus, enforcement of Wegscheider's condition is not required. The nonequilibrium driving force introduced by steady-state ATP hydrolysis propagates through the entire loop, ensuring the loop operates under a nonequilibrium steady state rather than at thermodynamic equilibrium.

**Propagation of nonequilibrium flux to adjacent loops sharing a reaction edge.** Now, consider a reaction network containing two elementary loops. The first loop consists of species A, B, and C, where the reaction $A \rightleftarrows B$ is coupled to ATP hydrolysis and therefore carries a positive net flux. The second loop consists of species B, C, and D, and shares the reaction edge $B \rightleftarrows C$ with the ATP hydrolysis driven loop

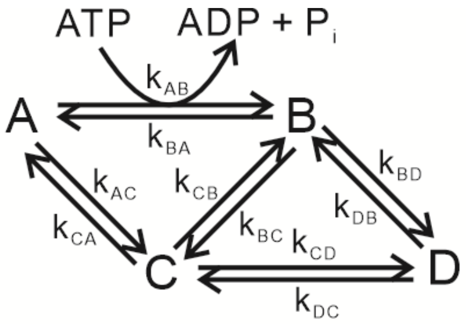


As before, we assume that the ATP-hydrolysis-coupled step $A \rightleftarrows B$ has a positive, non-zero flux: $J_{AB}^{net} = k_{AB}P_A^{SS} - k_{BA}P_B^{SS} > 0$. In addition, ATP, ADP, and $P_i$ continue to be held at fixed concentrations, so all nucleotide-dependent steps are represented by effective rate constants. The steady-state probability rate equations for the four species are:

$$\frac{dP_A(t)}{dt} = -(k_{AB} + k_{AC})P_A^{SS} + k_{BA}P_B^{SS} + k_{CA}P_C^{SS} = 0 \ , \tag{37}$$

$$\frac{dP_B(t)}{dt} = k_{AB}P_A^{SS} - (k_{BA} + k_{BC} + k_{BD})P_B^{SS} + k_{CB}P_C^{SS} + k_{DB}P_D^{SS} = 0 \ , \tag{38}$$

$$\frac{dP_C(t)}{dt} = k_{AC}P_A^{SS} + k_{BC}P_B^{SS} - (k_{CA} + k_{CB} + k_{CD})P_C^{SS} + k_{DC}P_D^{SS} = 0 \ , \tag{39}$$

$$\frac{dP_D(t)}{dt} = k_{BD}P_B^{SS} + k_{CD}P_C^{SS} - (k_{DB} + k_{DC})P_D^{SS} = 0 \ . \tag{40}$$

The net fluxes through each reaction are

$$J_{AB}^{net} = k_{AB}P_A^{SS} - k_{BA}P_B^{SS} \ , \tag{41}$$

$$J_{BC}^{net} = k_{BC}P_B^{SS} - k_{CB}P_C^{SS} \ , \tag{42}$$

$$J_{CA}^{net} = k_{CA}P_C^{SS} - k_{AC}P_A^{SS} \ , \tag{43}$$

$$J_{BD}^{net} = k_{BD}P_B^{SS} - k_{DB}P_D^{SS} \ , \tag{44}$$

$$J_{DC}^{net} = k_{DC}P_D^{SS} - k_{CD}P_C^{SS} \ . \tag{45}$$

Substituting these into the steady-state rate equations yields

$$J_{AB}^{net} = J_{CA}^{net} \ , \tag{46}$$

$$J_{AB}^{net} = J_{BC}^{net} + J_{BD}^{net} \ , \tag{47}$$

$$J_{CA}^{net} = J_{BC}^{net} + J_{DC}^{net} \ , \tag{48}$$

$$J_{BD}^{net} = J_{DC}^{net} \quad . \tag{49}$$

The above equalities show that the net flux through the ATP-hydrolysis-driven reaction $A \rightleftarrows B$ propagates into the adjacent loop through the “shared edge” $B \rightleftarrows C$. Given that $J_{AB}^{net} > 0$, the above relations require that at least one of the fluxes $J_{BC}^{net}$, $J_{BD}^{net}$, or $J_{DC}^{net}$ must also be non-zero and positive. As a consequence, the loop composed of species B, C, and D cannot satisfy detailed balance. Thus, any loop that shares a reaction edge with an ATP-hydrolysis-driven loop is necessarily maintained in a nonequilibrium steady-state and thus cannot satisfy detailed balance.

**Steady-state maintenance in loops indirectly coupled to ATP hydrolysis.** Finally, consider an elementary loop composed of species D, E, and F that do not contain an ATP-hydrolysis-coupled reaction and do not share a reaction edge with any loop according to

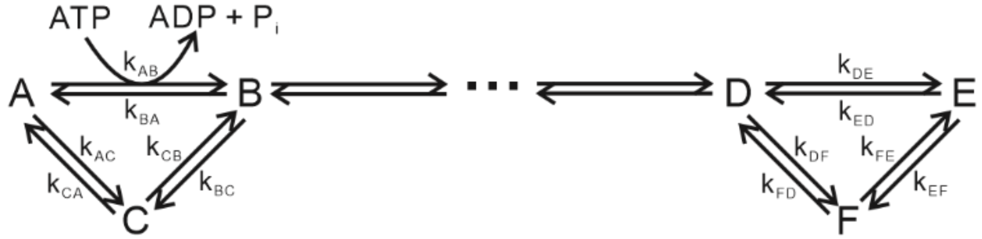


As schematically illustrated above, the DEF loop is indirectly coupled to the ATP-hydrolysis-driven ABC loop through intermediate reactions that are not explicitly shown (denoted by three dots). This coupling renders this loop a subsystem of a larger open reaction network. Given that the entire network is maintained in a nonequilibrium steady-state by ATP hydrolysis, the steady-state probabilities $P_i^{SS}$ of all species belonging to this loop are determined by the full network dynamics rather than by the internal reactions within the DEF loop.

Let $P_D^{SS}$, $P_E^{SS}$, and $P_F^{SS}$ denote the steady-state probabilities of species D, E, and F. The key point is that $P_D^{SS}$ is fixed by reactions connecting species D to the ATP hydrolysis driven portion of the network, even though those reactions are not shown explicitly here. Given that $P_D^{SS}$ is imposed by the global steady-state, it does not adopt the equilibrium value that would arise if the loop were isolated.

The steady-state probability equations for the internal reactions of the loop, treating $P_D^{SS}$ as a fixed boundary condition imposed by the overall steady-state of the network, are

$$\frac{dP_D(t)}{dt} = -(k_{DE} + k_{DF})P_D^{SS} + k_{ED}P_E^{SS} + k_{FD}P_F^{SS} = 0 \ , \quad (50)$$

$$\frac{dP_E(t)}{dt} = k_{DE}P_D^{SS} - (k_{ED} + k_{EF})P_E^{SS} + k_{FE}P_F^{SS} = 0 \ , \quad (51)$$

$$\frac{dP_F(t)}{dt} = k_{DF}P_D^{SS} + k_{EF}P_E^{SS} - (k_{FD} + k_{FE})P_F^{SS} = 0 \ . \quad (52)$$

Upon defining the net fluxes as

$$J_{DE}^{net} = k_{DE}P_D^{SS} - k_{ED}P_E^{SS} \ , \quad (53)$$

$$J_{EF}^{net} = k_{EF}P_E^{SS} - k_{FE}P_F^{SS} \ , \quad (54)$$

$$J_{FD}^{net} = k_{FD}P_F^{SS} - k_{DF}P_D^{SS} \ , \quad (55)$$

and rewriting the steady-state equations in terms of the net-flux, yields

$$J_{DE}^{net} = J_{FD}^{net} \ , \quad (56)$$

$$J_{DE}^{net} = J_{EF}^{net} \ , \quad (57)$$

$$J_{EF}^{net} = J_{FD}^{net} \ . \quad (58)$$

Thus, all reactions in the loop carry the same net flux according to

$$J_{DE}^{net} = J_{EF}^{net} = J_{FD}^{net} \ . \quad (59)$$

The above equalities enforce flux propagation through all loops kinetically connected to the steady-state ATP-hydrolysis-driven loop (i.e., the ABC loop), but do not require the fluxes to be zero. In summary, for the loop to satisfy detailed balance, all net fluxes would need to vanish according to

$$J_{DE}^{net} = J_{EF}^{net} = J_{FD}^{net} = 0 \ . \quad (60)$$

The above conditions would require the steady-state probabilities $P_D^{SS}$, $P_E^{SS}$, and $P_F^{SS}$ to adopt their equilibrium values and the rate constants to satisfy Wegscheider's condition. However, given that

$P_D^{SS}$ is fixed by the global nonequilibrium steady-state enforced by ATP hydrolysis under constant ATP and ADP concentration, $P_D^{SS}$ does not assume its equilibrium value. Imposing equilibrium within the DEF loop would therefore require fine-tuning the rate constants to match an externally imposed $P_D^{SS}$, and these rate constants do not necessarily satisfy Wegscheider's condition. Moreover, the steady-state equations can have solutions with non-zero fluxes for arbitrary positive rate constants, demonstrating that the loop is maintained under a nonequilibrium steady-state through its coupling to the larger network. In summary, even though loop does not contain steady-state ATP hydrolysis and does not share a reaction edge with a driven loop, it does not need to satisfy detailed balance.

**Irreversible reactions and the applicability of steady-state flux derivations.** The derivations above were prepared assuming that each reaction step is reversible, with forward and reverse fluxes $J_+$ and $J_-$. In CHAMPION70, however, some steps are irreversible. For an irreversible reaction, the reverse flux is zero, so the net flux reduces to $J^{net} = J_+$. This modification does not alter any of the steady-state flux relations derived in the previous sections. The steady-state probability rate equations depend only on the net fluxes through each reaction, and these equations remain valid whether a reaction is reversible or irreversible. When a reaction is irreversible, its net flux simply becomes the forward flux. The equalities discussed above, relating to net fluxes within a loop or across connected loops, remain unchanged.

A potential concern is that, if $J^{net} = 0$ for an irreversible reaction, the reaction would disappear, suggesting that irreversible reactions cannot be compliant with detailed balance. On the other hand, detailed balance requires zero net flux across every loop. An irreversible step can still be part of a loop that satisfies detailed balance, if other irreversible steps in the loop provide flux in the opposite direction so that the net flux across the loop is zero. The nonequilibrium driving force from ATP hydrolysis, however, generates a non-zero net flux that propagates through connected loops by the steady-state rate equations. Given that there is no opposing flux to cancel it, detailed balance cannot be satisfied anywhere, with the above reaction networks.

Importantly, all reaction loops within the CHAMPION70 (and CHAMP70) reaction network do not need to satisfy detailed balance, or to be compliant with Wegscheider's condition.

**Aggregation outcomes depend strongly on nucleation and misfolding kinetics.** Two kinetic features strongly influence folding and aggregation outcomes in the CHAMPION70 model: the aggregation nucleation rate and the rate for the unfolded state (U) transitioning into the misfolded intermediate (M*). We varied both parameters systematically to determine how they shape folding, aggregation, and chaperone rescue.

In order to evaluate the effect of nucleation rate, we ran CHAMPION70 under strongly aggregation-prone (SAP) conditions and altered the original nucleation rate by two orders of magnitude in both directions. We then simulated the four representative client-proteins, tracking aggregate formation with and without chaperones (Supplementary Fig. S14B-G). Faster nucleation produces more rapid accumulation of fully aggregated species. Under the fastest nucleation conditions (Supplementary Fig. S14F and S14G), the chaperone system provides a noticeably smaller rescue effect. In contrast, the slowest nucleation rate (Supplementary Fig. S14B and S14C) yields the strongest chaperone-mediated suppression of aggregation. Across all nucleation rates, the slow-folding and thermodynamically stable client consistently experiences the greatest extent of rescue by the chaperone.

To examine the influence of the U-to-M* transition, we accelerated this step by two and four orders of magnitude under weakly aggregation-prone (WAP) conditions, where the unfolded client-protein is somewhat kinetically trapped relative to U' and M* (Supplementary Fig. S15B-G). The U to M* step represents one of the kinetic routes to misfolding. Therefore, lowering its barrier increases the kinetic accessibility of misfolding and thus increases aggregation for both unstable clients. Even under these conditions, chaperones still substantially delay aggregate formation. Among the stable clients, the slow-folding variant aggregates more rapidly than the fast-folding one, consistent with the idea that slower folding prolongs the time spent in the unfolded state and increases the likelihood of misfolding and aggregation.

**Monomer-dependent secondary nucleation amplifies aggregation and undermines chaperone protection**. Secondary nucleation is a key mechanism that accelerates aggregate buildup by generating new aggregates from existing ones, bypassing the slow, rate-limiting primary nucleation step. Two types of secondary nucleation have been characterized based on their dependence on misfolded monomer concentration: monomer-dependent secondary nucleation and monomer-independent secondary nucleation. In monomer-dependent secondary nucleation, the surfaces of existing aggregates catalyze the formation of new aggregates.[12] Monomer-independent secondary nucleation, also known as fragmentation, results from the breaking of existing aggregates to give rise to new protein aggregates.[13] To evaluate how secondary nucleation influences aggregation dynamics, we incorporated monomer-dependent secondary nucleation reactions into CHAMPION70 and simulated aggregation kinetics across non-, weakly, and strongly aggregation-prone regimes for four representative client-proteins, both in the absence and presence of the Hsp70 system (Supplementary Fig. S16).

Non-aggregation-prone (NAP) proteins (Supplementary Fig. S16C, D, E, F) do not aggregate under any conditions, as aggregation pathways are turned off. Under WAP conditions, secondary nucleation does not substantially accelerate aggregation in the absence of chaperones (Supplementary Fig. S16G, H). In the presence of Hsp70 (Supplementary Fig. S16I, J), secondary nucleation increases the final aggregate concentration for thermodynamically unstable clients, which retain larger populations of non-native species even with chaperone action. Once initial aggregates form, these non-native species provide substrate for surface-catalyzed amplification. Thermodynamically stable clients remain largely protected because their folded states are strongly favored and chaperone interactions further suppress aggregation-prone conformations.

Under (SAP) conditions, secondary nucleation noticeably accelerates aggregation for all clients, both in the absence and presence of Hsp70. Early aggregate formation provides substantial surface area for autocatalytic amplification, producing faster increases in normalized aggregate levels and a pronounced intermediate aggregation shelf (Supplementary Fig. S16K, L). This effect is especially evident for fast-folding, thermodynamically stable clients, indicating that, once aggregate surfaces appear due to any preexisting aggregate, secondary nucleation can override intrinsic folding advantages. In the presence of chaperones (Supplementary Fig. S16M, N), the same behavior is observed, with elevated aggregation shelves and reduced times to full aggregation, demonstrating that secondary nucleation amplifies aggregate populations and disrupts chaperone-mediated kinetic protection under strongly aggregation-prone conditions.

**High native-state kinetic stability enhances chaperone activity.** In order to further examine how folding kinetics influence interactions with the Hsp70 system, we altered RNase $H^D$ to reduce its kinetic stability. Specifically, we increased both its folding and unfolding rates (Supplementary Fig. S17A) to lower the kinetic-barrier between the unfolded (U) and native (N) states relative to the original protein (Supplementary Fig. S17B). We denote this version of RNase $H^D$ as fast-folding RNase $H^D$.

We compared simulations of the fast-folding RNase $H^D$ and original RNase $H^D$ across NAP, WAP, and SAP regimes. Under NAP and WAP conditions, both variants exhibit identical aggregation behavior (Supplementary Fig. S17C, D, F, G). However, their interactions with Hsp70 differ. The original RNase $H^D$ engages in short-lived chaperone interactions, while the modified RNase $H^D$ (fast unfold, fast fold) shows essentially no chaperone interaction under these conditions (Supplementary Fig. S17E, H).

The largest effect of altered kinetic stability emerges under SAP conditions (Supplementary Fig. S17I, J). In the absence of Hsp70, the modified RNase $H^D$ aggregates more slowly than the original protein, consistent with rapid folding providing partial protection. In contrast, when Hsp70 is present, aggregation is substantially delayed only for the original RNase $H^D$. These results indicate that, while rapid folding can reduce aggregation in the absence of chaperones, slow unfolding, corresponding to a high kinetic-barrier between N and U, is more important for effective chaperone-mediated protection from aggregation.

### Supplementary Tables

**Supplementary Table S1.** List of abbreviations for all the CHAMP70 and CHAMPION70 individual species that are not involved in a complex with a chaperone. Additional species appearing in the kinetic schemes of Supplementary Tables S2 – S10 arise from combinations of molecules listed in this table. For instance, the T + K ⇄ TK kinetic step denotes the reversible formation of the ATP-DnaK complex TK, starting from ATP and DnaK.

| Abbreviation | Abbreviation meaning |
|---|---|
| U' | Denaturant-unfolded state immediately after dilution into an excess of denaturant-free solution (conformationally unrelaxed) |
| U | Unfolded state |
| N | Native state |
| I | Intermediate state |
| M* | Misfolded state |
| K | DnaK |
| J | DnaJ |
| E | GrpE |
| T | ATP |
| D | ADP |
| P | $P_i$ |
| $A_2$ | Dimer resulting from reaction of two M* |
| $A_{N+1}$ | Aggregate of size (N + 1), resulting from (N + 1) M* molecules |

| | |
|---|---|
| $A_{20}$ | Final aggregate (20-mer), resulting from 20 M* molecules |
| = or ⇄ | Reversible reaction |
| -> or → | Irreversible reaction |

**Supplementary Table S2**. Rate constants employed in the protein-folding and aggregation simulations in the absence of chaperones. The aggregation rate constants listed here correspond to weakly aggregation-prone (WAP) client-proteins. Rate constants for non-aggregation-prone (NAP) and strongly aggregation-prone (SAP) clients are provided in Supplementary Table S4.

| Reaction | Forward step # | Forward rate constant | Reverse step # | Reverse rate constant |
|---|---|---|---|---|
| $U \rightleftarrows N$ | f | protein-dependent | u | protein-dependent |
| $U \rightleftarrows M^*$ | 42 | $0.000004\ s^{-1}$ | 43 | $2.6\times10^{-9}\ s^{-1}$ |
| $U' \rightleftarrows U$ | 44 | $4000\ s^{-1}$ | 45 | $0.6\ s^{-1}$ |
| $U' \rightleftarrows M^*$ | 46 | $0.0000048\ s^{-1}$ | 47 | $0.0000011\ s^{-1}$ |
| $2 * M^* \rightleftarrows A_2$ | 75 | $150\ M^{-1}s^{-1}$ | 76 | $1.92\times10^{-12}\ s^{-1}$ |
| $A_2 + M^* \rightleftarrows A_3$ | 77 | $6.06\times10^{9}\ M^{-1}s^{-1}$ | 78 | $7.82\times10^{-12}\ s^{-1}$ |
| $A_3 + M^* \rightleftarrows A_4$ | N/A | $6.06\times10^{9}\ M^{-1}s^{-1}$ | N/A | $7.82\times10^{-12}\ s^{-1}$ |
| $A_4 + M^* \rightleftarrows A_5$ | N/A | $6.06\times10^{9}\ M^{-1}s^{-1}$ | N/A | $7.82\times10^{-12}\ s^{-1}$ |
| $A_5 + M^* \rightleftarrows A_6$ | N/A | $6.06\times10^{9}\ M^{-1}s^{-1}$ | N/A | $7.82\times10^{-12}\ s^{-1}$ |
| $A_6 + M^* \rightleftarrows A_7$ | N/A | $6.06\times10^{9}\ M^{-1}s^{-1}$ | N/A | $7.82\times10^{-12}\ s^{-1}$ |
| $A_7 + M^* \rightleftarrows A_8$ | N/A | $6.06\times10^{9}\ M^{-1}s^{-1}$ | N/A | $7.82\times10^{-12}\ s^{-1}$ |
| $A_8 + M^* \rightleftarrows A_9$ | N/A | $6.06\times10^{9}\ M^{-1}s^{-1}$ | N/A | $7.82\times10^{-12}\ s^{-1}$ |
| $A_9 + M^* \rightleftarrows A_{10}$ | N/A | $6.06\times10^{9}\ M^{-1}s^{-1}$ | N/A | $7.82\times10^{-12}\ s^{-1}$ |
| $A_{10} + M^* \rightleftarrows A_{11}$ | N/A | $6.06\times10^{9}\ M^{-1}s^{-1}$ | N/A | $7.82\times10^{-12}\ s^{-1}$ |
| $A_{11} + M^* \rightleftarrows A_{12}$ | N/A | $6.06\times10^{9}\ M^{-1}s^{-1}$ | N/A | $7.82\times10^{-12}\ s^{-1}$ |
| $A_{12} + M^* \rightleftarrows A_{13}$ | N/A | $6.06\times10^{9}\ M^{-1}s^{-1}$ | N/A | $7.82\times10^{-12}\ s^{-1}$ |

| | | | | |
|---|---|---|---|---|
| $A_{13} + M^* \rightleftarrows A_{14}$ | N/A | $6.06\times10^9$ $M^{-1}s^{-1}$ | N/A | $7.82\times10^{-12}$ $s^{-1}$ |
| $A_{14} + M^* \rightleftarrows A_{15}$ | N/A | $6.06\times10^9$ $M^{-1}s^{-1}$ | N/A | $7.82\times10^{-12}$ $s^{-1}$ |
| $A_{15} + M^* \rightleftarrows A_{16}$ | N/A | $6.06\times10^9$ $M^{-1}s^{-1}$ | N/A | $7.82\times10^{-12}$ $s^{-1}$ |
| $A_{16} + M^* \rightleftarrows A_{17}$ | N/A | $6.06\times10^9$ $M^{-1}s^{-1}$ | N/A | $7.82\times10^{-12}$ $s^{-1}$ |
| $A_{17} + M^* \rightleftarrows A_{18}$ | N/A | $6.06\times10^9$ $M^{-1}s^{-1}$ | N/A | $7.82\times10^{-12}$ $s^{-1}$ |
| $A_{18} + M^* \rightleftarrows A_{19}$ | N/A | $6.06\times10^9$ $M^{-1}s^{-1}$ | N/A | $7.82\times10^{-12}$ $s^{-1}$ |
| $A_{19} + M^* \rightleftarrows A_{20}$ | N/A | $6.06\times10^9$ $M^{-1}s^{-1}$ | N/A | $7.82\times10^{-12}$ $s^{-1}$ |

**Supplementary Table S3.** Folding (forward) and unfolding (reverse) rate constants for the client-proteins analyzed in this work.

| Protein | PDB ID | Kinetic / thermodynamic stability | Quadrant (defined in Fig. 5A) | Reaction | Forward rate const. ($s^{-1}$) | Reverse rate const. ($s^{-1}$) |
|---|---|---|---|---|---|---|
| Ubiquitin | 1UBQ | Fast/stable | I | U ⇄ N | 1480.299 | 0.0011137 |
| mAcP | 1APS | Slow/stable | II | U ⇄ N | 0.202 | $1.23\times10^{-4}$ |
| RNase $H^D$ (CHAMP70) | 1F21 | Slow/stable | II | U ⇄ N | 0.03 | $2.3768\times10^{-6}$ |
| RNase $H^D$* (CHAMPION70) | 1F21 | Slow/stable | II | U ⇄ I | 100 | 3.42 |
| | | | | I ⇄ N | 0.033934 | $6.6\times10^{-5}$ |
| Modified RNase $H^D$ (fast unfold, fast fold) (CHAMPION70) | 1F21 | Slow/stable | II | U ⇄ I | 100 | 3.42 |
| | | | | I ⇄ N | 3393.4 | 6.6 |
| Luciferase* | 1LCI | Slow/stable | II | U ⇄ N | 0.12 | $3.50\times10^{-5}$ |
| drkN SH3 | 2A36 | Slow/unstable | III | U ⇄ N | 1.5 | 0.7 |
| Formin-binding protein (FBP) | 1E0L | Fast/unstable | IV | U ⇄ N | 40134 | 5431.65 |

*rate constants were deduced from fitting experimental data with COPASI.

**Supplementary Table S4.** Key reactions and rate constants for the three aggregation-propensity regimes simulated in this work.

| **Aggregation-propensity regime** | **Kinetic step** | **Forward rate constant ($s^{-1}$)** | **Reverse rate constant ($s^{-1}$)** |
|---|---|---|---|
| Non-aggregation-prone (NAP) | U’ ⇄ U | 4,000 $s^{-1}$ | 0.6 $s^{-1}$ |
| | U’ ⇄ M* | -- | -- |
| | U ⇄ M* | -- | -- |
| Weakly aggregation-prone (WAP) | U’ ⇄ U | 4,000 $s^{-1}$ | 0.6 $s^{-1}$ |
| | U’ ⇄ M* | 0.0000048 $s^{-1}$ | 0.0000011 $s^{-1}$ |
| | U ⇄ M* | 0.000004 $s^{-1}$ | $2.6\times10^{-9}$ $s^{-1}$ |
| Strongly aggregation-prone (SAP) | U’ ⇄ U | 480 $s^{-1}$ | 110 $s^{-1}$ |
| | U’ ⇄ M* | 4,000 $s^{-1}$ | 0.6 $s^{-1}$ |
| | U ⇄ M* | 0.00004 $s^{-1}$ | $2.6\times10^{-8}$ $s^{-1}$ |

**Supplementary Table S5.** Elementary kinetic steps and associated rate constants for the original CHAMP70 model, reproduced from Sekhar *et al.*[1]

| Reaction | Forward step # | Forward rate | Reverse step # | Reverse rate |
|---|---|---|---|---|
| U ⇄ N | f | protein-dependent | u | protein-dependent |
| U + TK ⇄ UTK | 1 | 280,000 $M^{-1}s^{-1}$ | 2 | 3.7 $s^{-1}$ |
| U + DK ⇄ UDK | 3 | 1,500 $M^{-1}s^{-1}$ | 4 | 0.001 $s^{-1}$ |
| U + K ⇄ UK | 5 | 7,500 $M^{-1}s^{-1}$ | 6 | 0.0004 $s^{-1}$ |
| U + J ⇄ UJ | 7 | 100,000 $M^{-1}s^{-1}$ | 8 | 1.0 $s^{-1}$ |
| UK + T → UTK | 9 | 90,000 $M^{-1}s^{-1}$ | -- | -- |
| UTK → UDK + P | 10 | 0.0042 $s^{-1}$ | -- | -- |
| UJ + TK ⇄ UTKJ | 11 | 220,000 $M^{-1}s^{-1}$ | 12 | 0.012 $s^{-1}$ |
| UTK + J ⇄ UTKJ | 13 | 220,000 $M^{-1}s^{-1}$ | 14 | 0.012 $s^{-1}$ |
| UTKJ → UDKJ + P | 15 | 1.2 $s^{-1}$ | -- | -- |
| UDKJ → UDK + J | 16 | 1.0 $s^{-1}$ | -- | -- |
| UDKJ + E ⇄ UDKJE | 17 | 500,000 $M^{-1}s^{-1}$ | 18 | 0.11 $s^{-1}$ |
| UDK + E ⇄ UDKE | 19 | 500,000 $M^{-1}s^{-1}$ | 20 | 0.11 $s^{-1}$ |
| UDKJE → UDKE + J | 21 | 1.0 $s^{-1}$ | -- | -- |
| UDKE → UKE + D | 22 | 127 $s^{-1}$ | -- | -- |
| UKE + T → TK + U + E | 23 | 100,000 $M^{-1}s^{-1}$ | -- | -- |
| TK + J ⇄ TKJ | 24 | 220,000 $M^{-1}s^{-1}$ | 25 | 0.012 $s^{-1}$ |
| TK → DK + P | 26 | 0.00075 $s^{-1}$ | -- | -- |
| TKJ → DKJ + P | 27 | 1.2 $s^{-1}$ | -- | -- |
| DKJ → DK + J | 28 | 1.0 $s^{-1}$ | -- | -- |

| | | | | |
|---|---|---|---|---|
| DKJ + E ⇄ DKJE | 29 | 500,000 $M^{-1}s^{-1}$ | 30 | 0.11 $s^{-1}$ |
| DK ⇄ K + D | 31 | 0.022 $s^{-1}$ | 32 | 170000 $M^{-1}s^{-1}$ |
| DK + E ⇄ DKE | 33 | 500,000 $M^{-1}s^{-1}$ | 34 | 0.11 $s^{-1}$ |
| DKJE → DKE + J | 35 | 1.0 $s^{-1}$ | -- | -- |
| DKE → KE + D | 36 | 127 $s^{-1}$ | -- | -- |
| T + K → TK | 37 | 130,000 $M^{-1}s^{-1}$ | 38 | 0.00013 $s^{-1}$ |
| K + E → KE | 39 | 340,000 $M^{-1}s^{-1}$ | 40 | 0.033 $s^{-1}$ |
| KE + T → TK + E | 41 | 150,000 $M^{-1}s^{-1}$ | -- | -- |

**Supplementary Table S6.** Experimentally determined parameters for RNase $H^D$ used exclusively in the CHAMP70 simulations, reproduced from Sekhar *et al.*[1]

| CHAMP70 reaction | CHAMP70 forward rate constant | CHAMP70 reverse rate constant | Experimentally determined forward rate constant for RNase $H^D$ | Experimentally determined reverse rate constant for RNase $H^D$ |
|---|---|---|---|---|
| U + TK ⇄ UTK | 280000 $M^{-1}s^{-1}$ | 3.7 $s^{-1}$ | 280000 $M^{-1}s^{-1}$ | 27 $s^{-1}$ |
| U + DK ⇄ UDK | 1500 $M^{-1}s^{-1}$ | 0.001 $s^{-1}$ | 1800 $M^{-1}s^{-1}$ | 0.0015 $s^{-1}$ |
| U + J ⇄ UJ | 100000 $M^{-1}s^{-1}$ | 1.322 $s^{-1}$ | 100000 $M^{-1}s^{-1}$ | 13.22 $s^{-1}$ |

**Supplementary Table S7**. Rate constants used in the CHAMPION70 model. Forward and reverse rate constants correspond to the kinetic-step numbers shown in Fig. 4 and Supplementary Fig. S3 and S4. For steps that differ across aggregation-propensity regimes, the weakly aggregation-prone (WAP) rate constants were used here. The corresponding rate constants for the non-aggregation-prone (NAP) and strongly-aggregation-prone (SAP) regimes are provided in Supplementary Table S4.

| Kinetic step | Forward step # | Forward rate constant | Reverse step # | Reverse rate constant |
|---|---|---|---|---|
| U ⇄ N | f | protein-dependent | u | protein-dependent |
| U + TK ⇄ UTK | 1 | 280,000 $M^{-1}s^{-1}$ | 2 | 3.7 $s^{-1}$ |
| U + DK ⇄ UDK | 3 | 1,500 $M^{-1}s^{-1}$ | 4 | 0.001 $s^{-1}$ |
| U + K ⇄ UK | 5 | 7,500 $M^{-1}s^{-1}$ | 6 | 0.0004 $s^{-1}$ |
| U + J ⇄ UJ | 7 | 100,000 $M^{-1}s^{-1}$ | 8 | 1.0 $s^{-1}$ |
| UK + T → UTK | 9 | 90,000 $M^{-1}s^{-1}$ | -- | -- |
| UTK → UDK + P | 10 | 0.0042 $s^{-1}$ | -- | -- |
| UJ + TK ⇄ UTKJ | 11 | 216,000 $M^{-1}s^{-1}$ | 12 | 0.012 $s^{-1}$ |
| UTK + J ⇄ UTKJ | 13 | 216,000 $M^{-1}s^{-1}$ | 14 | 0.012 $s^{-1}$ |
| UTKJ → UDKJ + P | 15 | 0.52 $s^{-1}$ | -- | -- |
| UDKJ → UDK + J | 16 | 1.0 $s^{-1}$ | -- | -- |
| UDKJ + E ⇄ UDKJE | 17 | 500,000 $M^{-1}s^{-1}$ | 18 | 0.11 $s^{-1}$ |
| UDK + E ⇄ UDKE | 19 | 500,000 $M^{-1}s^{-1}$ | 20 | 0.11 $s^{-1}$ |
| UDKJE → UDKE + J | 21 | 1.0 $s^{-1}$ | -- | -- |
| UDKE → UKE + D | 22 | 127 $s^{-1}$ | -- | -- |
| UKE + T → TK + U + E | 23 | 100,000 $M^{-1}s^{-1}$ | -- | -- |
| TK + J ⇄ TKJ | 24 | 216,000 $M^{-1}s^{-1}$ | 25 | 0.012 $s^{-1}$ |

| | | | | |
|---|---|---|---|---|
| TK → DK + P | 26 | 0.00075 $s^{-1}$ | -- | -- |
| TKJ → DKJ + P | 27 | 0.52 $s^{-1}$ | -- | -- |
| DKJ → DK + J | 28 | 1.0 $s^{-1}$ | -- | -- |
| DKJ + E ⇄ DKJE | 29 | 500,000 $M^{-1}s^{-1}$ | 30 | 0.11 $s^{-1}$ |
| DK ⇄ K + D | 31 | 0.022 $s^{-1}$ | 32 | 170000 $M^{-1}s^{-1}$ |
| DK + E ⇄ DKE | 33 | 500,000 $M^{-1}s^{-1}$ | 34 | 0.11 $s^{-1}$ |
| DKJE → DKE + J | 35 | 1.0 $s^{-1}$ | -- | -- |
| DKE → KE + D | 36 | 127 $s^{-1}$ | -- | -- |
| T + K → TK | 37 | 130,000 $M^{-1}s^{-1}$ | 38 | 0.00013 $s^{-1}$ |
| K + E → KE | 39 | 340,000 $M^{-1}s^{-1}$ | 40 | 0.033 $s^{-1}$ |
| KE + T → TK + E | 41 | 150,000 $M^{-1}s^{-1}$ | -- | -- |
| U ⇄ M* | 42 | 0.000004 $s^{-1}$ | 43 | $2.6\times10^{-9}$ $s^{-1}$ |
| U’ ⇄ U | 44 | 4,000 $s^{-1}$ | 45 | 0.6 $s^{-1}$ |
| U’ ⇄ M* | 46 | 0.0000048 $s^{-1}$ | 47 | 0.0000011 $s^{-1}$ |
| M* + TK ⇄ M*TK | 48 | 280,000 $M^{-1}s^{-1}$ | 49 | 3.7 $s^{-1}$ |
| M* + DK ⇄ M*DK | 50 | 1,500 $M^{-1}s^{-1}$ | 51 | 0.001 $s^{-1}$ |
| M* + K ⇄ M*K | 52 | 7,500 $M^{-1}s^{-1}$ | 53 | 0.0004 $s^{-1}$ |
| M* + J ⇄ M*J | 54 | 100,000 $M^{-1}s^{-1}$ | 55 | 1 $s^{-1}$ |
| M*K + T → M*TK | 56 | 90,000 $M^{-1}s^{-1}$ | -- | -- |
| M*J + TK ⇄ M*TKJ | 57 | 216,000 $M^{-1}s^{-1}$ | 58 | 0.012 $s^{-1}$ |
| M*TK + J ⇄ M*TKJ | 59 | 216,000 $M^{-1}s^{-1}$ | 60 | 0.012 $s^{-1}$ |
| M*TKJ → U*DKJ + P | 61 | 0.52 $s^{-1}$ | -- | -- |
| U*DKJ → U*DK + J | 62 | 1 $s^{-1}$ | -- | -- |
| U*DKJ + E ⇄ U*DKJE | 63 | 500,000 $M^{-1}s^{-1}$ | 64 | 0.11 $s^{-1}$ |

| | | | | |
|---|---|---|---|---|
| U*DK + E ⇄ U*DKE | 65 | 500,000 $M^{-1}s^{-1}$ | 66 | 0.11 $s^{-1}$ |
| U*DKJE → U*DKE + J | 67 | 1 $s^{-1}$ | -- | -- |
| U*DKE → U*KE + D | 68 | 127 $s^{-1}$ | -- | -- |
| M*TK → U*DK + P | 69 | 0.0042 $s^{-1}$ | -- | -- |
| U*KE + T → TK + N + E | 70 | 100,000 $M^{-1}s^{-1}$ | -- | -- |
| M*DK + E ⇄ M*DKE | 71 | 500,000 $M^{-1}s^{-1}$ | 72 | 0.11 $s^{-1}$ |
| M*DKE → M*KE + D | 73 | 127 $s^{-1}$ | -- | -- |
| M*KE + T → TK + M* + E | 74 | 100,000 $M^{-1}s^{-1}$ | -- | -- |
| 2 * M* ⇄ $A_2$ | 75 | 150 $M^{-1}s^{-1}$ | 76 | $1.92\times10^{-12}$ $s^{-1}$ |
| $A_2$ + M* ⇄ $A_3$ | 77 | $6.06\times10^{9}$ $M^{-1}s^{-1}$ | 78 | $7.82\times10^{-12}$ $s^{-1}$ |
| $A_3$ + M* ⇄ $A_4$ | N/A | $6.06\times10^{9}$ $M^{-1}s^{-1}$ | N/A | $7.82\times10^{-12}$ $s^{-1}$ |
| $A_4$ + M* ⇄ $A_5$ | N/A | $6.06\times10^{9}$ $M^{-1}s^{-1}$ | N/A | $7.82\times10^{-12}$ $s^{-1}$ |
| $A_5$ + M* ⇄ $A_6$ | N/A | $6.06\times10^{9}$ $M^{-1}s^{-1}$ | N/A | $7.82\times10^{-12}$ $s^{-1}$ |
| $A_6$ + M* ⇄ $A_7$ | N/A | $6.06\times10^{9}$ $M^{-1}s^{-1}$ | N/A | $7.82\times10^{-12}$ $s^{-1}$ |
| $A_7$ + M* ⇄ $A_8$ | N/A | $6.06\times10^{9}$ $M^{-1}s^{-1}$ | N/A | $7.82\times10^{-12}$ $s^{-1}$ |
| $A_8$ + M* ⇄ $A_9$ | N/A | $6.06\times10^{9}$ $M^{-1}s^{-1}$ | N/A | $7.82\times10^{-12}$ $s^{-1}$ |
| $A_9$ + M* ⇄ $A_{10}$ | N/A | $6.06\times10^{9}$ $M^{-1}s^{-1}$ | N/A | $7.82\times10^{-12}$ $s^{-1}$ |
| $A_{10}$ + M* ⇄ $A_{11}$ | N/A | $6.06\times10^{9}$ $M^{-1}s^{-1}$ | N/A | $7.82\times10^{-12}$ $s^{-1}$ |
| $A_{11}$ + M* ⇄ $A_{12}$ | N/A | $6.06\times10^{9}$ $M^{-1}s^{-1}$ | N/A | $7.82\times10^{-12}$ $s^{-1}$ |
| $A_{12}$ + M* ⇄ $A_{13}$ | N/A | $6.06\times10^{9}$ $M^{-1}s^{-1}$ | N/A | $7.82\times10^{-12}$ $s^{-1}$ |
| $A_{13}$ + M* ⇄ $A_{14}$ | N/A | $6.06\times10^{9}$ $M^{-1}s^{-1}$ | N/A | $7.82\times10^{-12}$ $s^{-1}$ |
| $A_{14}$ + M* ⇄ $A_{15}$ | N/A | $6.06\times10^{9}$ $M^{-1}s^{-1}$ | N/A | $7.82\times10^{-12}$ $s^{-1}$ |
| $A_{15}$ + M* ⇄ $A_{16}$ | N/A | $6.06\times10^{9}$ $M^{-1}s^{-1}$ | N/A | $7.82\times10^{-12}$ $s^{-1}$ |
| $A_{16}$ + M* ⇄ $A_{17}$ | N/A | $6.06\times10^{9}$ $M^{-1}s^{-1}$ | N/A | $7.82\times10^{-12}$ $s^{-1}$ |

| | | | | |
|---|---|---|---|---|
| $A_{17} + M^* \rightleftarrows A_{18}$ | N/A | $6.06\times10^{9}\ M^{-1}s^{-1}$ | N/A | $7.82\times10^{-12}\ s^{-1}$ |
| $A_{18} + M^* \rightleftarrows A_{19}$ | N/A | $6.06\times10^{9}\ M^{-1}s^{-1}$ | N/A | $7.82\times10^{-12}\ s^{-1}$ |
| $A_{19} + M^* \rightleftarrows A_{20}$ | N/A | $6.06\times10^{9}\ M^{-1}s^{-1}$ | N/A | $7.82\times10^{-12}\ s^{-1}$ |

**Supplementary Table S8.** Kinetic steps and rate constants used in simulations incorporating secondary nucleation.

| Kinetic step | Forward rate constant | Reverse rate constant |
|---|---|---|
| 2 * M* + $A_2 \rightleftarrows A_2 + A_2$ | $6.06\times10^{9}$ $M^{-2}s^{-1}$ | $7.82\times10^{-12}$ $M^{-1}s^{-1}$ |
| 2 * M* + $A_3 \rightleftarrows A_2 + A_3$ | $6.06\times10^{9}$ $M^{-2}s^{-1}$ | $7.82\times10^{-12}$ $M^{-1}s^{-1}$ |
| 2 * M* + $A_4 \rightleftarrows A_2 + A_4$ | $6.06\times10^{9}$ $M^{-2}s^{-1}$ | $7.82\times10^{-12}$ $M^{-1}s^{-1}$ |
| 2 * M* + $A_5 \rightleftarrows A_2 + A_5$ | $6.06\times10^{9}$ $M^{-2}s^{-1}$ | $7.82\times10^{-12}$ $M^{-1}s^{-1}$ |
| 2 * M* + $A_6 \rightleftarrows A_2 + A_6$ | $6.06\times10^{9}$ $M^{-2}s^{-1}$ | $7.82\times10^{-12}$ $M^{-1}s^{-1}$ |
| 2 * M* + $A_7 \rightleftarrows A_2 + A_7$ | $6.06\times10^{9}$ $M^{-2}s^{-1}$ | $7.82\times10^{-12}$ $M^{-1}s^{-1}$ |
| 2 * M* + $A_8 \rightleftarrows A_2 + A_8$ | $6.06\times10^{9}$ $M^{-2}s^{-1}$ | $7.82\times10^{-12}$ $M^{-1}s^{-1}$ |
| 2 * M* + $A_9 \rightleftarrows A_2 + A_9$ | $6.06\times10^{9}$ $M^{-2}s^{-1}$ | $7.82\times10^{-12}$ $M^{-1}s^{-1}$ |
| 2 * M* + $A_{10} \rightleftarrows A_2 + A_{10}$ | $6.06\times10^{9}$ $M^{-2}s^{-1}$ | $7.82\times10^{-12}$ $M^{-1}s^{-1}$ |
| 2 * M* + $A_{11} \rightleftarrows A_2 + A_{11}$ | $6.06\times10^{9}$ $M^{-2}s^{-1}$ | $7.82\times10^{-12}$ $M^{-1}s^{-1}$ |
| 2 * M* + $A_{12} \rightleftarrows A_2 + A_{12}$ | $6.06\times10^{9}$ $M^{-2}s^{-1}$ | $7.82\times10^{-12}$ $M^{-1}s^{-1}$ |
| 2 * M* + $A_{13} \rightleftarrows A_2 + A_{13}$ | $6.06\times10^{9}$ $M^{-2}s^{-1}$ | $7.82\times10^{-12}$ $M^{-1}s^{-1}$ |
| 2 * M* + $A_{14} \rightleftarrows A_2 + A_{14}$ | $6.06\times10^{9}$ $M^{-2}s^{-1}$ | $7.82\times10^{-12}$ $M^{-1}s^{-1}$ |
| 2 * M* + $A_{15} \rightleftarrows A_2 + A_{15}$ | $6.06\times10^{9}$ $M^{-2}s^{-1}$ | $7.82\times10^{-12}$ $M^{-1}s^{-1}$ |
| 2 * M* + $A_{16} \rightleftarrows A_2 + A_{16}$ | $6.06\times10^{9}$ $M^{-2}s^{-1}$ | $7.82\times10^{-12}$ $M^{-1}s^{-1}$ |
| 2 * M* + $A_{17} \rightleftarrows A_2 + A_{17}$ | $6.06\times10^{9}$ $M^{-2}s^{-1}$ | $7.82\times10^{-12}$ $M^{-1}s^{-1}$ |
| 2 * M* + $A_{18} \rightleftarrows A_2 + A_{18}$ | $6.06\times10^{9}$ $M^{-2}s^{-1}$ | $7.82\times10^{-12}$ $M^{-1}s^{-1}$ |
| 2 * M* + $A_{19} \rightleftarrows A_2 + A_{19}$ | $6.06\times10^{9}$ $M^{-2}s^{-1}$ | $7.82\times10^{-12}$ $M^{-1}s^{-1}$ |
| 2 * M* + $A_{20} \rightleftarrows A_2 + A_{20}$ | $6.06\times10^{9}$ $M^{-2}s^{-1}$ | $7.82\times10^{-12}$ $M^{-1}s^{-1}$ |

**Supplementary Table S9.** Kinetic steps and rate constants that differ among the CHAMPION70, hold-only, fold-only and unfold-only kinetic-models. Only the kinetic steps listed here vary across the above models. All other kinetic steps and parameters are identical to those listed in Supplementary Table S7.

| Type of simulation | Kinetic step (pertaining to client-protein release from Hsp70) | Rate constant |
|---|---|---|
| CHAMPION70 | UKE + T → TK + U + E | 100000 $M^{-1}s^{-1}$ |
| | U*KE + T → TK + N + E | 100000 $M^{-1}s^{-1}$ |
| hold-only kinetic-model | UKE + T → TK + U + E | 100000 $M^{-1}s^{-1}$ |
| | U*KE + T → TK + M* + E | 100000 $M^{-1}s^{-1}$ |
| fold-only kinetic-model | UKE + T → TK + N + E | 100000 $M^{-1}s^{-1}$ |
| | U*KE + T → TK + N + E | 100000 $M^{-1}s^{-1}$ |
| unfold-only kinetic-model | UKE + T → TK + U + E | 100000 $M^{-1}s^{-1}$ |
| | U*KE + T → TK + U + E | 100000 $M^{-1}s^{-1}$ |

**Supplementary Table S10.** Kinetic steps used in the simulations modeling asynchronous release of clients from multiple Hsp70 binding sites, corresponding to Fig. S10D. All steps are treated as irreversible, with rate constants of 100,000 $M^{-1}s^{-1}$.

| Number of binding sites | Kinetic step |
|---|---|
| 1 binding site | U*KE + T → N + TK + E |
| 2 binding sites | U*KEU*KE + T → NU*KE + TK + E |
| | U*KEU*KE + T → U*KEN + TK + E |
| | NU*KE + T → NN + TK + E |
| | U*KEN + T → NN + TK + E |
| 3 binding sites | U*KEU*KEU*KE + T → NU*KEU*KE + TK + E |
| | U*KEU*KEU*KE + T → U*KENU*KE + TK + E |
| | U*KEU*KEU*KE + T → U*KEU*KEN + TK + E |
| | NU*KEU*KE + T → NNU*KE + TK + E |
| | NU*KEU*KE + T → NU*KEN + TK + E |
| | U*KENU*KE + T → NNU*KE + TK + E |
| | U*KENU*KE + T → U*KENN + TK + E |
| | U*KEU*KEN + T → NU*KEN + TK + E |
| | U*KEU*KEN + T → U*KENN + TK + E |
| | U*KENN + T → NNN + TK + E |
| | NU*KEN + T → NNN + TK + E |
| | NNU*KE + T → NNN + TK + E |

**Supplementary Table S11.** Main characteristics of the 61 small, single-domain proteins featured in Fig. 5A, 6A and 7A.[14]

| Protein name | Abbreviation | PDB code + ref | Temperature (K) | Length (# of amino acids) | MW (Kda) | $k_f$ ($s^{-1}$) | $k_u$ ($s^{-1}$) | $\Delta G^O$ (kcal/mol) |
|---|---|---|---|---|---|---|---|---|
| Muscle acylphosphatase | mAcP | 1APS[15] | 298.15 | 99 | 10.89 | 0.202 | 0.000123 | -4.36849 |
| SH3 domain (PI3 kinase) | SH3PI3kinase | 1PNJ[16] | 293.15 | 90 | 9.9 | 0.368 | 0.000676 | -3.65675 |
| 9th type III module of fibronectin | fibronectinFN3 | 1FNF[17] | 298.15 | 90 | 9.9 | 0.407 | 0.0550 | -1.18067 |
| 18th module of muscle protein Twitchin | Twitchin | 1WIT[18] | 293.15 | 93 | 10.23 | 1.49 | 0.000275 | -4.99176 |
| Common type acylphosphatase | AcP | 2ACY[19] | 301.15 | 98 | 10.78 | 2.23 | 0.00150 | -4.35282 |
| SH3 domain (alpha-spectrin) | SH3alpha | 1SHG[15] | 298.15 | 62 | 6.82 | 3.00 | 0.00823 | -3.48299 |
| Third fibronectin type III repeat of tenascin | tenascin | 1TEN[20] | 293.15 | 90 | 9.9 | 3.00 | 0.000453 | -5.10785 |
| FK506 Binding Protein | FKBP | 1FKF[15] | 298.15 | 110 | 12.1 | 4.95 | 0.000304 | -5.72627 |

| | | | | | | | | |
|---|---|---|---|---|---|---|---|---|
| Lyme disease variable surface antigen | VSA | 1L8W[15] | 298.15 | 338 | 37.18 | 7.39 | 0.000203 | -6.19854 |
| Ribosomal protein L23 | L23 | 1N88[15] | 298.15 | 96 | 10.56 | 7.39 | 0.0202 | -3.48299 |
| Sho1 SH3 domain | SH3sho1 | NA[15] | 298.15 | 76 | 8.36 | 8.17 | 0.0821 | -2.71555 |
| Actin binding protein ABP1 SH3 domain | SH3ABP | 1JO8[15] | 298.15 | 68 | 7.48 | 12.2 | 0.0672 | -3.06975 |
| Ubiquitin-related modifier 1 | Ubmod | NA[15] | 298.15 | 101 | 11.11 | 13.5 | 0.0369 | -3.48299 |
| Histidine-containing phosphocarrier protein | HPr | 1POH[21] | 293.15 | 85 | 9.35 | 14. | 0.00203 | -5.16589 |
| C-terminal domain from ribosomal protein L9 | CterminalL9 | 1DIV[15] | 298.15 | 92 | 10.12 | 27.1 | 0.000371 | -6.61177 |
| 17$^{th}$ domain of brain alpha-spectrin | 17alpha | 1CUN[22] | 298.15 | 116 | 12.76 | 30.0 | 0.000410 | -6.61177 |
| SH3-like domain of virulence protein internalin B | SH3B | 1M9S[15] | 298.15 | 85 | 9.35 | 54.6 | 0.183 | -3.36492 |

| | | | | | | | | |
|---|---|---|---|---|---|---|---|---|
| Immunoglobulin light chain-binding domain of protein L | proteinL | 2PTL[15] | 298.15 | 79 | 8.69 | 60.3 | 0.0369 | -4.36849 |
| SH3 domain (src) | SH3src | 1RLQ[15] | 298.15 | 61 | 6.71 | 81.5 | 0.273 | -3.36492 |
| Spliceosomal protein U1A | U1A | 1URN[15] | 298.15 | 102 | 11.22 | 99.5 | $8.29\times10^{-6}$ | -9.62249 |
| 16th domain of brain alpha-spectrin | 16alpha | 1CUN[22] | 298.15 | 116 | 12.76 | 122 | 0.00248 | -6.37564 |
| fyn SH3 | fynSH3 | 1AVZ[15] | 298.15 | 78 | 8.58 | 134 | 0.0136 | -5.4311 |
| CspA | CspA | 1MJC[23] | 298.15 | 69 | 7.59 | 200 | 4.06 | -2.30231 |
| Chymotrypsin inhibitor 2 | CI2 | 2CI2[15] | 298.15 | 64 | 7.04 | 330 | 0.000036 | -9.50443 |
| TRF1 Myb domain | TRF1 | 1BA5[24] | 298.15 | 53 | 5.83 | 365 | 3.32 | -2.77458 |
| B1 domain of Streptococcal protein G | proteinG | 3GB1[25] | 295 | 62 | 6.82 | 412 | 0.129 | -4.71369 |
| Ribosomal protein S6 | S6 | 1RIS[15] | 298.15 | 101 | 11.11 | 446 | 0.000249 | -8.50085 |
| Cold shock protein B (Thermatoga maritima) | CspBTm | 1G6P[26] | 298.15 | 66 | 7.26 | 545 | 0.0183 | -6.08047 |
| Cold shock protein B (Bacillus subtilis) | CspBBs | 1CSP[26] | 298.15 | 67 | 7.37 | 665 | 9.97 | -2.47942 |

| | | | | | | | | |
|---|---|---|---|---|---|---|---|---|
| N-terminal domain from ribosomal protein L9 | NterminalL9 | 1DIV[15] | 298.15 | 56 | 6.16 | 735 | 1.11 | -3.83719 |
| Cyclophilin A | CycA | 1LOP[27] | 293.15 | 164 | 18.04 | 735 | 0.0000304 | -9.86743 |
| Activation domain of procarboxypeptidase A2 | ADAh2 | 1O6X[15] | 298.15 | 81 | 8.91 | 898 | 0.670 | -4.25043 |
| Hypothetical protein Tm1083 | Tm1083 | NA[15] | 298.15 | 124 | 13.64 | 992 | 0.00499 | -7.20211 |
| DNA-binding protein sso7D | sso7D | 1C8C[28] | 293 | 64 | 7.04 | 1040 | 0.0392 | -5.91163 |
| Acyl-coenzyme A binding protein | ACBP | 1NTI[15] | 298.15 | 86 | 9.46 | 1100 | 0.0202 | -6.43467 |
| Cold shock protein B (Bacillus caldolyticus) | CspBBc | 1C9O[26] | 298.15 | 66 | 7.26 | 1340 | 0.670 | -4.48656 |
| Colicin E7 immunity protein | ColE7 | 1AYI[15] | 298.15 | 94 | 10.34 | 1340 | 9.97 | -2.89265 |
| Ubiquitin | Ubiquitin | 1UBQ[15] | 298.15 | 76 | 8.36 | 1480 | 0.00111 | -8.32375 |
| Colicin E9 immunity protein | ColE9 | 1IMQ[15] | 298.15 | 93 | 10.23 | 1480 | 0.150 | -5.4311 |

| | | | | | | | | |
|---|---|---|---|---|---|---|---|---|
| Chemotaxis protein CheW | CheW | 1K0S[15] | 298.15 | 151 | 16.61 | 1640 | $5.56\times10^{-6}$ | -11.5116 |
| RAP1 Myb domain | RAP1 | 1FEX[24] | 298.15 | 60 | 6.6 | 3640 | 18.2 | -3.12879 |
| Yes kinase-associated protein | YesK | 1JMQ[29] | 298 | 40 | 4.4 | 4450 | 812 | -1.00307 |
| Ras-binding domain of C-raf-1 | Craf1 | 1RFA[15] | 298.15 | 80 | 8.8 | 4450 | 0.0608 | -6.61177 |
| c-Myb-transforming protein | CMYB | 1GV2[24] | 298.15 | 55 | 6.05 | 6000 | 5.47 | -4.13236 |
| Src SH2 domain | SH2src | 1SPR[15] | 298.15 | 110 | 12.1 | 6000 | 0.0302 | -7.20211 |
| Prototype WW domain | prWW | 1E0M[29] | 298 | 38 | 4.18 | 7330 | 1210 | -1.06207 |
| Hypothetical protein encoded by the Yjbj gene from E coli | Yjbj | 1JYG[15] | 298.15 | 89 | 9.79 | 8960 | 90.0 | -2.71555 |
| Villin headpiece subdomain | villin14T | 1VII[30] | 298.15 | 36 | 3.96 | 12100 | 200 | -2.42038 |
| Pin WW domain | PinWW | 1PIN[31] | 312.65 | 34 | 3.74 | 13400 | 602 | -1.91905 |
| Peripheral subunit-binding domain (PSBD) | PSBD | 2PDD[32] | 298.15 | 41 | 4.51 | 18000 | 221 | -2.59748 |
| lambda-repressor | lambdarep | 1LMB[15] | 298.15 | 80 | 8.8 | 32900 | 24.5 | -4.25043 |

| | | | | | | | | |
|---|---|---|---|---|---|---|---|---|
| Formin binding protein | FBP | 1E0L[29] | 298 | 37 | 4.07 | 40100 | 5430 | -1.18008 |
| 15th domain of brain alpha-spectrin | 15alpha | 1U5P[22] | 298.15 | 116 | 12.76 | 59900 | 1.35 | -6.31661 |
| B domain of staphylococcal protein A | proteinA | 1BDD[33] | 310.15 | 58 | 6.38 | 121000 | 66.7 | -4.60573 |
| BBA5 mini protein | BBA5 | 1T8J[34] | 298 | 23 | 2.53 | 133000 | 540000 | 0.826056 |
| alpha3D | a3D | 2A3D[35] | 298.15 | 73 | 8.03 | 199000 | 2000 | -2.71555 |
| Cyt b562 | cytb562 | 256B[36] | 298.15 | 106 | 11.66 | 220000 | 0.00674 | -10.2128 |
| Trp cage protein | Cage | 1L2Y[37] | 298.15 | 20 | 2.2 | 268000 | 98700 | -0.59034 |
| GA module of albumin binding domain | ABD | 1PRB[38] | 348 | 47 | 5.17 | 985000 | 19900 | -2.68726 |

**Supplementary Table S12.** Individual time intervals used to monitor timecourses in the COPASI simulations of protein folding carried out in the absence and presence of chaperones.

| Figure number(s) | Duration (s) | Number of time intervals | Interval size (s) |
|---|---|---|---|
| Fig. 2E, 3A | 160 | 1600 | 0.1 |
| Fig. 2J, 3C, 3E<br>Supplementary Fig. S1A-B, S6 | 36000 | 3600 | 10 |
| Supplementary Fig. S10D | 1 | 4000 | 0.00025 |
| Fig. 5B-J, 6B-M, 7B-G<br>Supplementary Fig. S13B-J, S14B-G, S15B-G, S16C-N, S17C-K, S18A-L | 0.001 | 2000 | $5\times10^{-7}$ |
| | 0.05 | 50 | 0.001 |
| | 100 | 2000 | 0.05 |
| | 100000 | 1000 | 100 |
| | 100000000 | 1000 | 100000 |
| | 100000000000 | 1000 | 100000000 |
| Supplementary Fig. S7 | 1 | 100 | 0.01 |
| | 120 | 120 | 1 |

**Supplementary Table S13.** Initial chaperone and nucleotide concentrations employed in the kinetic simulations carried out in this work. Note that, if present, ATP, ADP and $P_i$ concentrations were kept constant throughout. Also note that the simulations marked by an asterisk (*), also had initial ADP and $P_i$ concentrations (kept constant) of 0.12 mM and 5 mM, respectively.

| Figure number(s) | Client protein | DnaK (μM) | DnaJ (μM) | GrpE (μM) | ATP (mM) |
|---|---|---|---|---|---|
| Fig. 2C | 5 μM | -- | -- | -- | -- |
| Fig. 2H<br>Supplementary Fig. S1A-B | 0.1 nM, 1 nM, 10 nM, 50 nM, or 200 nM | -- | -- | -- | -- |
| Fig. 3C | 100 nM | 3 | 1 | 1.5 | 5 |
| Fig. 3E | 0.1 nM, 1 nM, 10 nM, 50 nM, or 200 nM | 0.3 | 0.1 | 0.5 | 5 |
| Fig. 5B-J, 6B-M<br>Supplementary Fig. S13B-J, S14B-G, S15B-G, S16C-N, S17C-K, S18A-L | 15 μM | 25 | 5 | 10 | 3.5* |
| Fig. 7B-G | 15 μM | 5, 25, 250 | 5 | 10 | 3.5* |
| Supplementary Fig. S10D | 15 μM | 15 | -- | 15 | 3.5* |
| Fig. 3A | 5 μM | 15 | 3 | 6 | 1 |
| Supplementary Fig. S6 | 100 nM | 3 | 1 | 1.5 | 5 |
| Supplementary Fig. S7 | -- | 1 | 2 | 2 | 0.005 |

| | | | | | |
|---|---|---|---|---|---|
| Supplementary Fig. S8 | -- | 0.5 | 0.5,<br>1, 2,<br>3, 5,<br>8, 15 | -- | 0.008 |

**Supplementary Table S14.** Comparisons between experimentally determined and computationally predicted peptide-binding rate constants for various chaperone combinations within the K/J/E cycle.

| Peptide-binding process | Experimental[39] ($s^{-1}$) | Predicted ($s^{-1}$) |
|---|---|---|
| Peptide + [DnaK + ATP] | 2.3 ± 0.2[a] — 6.8 ± 0.7[b] | 4.0 |
| Peptide + [DnaK + DnaJ + ATP] | 0.0200 ± 0.0002[a] | 0.0059 |
| Peptide + [DnaK + ADP + $P_i$] | 0.04 ± 0.02[a] | 0.05 |
| Peptide + [DnaK + GrpE + ATP] | 5.7 ± 0.2[a] | 3.9 |

[a] Experimental peptide binding rate constant determined with a-pp peptide (acrylodan-labeled 23-residue mitochondrial aspartate aminotransferase presequence peptide [pp]).
[b] Experimental peptide binding rate constant determined with a-p5 peptide (acrylodan-labeled 9-residue peptide corresponding to residues 8-15 of mitochondrial aspartate aminotransferase presequence).

**Supplementary Table S15.** Comparisons between experimentally determined and computationally predicted stopped-flow rate constants for the interaction of RNase $H^D$ with the K/J/E chaperone system.

| Species in Refolding Mixture | Experimental[40] ($s^{-1}$) | CHAMPION70 Predicted ($s^{-1}$) | Chi-Square |
|---|---|---|---|
| RNase $H^D$ | 0.034 ± 0.002 | 0.034 | $1.4\times10^{-20}$ |
| RNase $H^D$, DnaJ | 0.029 ± 0.003 | 0.028 | $1.9\times10^{-13}$ |
| RNase $H^D$, ATP-DnaK | 0.029 ± 0.009 | 0.020 | $1.4\times10^{-13}$ |
| RNase $H^D$, ATP-DnaK, DnaJ | 0.066 ± 0.002 | 0.066 | $1.2\times10^{-12}$ |
| RNase $H^D$, ATP-DnaK, DnaJ, GrpE | 0.0290 ± 0.0006 | 0.0170 | $3.1\times10^{-13}$ |

**Supplementary Table S16.** Comparison of experimentally determined human Hsp70A1 rate constants with the corresponding DnaK rate constants used in the CHAMPION70 model. Human Hsp70A1 kinetic values were obtained by Wu *et al.*,[41] where single-molecule FRET and kinetic analysis were used to quantify ATP binding, ATP hydrolysis, and ATP-induced conformational transitions in the stress-inducible Hsp70A1 chaperone.

| Kinetic step | Human Hsp70A1 rate constant | CHAMPION70 (*E. Coli* DnaK) rate constant |
|---|---|---|
| ATP + Hsp70 → ATP·Hsp70 (T + K → TK) | (3.9–6.1) × $10^5$ $M^{-1}s^{-1}$ | 1.3 × $10^5$ $M^{-1}s^{-1}$ |
| ATP·Hsp70 → ATP + Hsp70 (TK → T + K) | 0.05–0.43 $s^{-1}$ | 1.3 × $10^{-4}$ $s^{-1}$ |
| ATP·Hsp70 → ADP·Hsp70 + $P_i$ (TK → DK + P) | 0.0013 $s^{-1}$ | 7.5 × $10^{-4}$ $s^{-1}$ |
| ATP·Hsp70·Hsp40 → ADP·Hsp70·Hsp40 + $P_i$ (TKJ → DKJ + P) | 0.0067 $s^{-1}$ | 0.52 $s^{-1}$ |
| ATP·Hsp70 + Hsp40 → ATP·Hsp70·Hsp40 (TK + J → TKJ) | 3.5 × $10^5$ $M^{-1}s^{-1}$ | 2.2 × $10^5$ $M^{-1}s^{-1}$ |

### Supplementary Figures

#### Fig. S1

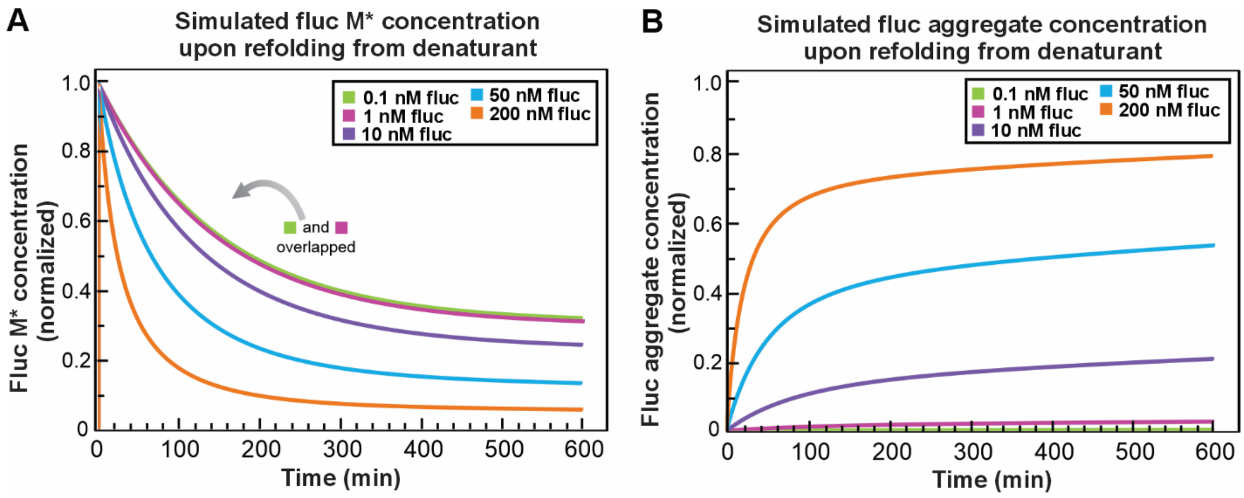


**Supplementary Fig. S1. Simulated time-dependent M* and $A_{20}$ concentrations during fluc refolding in the absence of chaperones. A.** Simulated concentration-dependent fluc (0.1-200 nM) refolding over 600 minutes in the absence of chaperones, showing the time-dependent behavior of the misfolded intermediate (M*). **B.** Simulated concentration-dependent fluc (0.1-200 nM) refolding over 600 minutes in the absence of chaperones, showing the timecourse of formation of the final aggregate ($A_{20}$). Experimental conditions for these kinetic simulations are according to the literature.[42]

**Fig. S2**

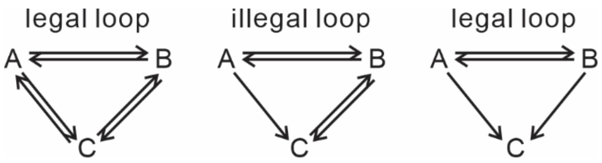


**Supplementary Fig. S2. Examples of legal and illegal loops in a reaction network.** Legal loops correspond to pathways that satisfy the thermodynamic requirement of detailed balance, while illegal loops correspond to pathways that violate this principle.[7]

### Fig. S3

#### Hold-only kinetic-model (non-fold-promoting scheme)

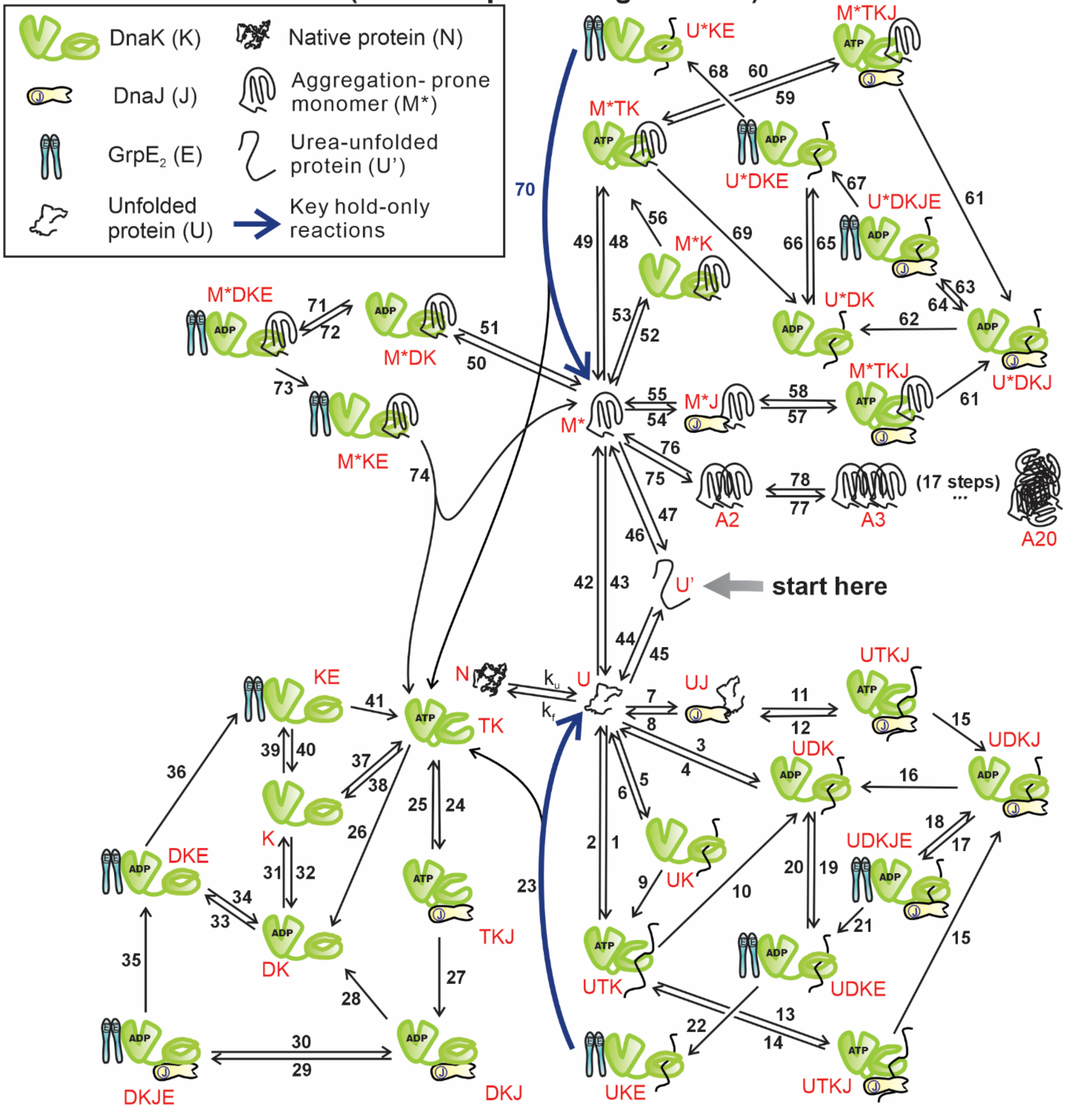


DK: (ADP)DnaK
TK: (ATP)DnaK
TKJ: (ATP)DnaK-DnaJ
DKJ: (ADP)DnaK-DnaJ
M*J: M*-DnaJ
M*K: M*-DnaK
M*KE: M*-DnaK-GrpE$_2$

KE: DnaK-GrpE$_2$
DKJE: (ADP)DnaK-DnaJ-GrpE$_2$
DKE: (ADP)DnaK-GrpE$_2$
UKE: U-DnaK-GrpE$_2$
M*TKJ: M*-(ATP)DnaK-DnaJ
M*DKJ: M*-(ADP)DnaK-DnaJ

UJ: U-DnaJ
UK: U-DnaK
UDK: U-(ADP)DnaK
UTK: U-(ATP)DnaK
M*DK: M*-(ADP)DnaK
UTK: U-(ATP)DnaK

UTKJ: U-(ATP)DnaK-DnaJ
UDKJ: U-(ADP)DnaK-DnaJ
UDKJE: U-DnaJ-(ADP)DnaK-GrpE$_2$
UDKE: U-(ADP)DnaK-GrpE$_2$
M*DKJE: M*-DnaJ-(ADP)DnaK-GrpE$_2$
M*DKE: M*-(ADP)DnaK-GrpE$_2$

**Supplementary Fig. S3. Reaction scheme for the hold-only kinetic-model.** According to this hold-only kinetic model (non-fold-promoting scheme), unfolded and misfolded client-proteins are released from DnaK in the same conformation as the one assumed upon initial interaction with the chaperone. The key hold-only processes that distinguish this kinetic model from the fold-only, unfold-only, and CHAMPION70 kinetic models are highlighted in blue. Independent nucleotides involved in some of the steps are not explicitly shown, for simplicity.

**Fig. S4**

### Fold-only kinetic-model (fold-promoting scheme)

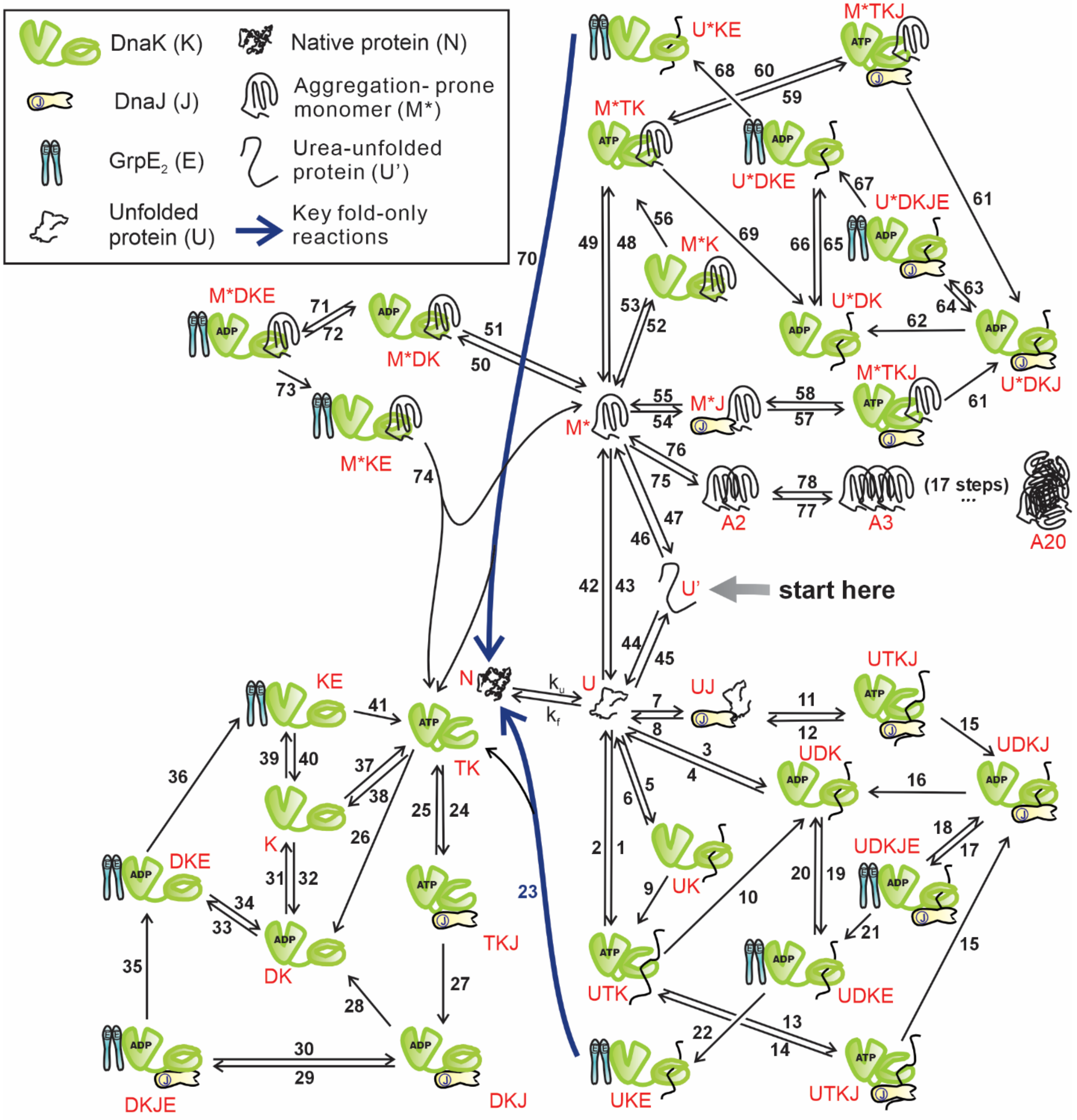


DK: (ADP)DnaK
TK: (ATP)DnaK
TKJ: (ATP)DnaK-DnaJ
DKJ: (ADP)DnaK-DnaJ
M*J: M*-DnaJ
M*K: M*-DnaK
M*KE: M*-DnaK-GrpE$_2$

KE: DnaK-GrpE$_2$
DKJE: (ADP)DnaK-DnaJ-GrpE$_2$
DKE: (ADP)DnaK-GrpE$_2$
UKE: U-DnaK-GrpE$_2$
M*TKJ: M*-(ATP)DnaK-DnaJ
M*DKJ: M*-(ADP)DnaK-DnaJ

UJ: U-DnaJ
UK: U-DnaK
UDK: U-(ADP)DnaK
UTK: U-(ATP)DnaK
M*DK: M*-(ADP)DnaK
UTK: U-(ATP)DnaK

UTKJ: U-(ATP)DnaK-DnaJ
UDKJ: U-(ADP)DnaK-DnaJ
UDKJE: U-DnaJ-(ADP)DnaK-GrpE$_2$
UDKE: U-(ADP)DnaK-GrpE$_2$
M*DKJE: M*-DnaJ-(ADP)DnaK-GrpE$_2$
M*DKE: M*-(ADP)DnaK-GrpE$_2$

**Supplementary Fig. S4. Reaction scheme for the fold-only kinetic-model.** According to this fold-only kinetic model (fold-promoting scheme), unfolded and misfolded client-proteins are released from DnaK in their native conformation. The key fold-only processes that distinguish this kinetic model from the hold-only, unfold-only, and CHAMPION70 models are highlighted in blue. Independent nucleotides involved in some of the steps are not explicitly shown, for simplicity.

### Fig. S5

#### Unfold-only kinetic-model

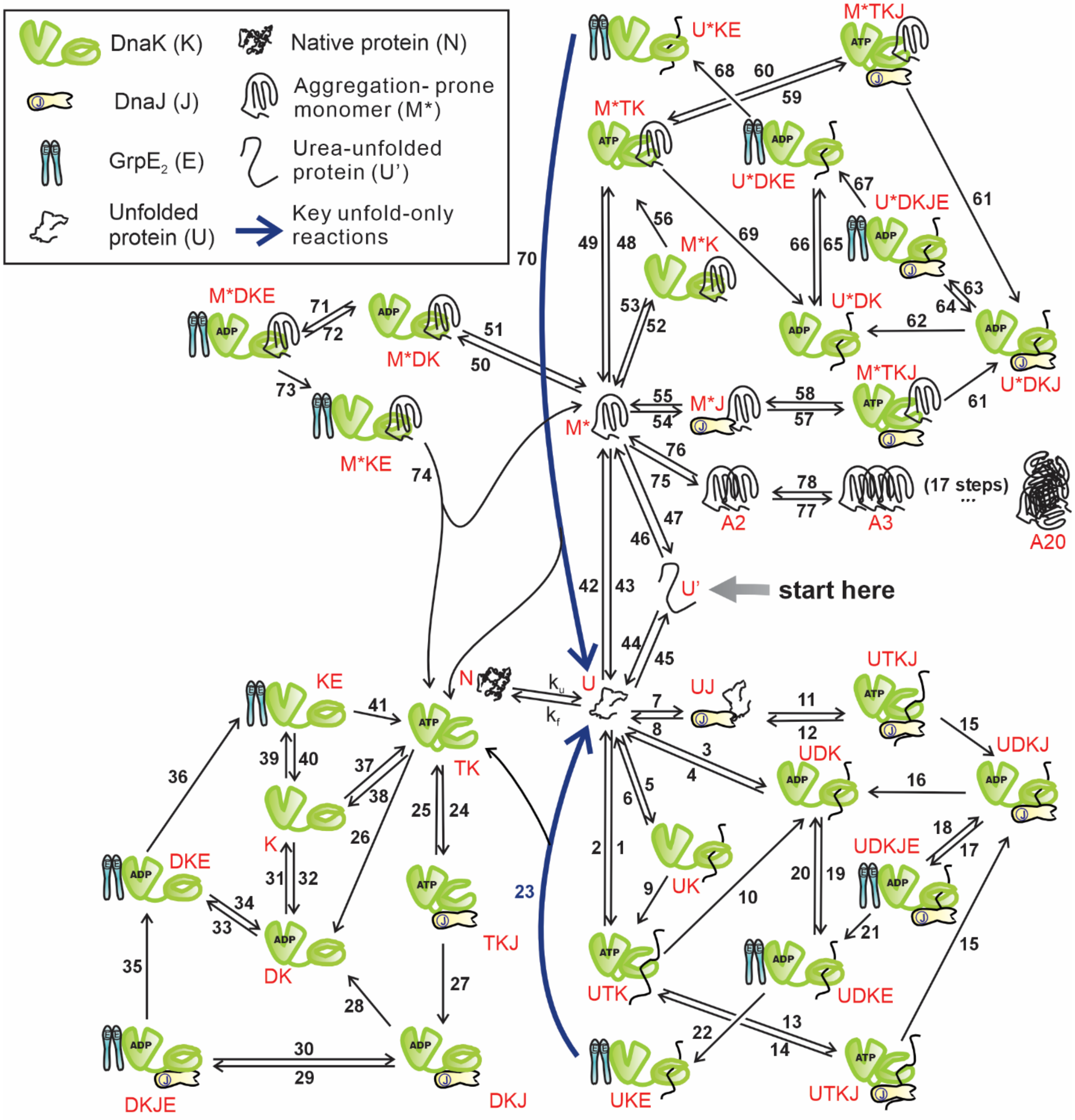


DK: (ADP)DnaK
TK: (ATP)DnaK
TKJ: (ATP)DnaK-DnaJ
DKJ: (ADP)DnaK-DnaJ
M*J: M*-DnaJ
M*K: M*-DnaK
M*KE: M*-DnaK-GrpE$_2$

KE: DnaK-GrpE$_2$
DKJE: (ADP)DnaK-DnaJ-GrpE$_2$
DKE: (ADP)DnaK-GrpE$_2$
UKE: U-DnaK-GrpE$_2$
M*TKJ: M*-(ATP)DnaK-DnaJ
M*DKJ: M*-(ADP)DnaK-DnaJ

UJ: U-DnaJ
UK: U-DnaK
UDK: U-(ADP)DnaK
UTK: U-(ATP)DnaK
M*DK: M*-(ADP)DnaK
UTK: U-(ATP)DnaK

UTKJ: U-(ATP)DnaK-DnaJ
UDKJ: U-(ADP)DnaK-DnaJ
UDKJE: U-DnaJ-(ADP)DnaK-GrpE$_2$
UDKE: U-(ADP)DnaK-GrpE$_2$
M*DKJE: M*-DnaJ-(ADP)DnaK-GrpE$_2$
M*DKE: M*-(ADP)DnaK-GrpE$_2$

**Supplementary Fig. S5. Reaction scheme for the unfold-only kinetic-model.** According to this unfold-only kinetic model, unfolded and misfolded client-proteins are released from DnaK in their unfolded conformation. The key unfold-only processes that distinguish this kinetic model from the hold-only, fold-only, and CHAMPION70 models are highlighted in blue. Independent nucleotides involved in some of the steps are not explicitly shown, for simplicity.

**Fig. S6**

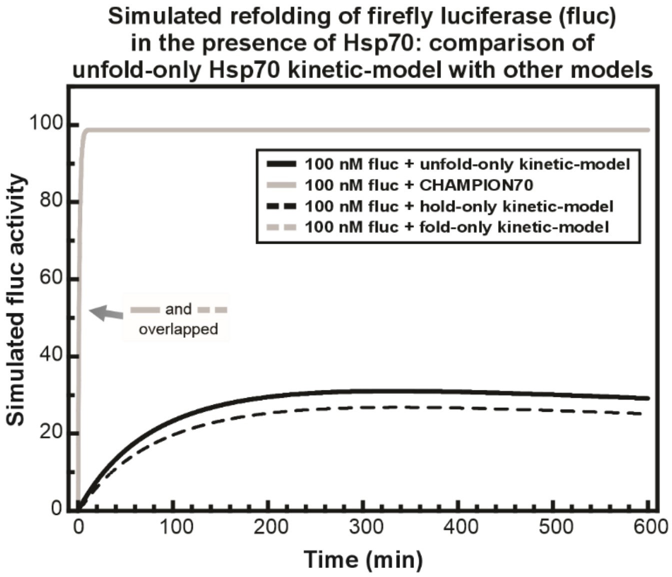


**Supplementary Fig. S6. Computational evaluation of unfold-only kinetic-model in the context of fluc refolding.** According to the unfold-only kinetic-model, the misfolded intermediate M* is converted to the unfolded state U within its Hsp70-bound state. and then released as U. In contrast, the fold-only kinetic model includes conversion of M* to some unfolded species denoted as U* within the chaperone-bound state. This species is then released as N. The unfold-only kinetic model is also different from the hold-only model, where M* does not undergo any conformational changes while chaperone-bound and is released from the chaperone as M*. As shown in the graph, conversion of M* to U while Hsp70-bound (within the unfold-only model) is not sufficient to reproduce the experimental fluc refolding behavior observed in Fig. 3B of the main article. In contrast, the CHAMPION70 and fold-only kinetic models are both able to reproduce the experimental results. However, as discussed in the main article, only the CHAMPION70 (but not the fold-only) model is also able to reproduce the experimental data pertaining to the interaction of RNase $H^D$ with Hsp70, which is a non-aggregation-prone (NAP) protein.

**Fig. S7**

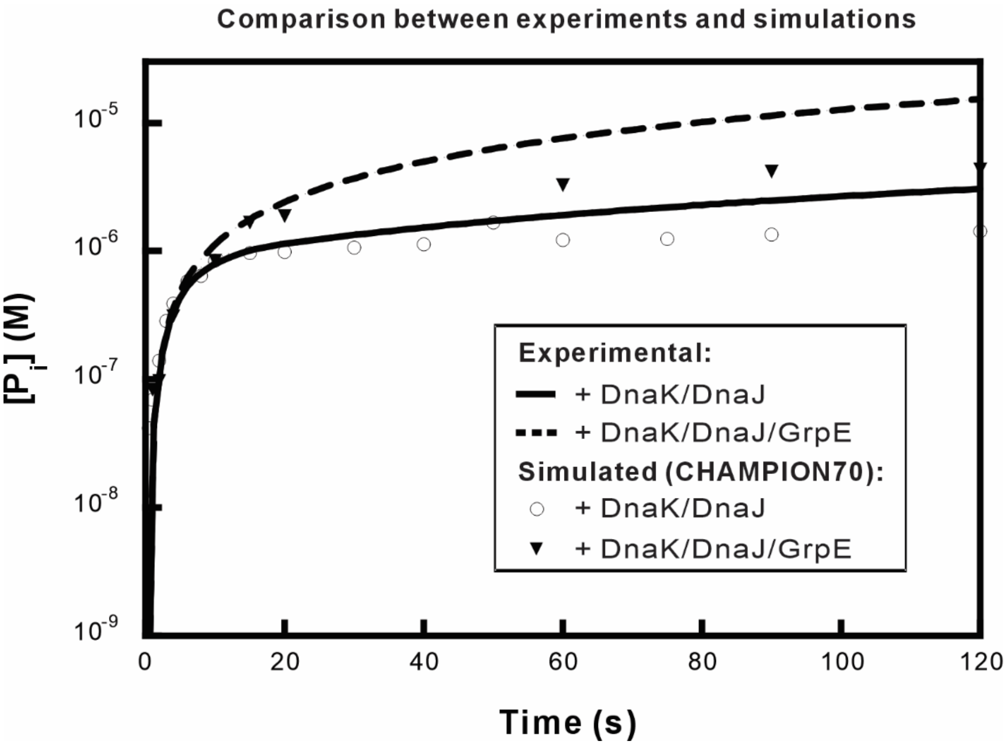


**Supplementary Fig. S7. Comparison between experimental and simulated ATP hydrolysis timecourses.** Comparison between experimentally observed[43] and computationally predicted timecourses of DnaK-catalyzed ATP hydrolysis in the presence of DnaK and DnaJ (solid line and circles) or DnaK, DnaJ, and GrpE (dashed line and triangles). Experimental multiple-turnover ATPase activity was measured by rapid-quench experiments. Experimental samples consisted of 1 μM DnaK, 5 μM ATP, and 2 μM DnaJ (circles) or 1 μM DnaK, 5 μM ATP, 2 μM DnaJ, and 2 μM GrpE (triangles). These concentrations were used as initial concentrations, in all the computer simulations.

**Fig. S8**

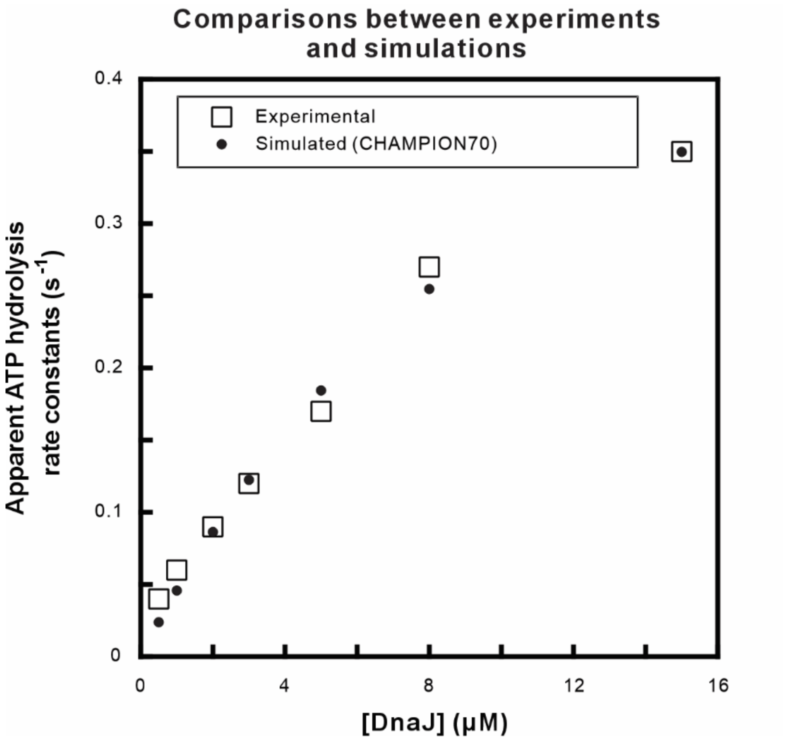


**Supplementary Fig. S8. Comparison between computations and experiments focusing on the DnaJ-dependent stimulation of DnaK ATPase activity in the absence of client-protein.** Experimentally observed[4] and computationally predicted DnaJ-dependent stimulation of DnaK ATP hydrolysis in the absence of a client-protein. Experimental hydrolysis rates were determined by quench-flow experiments. Experimental samples consisted of 500 nM DnaK and 8 μM ATP, which were preequilibrated for 10 s before mixing with varying concentrations of DnaJ. The same procedure was employed in the computer simulations by preequilibrating DnaK and ATP (kept at constant concentration) for 10 s and then using the resulting equilibrated DnaK concentration as the initial condition, together with the specified DnaJ concentration. The rate constant for ATP hydrolysis in the presence of DnaJ (reaction #27) was optimized to fit this experimental data.

**Fig. S9**

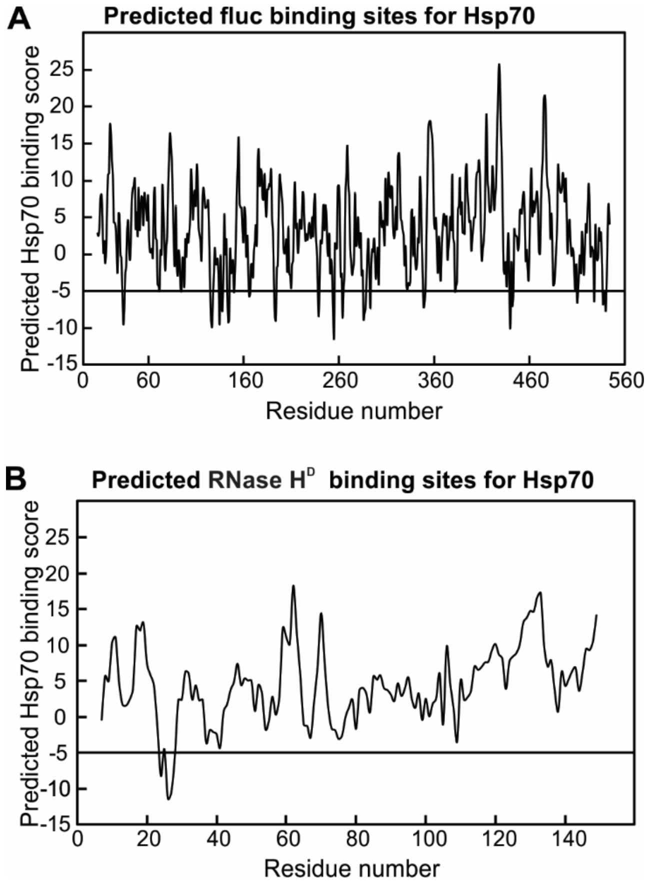


**Supplementary Fig. S9. Computationally predicted Hsp70-binding sites of fluc and RNase H$^{D}$.** To carry out this prediction, we employed the sequence-based algorithm developed by Rüdiger *et al.* (1997),[44] using a cutoff binding score of ≤ -5, to identify Hsp70 binding sites. **A**. Firefly luciferase (fluc) is predicted to have ca. 17 Hsp70-binding sites. **B.** RNase H$^{D}$ is predicted to have a single Hsp70-binding site.

**Fig. S10**

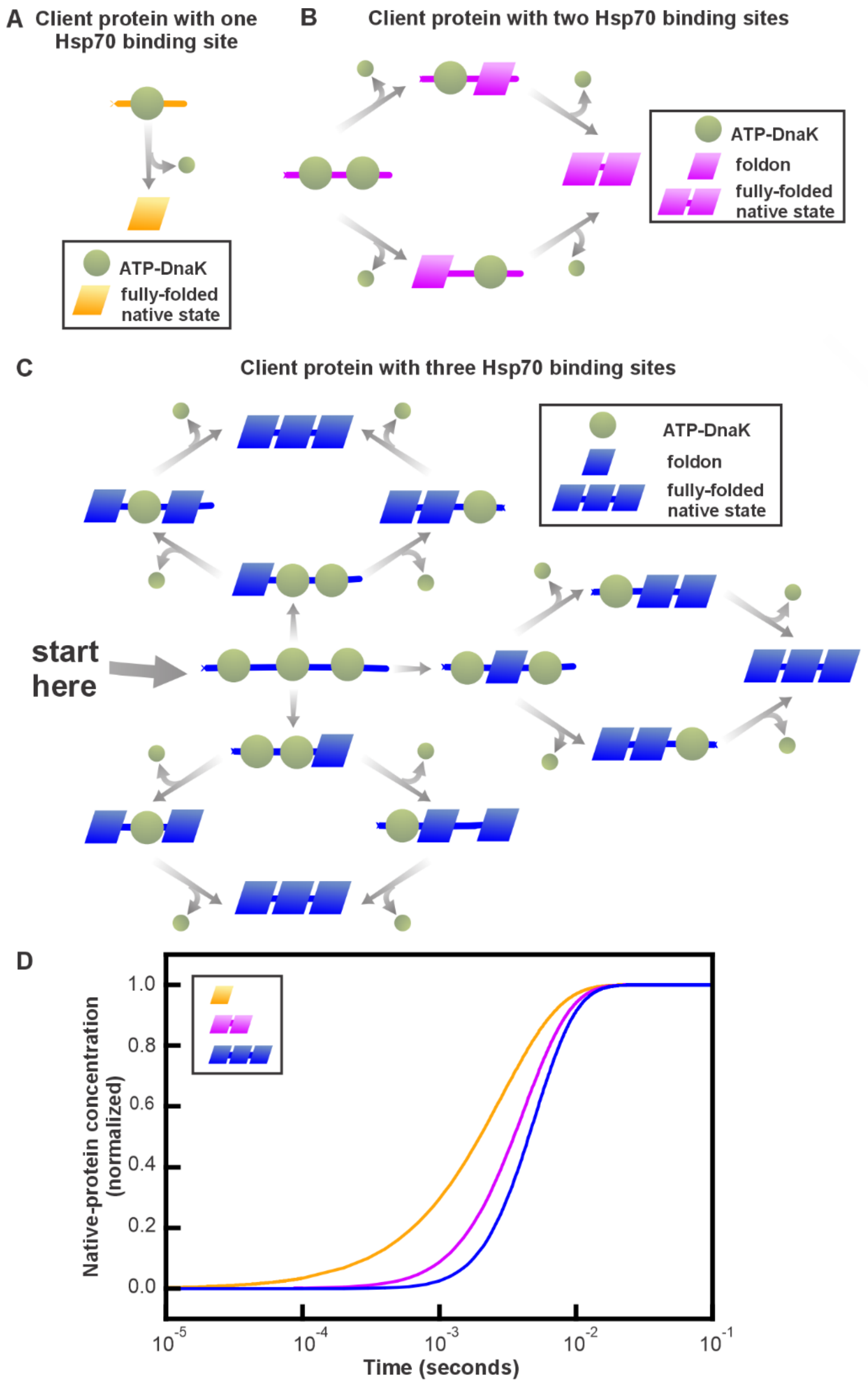

A Client protein with one Hsp70 binding site
ATP-DnaK
fully-folded native state
B Client protein with two Hsp70 binding sites
ATP-DnaK
foldon
fully-folded native state
C Client protein with three Hsp70 binding sites
ATP-DnaK
foldon
fully-folded native state
start here
D
Native-protein concentration (normalized)
1.0
0.8
0.6
0.4
0.2
0.0
10^-5
10^-4
10^-3
10^-2
10^-1
Time (seconds)

**Fig. S10. The presence of multiple chaperone binding sites leads to a slower release of client proteins from the chaperone. A.** Kinetic scheme for client-protein released from a chaperone in a single step. **B.** Kinetic scheme for the asynchronous chaperone release of a client-protein bearing two chaperone-binding sites. **C.** Kinetic scheme for asynchronous release of a client-protein bearing three chaperone-binding sites. **D.** Simulations illustrating the asynchronous release from the chaperone of client-proteins bearing 1, 2, or 3 chaperone-binding sites. The computer simulations included the following initial concentrations: 15 μM client-protein bound to DnaK and GrpE in the presence of 3.5 mM ATP, 0.12 mM ADP, and 5 mM $P_i$. All the specific kinetic steps are in Supplementary Table S10.

**Fig. S11**

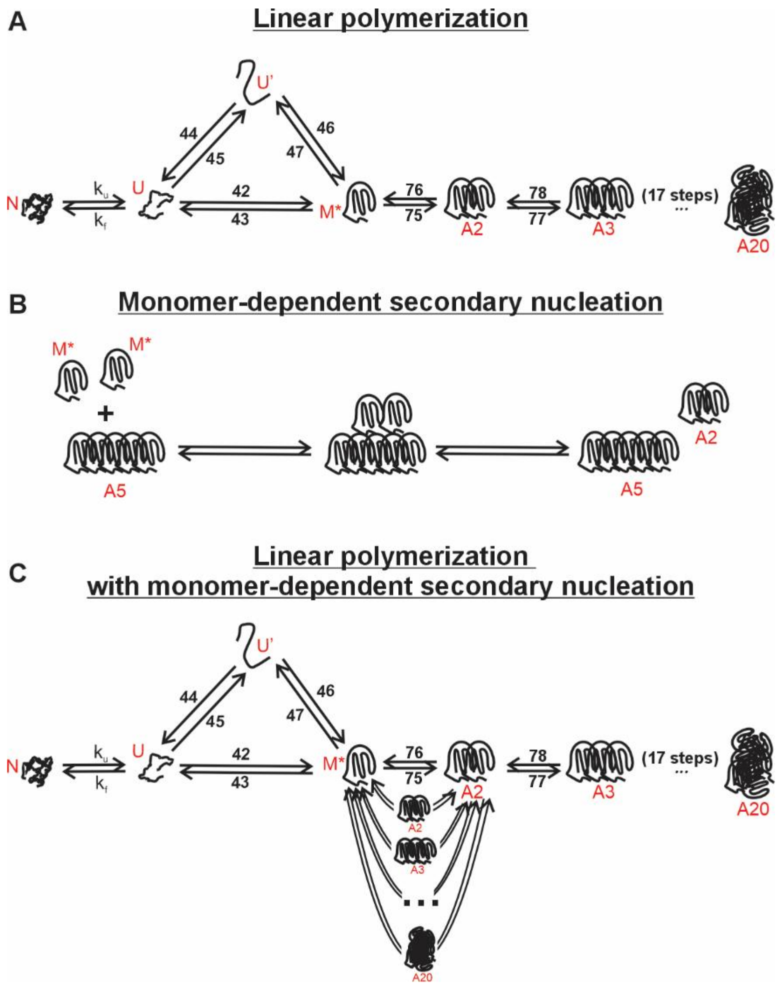


**Supplementary Fig. S11. Schematic illustration of the aggregation pathways evaluated in this study. A.** Linear polymerization pathway, according to which misfolded monomers (M*) undergo sequential elongation to generate growing aggregates ($A_2$–$A_{20}$). **B.** Monomer-dependent secondary nucleation pathway, according to which aggregated surfaces catalyze the formation of new $A_2$ nuclei. **C.** Secondary nucleation pathway, which illustrates how aggregated surfaces bypass the rate-limiting primary nucleation step to accelerate aggregation.

**Fig. S12**

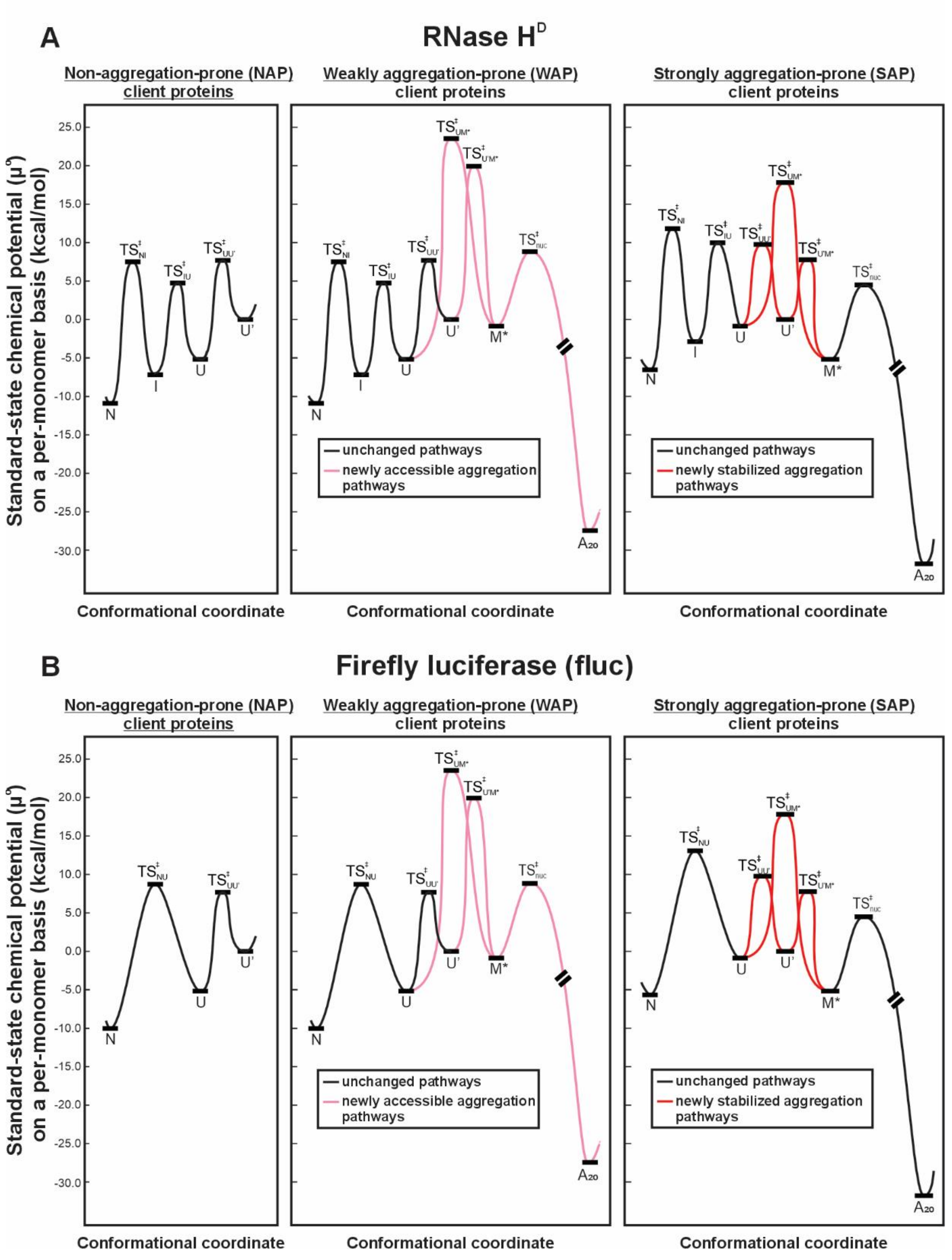


**Supplementary Fig. S12. Standard-state chemical potential diagrams for RNase H$^{D}$ and fluc under different aggregation-propensity regimes. A.** Standard-state chemical potential diagrams of RNase $H^D$ under NAP, WAP, and SAP aggregation-propensity conditions. **B.** Corresponding diagrams for fluc under the same aggregation-propensity regimes. Under WAP conditions, misfolding and aggregation pathways (pink) are included. Under SAP conditions, the kinetic barriers leading to misfolding (U' ⇄ U, U' ⇄ M*, U ⇄ M*) are lower, providing a more kinetically accessible path to misfolding and subsequent aggregation. In addition, under SAP aggregation-propensity conditions, U is destabilized leading to raising the free energy of the folding pathway. Moreover, M* is stabilized leading to a lowering of the free energy of the nucleation and elongation aggregation pathways.

### Fig. S13

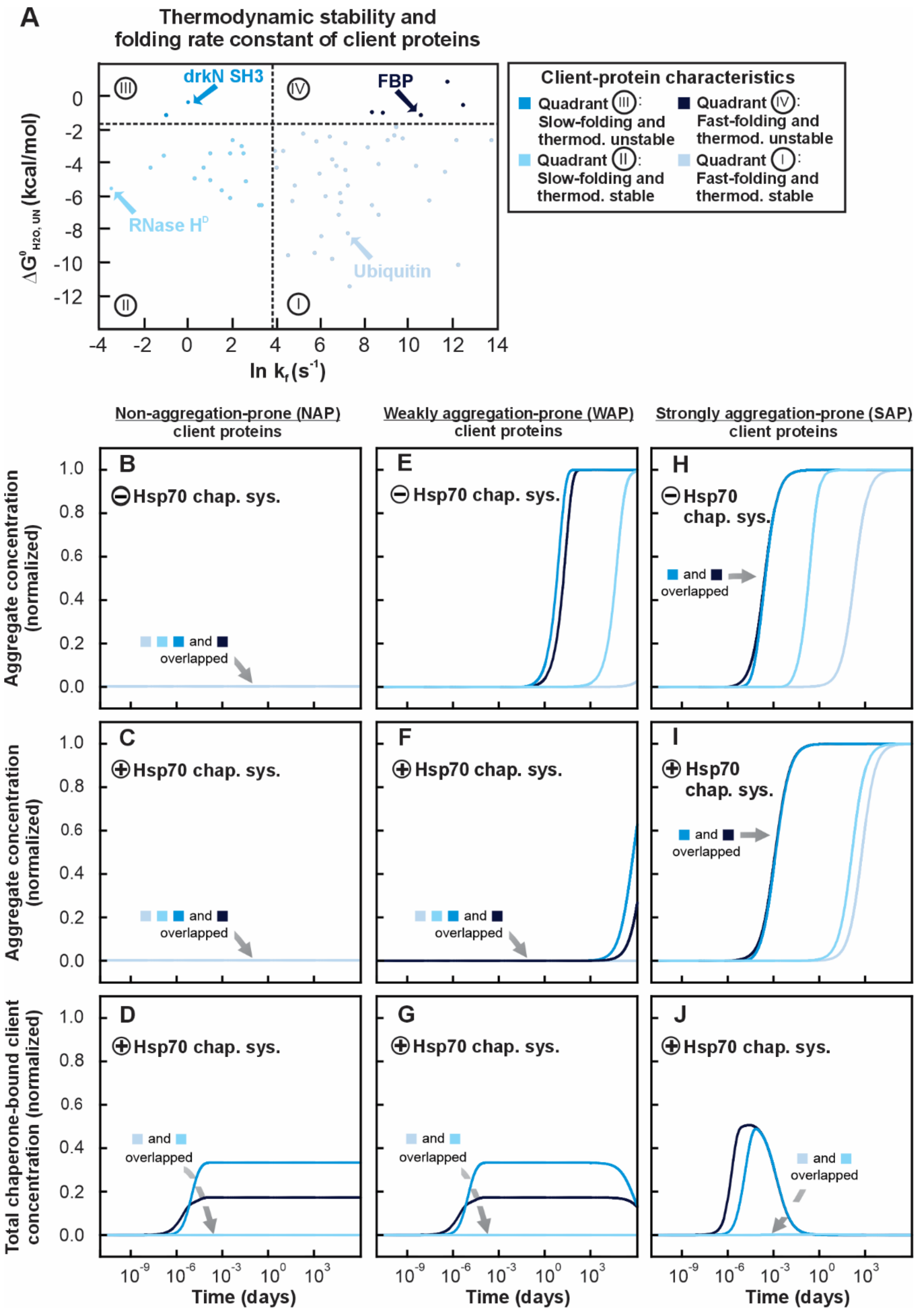

A
Thermodynamic stability and folding rate constant of client proteins
drkN SH3
FBP
RNase H[D]
Ubiquitin
ln $k_f$ ($s^{-1}$)
$\Delta G^0_{H2O, UN}$ (kcal/mol)
Client-protein characteristics
Quadrant III: Slow-folding and thermod. unstable
Quadrant IV: Fast-folding and thermod. unstable
Quadrant II: Slow-folding and thermod. stable
Quadrant I: Fast-folding and thermod. stable
Non-aggregation-prone (NAP) client proteins
Weakly aggregation-prone (WAP) client proteins
Strongly aggregation-prone (SAP) client proteins
Aggregate concentration (normalized)
Total chaperone-bound client concentration (normalized)
B
E
H
C
F
I
D
G
J
Hsp70 chap. sys.
and overlapped
Time (days)

**Supplementary Fig. S13. Computationally simulated aggregation and chaperone-binding dynamics of representative client-proteins initiated from the native-state. A.** Diagram illustrating the thermodynamic and kinetic characteristics of 61 small client proteins sorted according to their known thermodynamic stability and folding rate constants.[14] Four proteins were selected (various shades of blue) as representative chaperone clients for each of the four quadrants. All the CHAMPION70 simulations began with 15 µM client-protein in the native-state (N), in contrast to Fig. 5 where simulations began from the U' state. Simulations in the presence of chaperones included 25 µM DnaK, 5 µM DnaJ, 10 µM GrpE, 3.5 mM ATP, 0.12 mM ADP, and 5 mM $P_i$. **B, E, H.** Simulated formation of fully aggregated $A_{20}$ species in the absence of chaperones for clients with three aggregation propensities: non-aggregation-prone (NAP) (**B**), weakly aggregation-prone (WAP) (**E**), and strongly aggregation-prone (SAP) (**H**), shown for each of the four representative clients from each panel-A region. **C, F, I.** Simulated formation of $A_{20}$ in the presence of chaperones for non-aggregation-prone (NAP) (**C**), weakly aggregation-prone (WAP) (**F**), and strongly aggregation-prone (SAP) (**I**) client-proteins. **D, G, J.** Simulated association of clients within the Hsp70 cycle for non-aggregation-prone (NAP) (**D**), weakly aggregation-prone (WAP) (**G**), and strongly aggregation-prone (SAP) (**J**) client proteins.

### Fig. S14

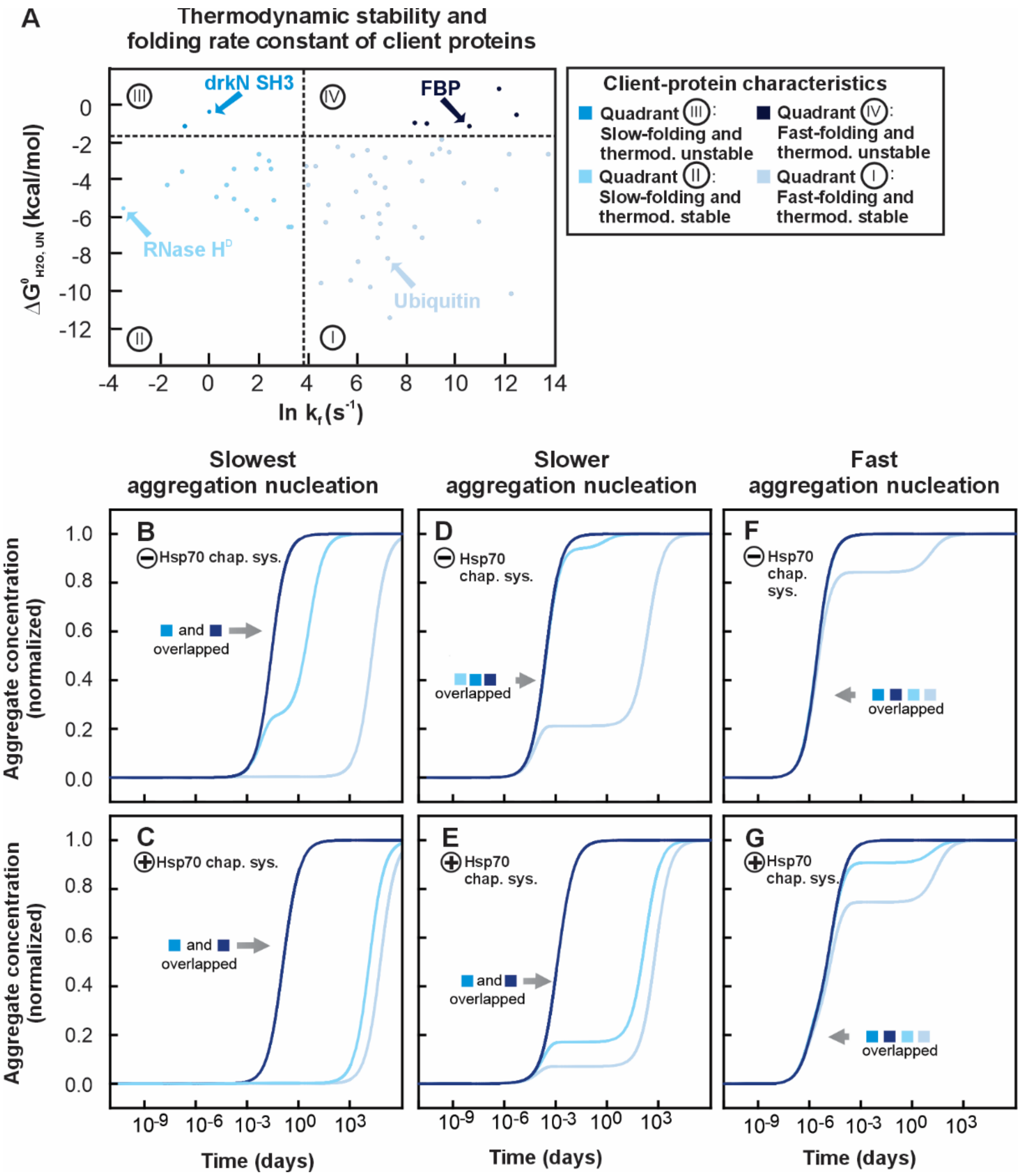


**Supplementary Fig. S14. Computationally predicted influence of aggregation nucleation rates on client-protein folding and aggregation in the absence and presence of the Hsp70 chaperone system. A.** Diagram illustrating the thermodynamic and kinetic characteristics of 61 small client proteins sorted according to their known thermodynamic stability and folding rate constants.[14] Four proteins were selected (various shades of blue) as representative chaperone clients for each of the four quadrants. All the CHAMPION70 simulations began with 15 μM client-protein in the conformationally unrelaxed denatured unfolded state (U'). Simulations in the presence of chaperones included 25 μM DnaK, 5 μM DnaJ, 10 μM GrpE, 3.5 mM ATP, 0.12 mM ADP, and 5 mM $P_i$. Formation of the fully aggregated state ($A_{20}$) was simulated using strongly aggregation-prone (SAP) rate constants. **B, D, F.** Simulated formation of fully aggregated $A_{20}$ species in the absence of chaperones for clients with nucleation rates that are two-fold slower than the CHAMPION70 value (reaction #75 and #76) (**B**), equal to the CHAMPION70 value (**D**), and two-fold faster than the CHAMPION70 value (**F**). **C, E, G.** Simulated formation of $A_{20}$ in the presence of chaperones for nucleation rates that are two-fold slower than the CHAMPION70 value (**C**), equal to the CHAMPION70 value (**E**), and two-fold slower than the CHAMPION70 value (**G**).

**Fig. S15**

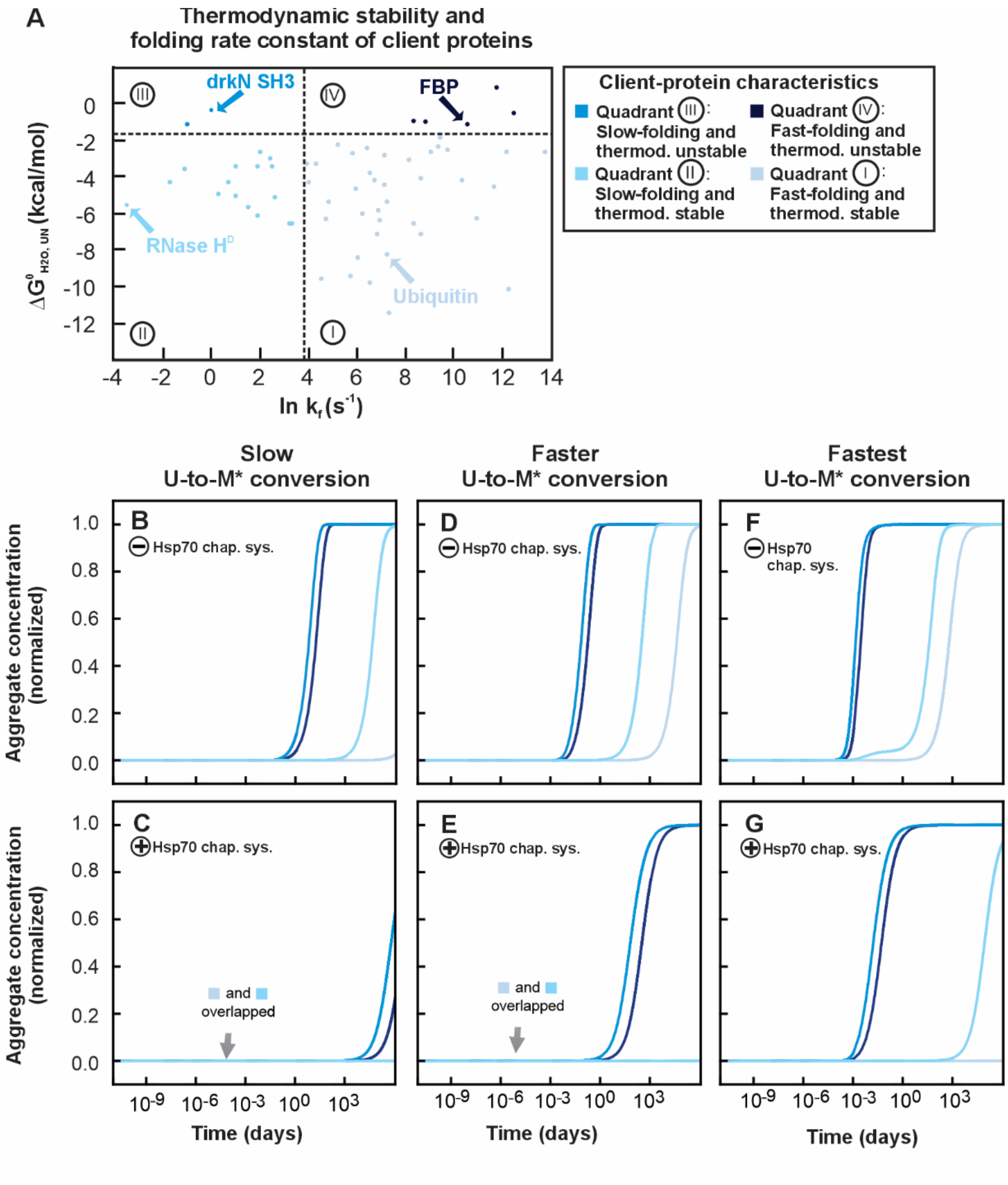


**Supplementary Fig. S15. Computationally predicted influence U to M* barrier height in the absence and presence of the Hsp70 chaperone system. A.** Diagram illustrating the thermodynamic and kinetic characteristics of 61 small client proteins sorted according to their known thermodynamic stability and folding rate constants.[14] Four proteins were selected (various shades of blue) as representative chaperone clients for each of the four quadrants. All the

CHAMPION70 simulations began with 15 μM client-protein in the conformationally unrelaxed denatured unfolded state (U’). Simulations in the presence of chaperones included 25 μM DnaK, 5 μM DnaJ, 10 μM GrpE, 3.5 mM ATP, 0.12 mM ADP, and 5 mM $P_i$. Formation of the fully aggregated state ($A_{20}$) was simulated using weakly aggregation-prone (WAP) rate constants. **B, D, F.** Simulated formation of fully aggregated $A_{20}$ species in the absence of chaperones for clients with U to M* rate constants that are equal to the CHAMPION70 rates (reaction #42 and #43) (**B**), two-fold faster than the CHAMPION70 rates (**D**), and four-fold faster than the CHAMPION70 rates (**F**), shown for each of the four representative clients from each panel-A region. **C, E, G.** Simulated formation of $A_{20}$ in the presence of chaperones for clients with U to M* rate constants that are equal to the CHAMPION70 rates (**C**), two-fold faster than the CHAMPION70 rates (**E**), and four-fold faster than the CHAMPION70 rates (**G**).

### Fig. S16

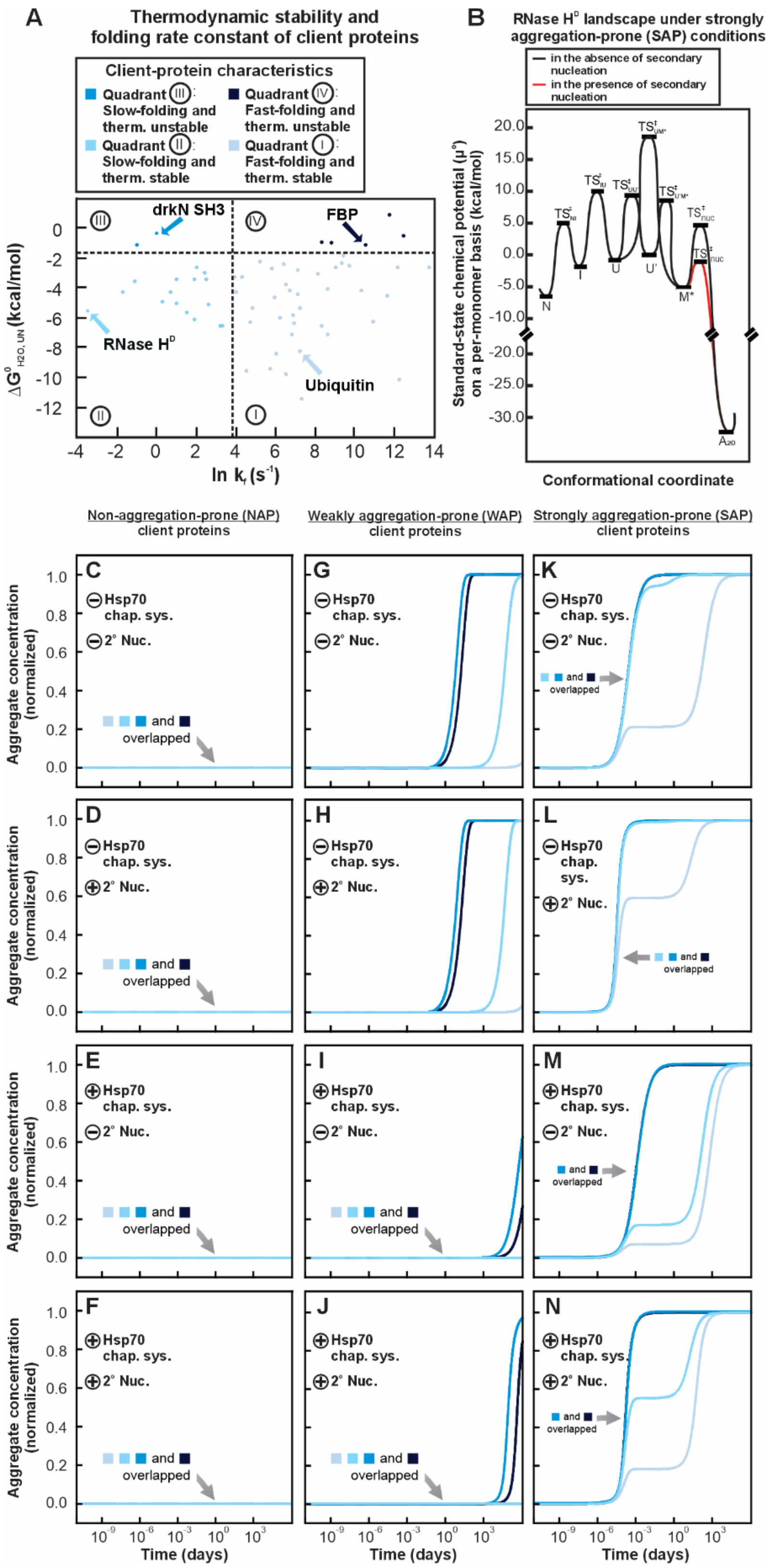

A
Thermodynamic stability and folding rate constant of client proteins
Client-protein characteristics
Quadrant III: Slow-folding and therm. unstable
Quadrant IV: Fast-folding and therm. unstable
Quadrant II: Slow-folding and therm. stable
Quadrant I: Fast-folding and therm. stable
drkN SH3
FBP
RNase H^D
Ubiquitin
ln k_f (s^-1)
B
RNase H^D landscape under strongly aggregation-prone (SAP) conditions
in the absence of secondary nucleation
in the presence of secondary nucleation
Standard-state chemical potential (μ°) on a per-monomer basis (kcal/mol)
Conformational coordinate
Non-aggregation-prone (NAP) client proteins
Weakly aggregation-prone (WAP) client proteins
Strongly aggregation-prone (SAP) client proteins
Aggregate concentration (normalized)
Hsp70 chap. sys.
2° Nuc.
and overlapped
Time (days)

**Fig. S16. Computationally predicted role of monomer-dependent secondary nucleation on aggregation outcomes in the absence and presence of Hsp70 chaperone system. A.** Diagram illustrating the thermodynamic and kinetic characteristics of 61 small client proteins sorted according to their known thermodynamic stability and folding rates.[14] Four proteins were selected (various shades of blue) as representative chaperone clients for each of the four quadrants. All the CHAMPION70 simulations began with 15 μM client-protein in the conformationally unrelaxed denatured unfolded state (U'). Simulations in the presence of chaperones included 25 μM DnaK, 5 μM DnaJ, 10 μM GrpE, 3.5 mM ATP, 0.12 mM ADP, and 5 mM $P_i$. **B.** Standard-state free-energy diagram for RNase $H^D$ model client-protein under strongly aggregation-prone (SAP) conditions, shown on a per-monomer basis. Aggregation pathway in red highlights how the presence of monomer-dependent secondary nucleation affects the kinetic accessibility of aggregate nucleation. **C, G, K.** Simulated formation of fully aggregated $A_{20}$ species in the absence of chaperones and the absence of secondary nucleation for clients with three aggregation propensities: non-aggregation-prone (NAP) (**C**), weakly aggregation-prone (WAP) (**G**), and strongly aggregation-prone (SAP) (**K**), shown for each of the four representative clients from each panel-A region. **D, H, L.** Simulated formation of $A_{20}$ in the absence of chaperones and the presence of secondary nucleation for non-aggregation-prone (NAP) (**D**), weakly aggregation-prone (WAP) (**H**), and strongly aggregation-prone (SAP) (**L**) client-proteins. **E, I, M.** Simulated formation of $A_{20}$ in the presence of chaperones and the absence of secondary nucleation for non-aggregation-prone (NAP) (**E**), weakly aggregation-prone (WAP) (**I**), and strongly aggregation-prone (SAP) (**M**) client-proteins. **F, J, N.** Simulated formation of $A_{20}$ in the presence of chaperones and the presence of secondary nucleation for non-aggregation-prone (NAP) (**F**), weakly aggregation-prone (WAP) (**J**), and strongly aggregation-prone (SAP) (**N**) client-proteins.

### Fig. S17

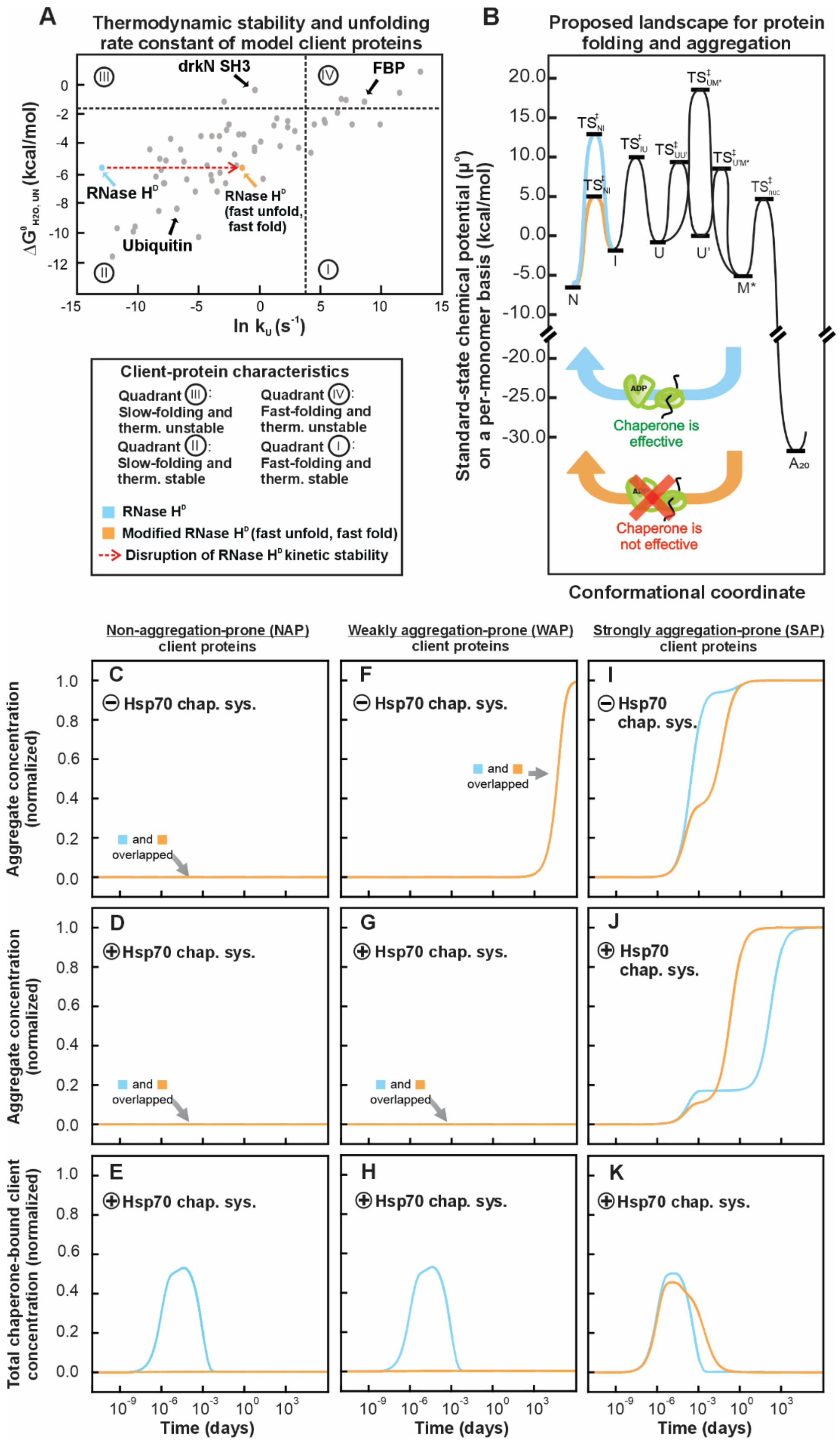
A
Thermodynamic stability and unfolding rate constant of model client proteins
drkN SH3
FBP
RNase H$^D$
RNase H$^D$ (fast unfold, fast fold)
Ubiquitin
$\Delta G^0_{H2O, UN}$ (kcal/mol)
ln $k_u$ ($s^{-1}$)
Client-protein characteristics
Quadrant III: Slow-folding and therm. unstable
Quadrant IV: Fast-folding and therm. unstable
Quadrant II: Slow-folding and therm. stable
Quadrant I: Fast-folding and therm. stable
RNase H$^D$
Modified RNase H$^D$ (fast unfold, fast fold)
Disruption of RNase H$^D$ kinetic stability
B
Proposed landscape for protein folding and aggregation
Standard-state chemical potential ($\mu^0$) on a per-monomer basis (kcal/mol)
Chaperone is effective
Chaperone is not effective
Conformational coordinate
Non-aggregation-prone (NAP) client proteins
Weakly aggregation-prone (WAP) client proteins
Strongly aggregation-prone (SAP) client proteins
Aggregate concentration (normalized)
Total chaperone-bound client concentration (normalized)
Hsp70 chap. sys.
and overlapped
Time (days)

**Fig. S17. Computationally predicted role of kinetic stability upon client-protein interaction with the Hsp70 chaperone system. A.** Diagram illustrating the thermodynamic and kinetic characteristics of 61 small client proteins sorted according to their known thermodynamic stability and folding rate constants.[14] RNase $H^D$ was modified by adjusting its folding and unfolding rate constants so that the native-state retained the same chemical potential but folded substantially faster (see Supplementary Table 3 for rate constants). All the CHAMPION70 simulations began with 15 µM client-protein in the conformationally unrelaxed denatured unfolded state (U'). Simulations in the presence of chaperones included 25 µM DnaK, 5 µM DnaJ, 10 µM GrpE, 3.5 mM ATP, 0.12 mM ADP, and 5 mM $P_i$. **B.** Standard-state free-energy diagrams for the original RNase $H^D$ (blue) and the modified fast-fold/fast-unfold RNase $H^D$ variant (orange), shown on a per-monomer basis. The altered U to N barrier height reflects the change in kinetic stability. **C, F, I.** Simulated formation of fully aggregated $A_{20}$ species in the absence of chaperones for clients with three aggregation propensities: non-aggregation-prone (NAP) (**C**), weakly aggregation-prone (WAP) (), and strongly aggregation-prone (SAP) (**I**), shown for both the original RNase $H^D$ and the fast-fold/fast-unfold RNase $H^D$. **D, G, J.** Simulated formation of $A_{20}$ in the presence of chaperones for non-aggregation-prone (NAP) (**D**), weakly aggregation-prone (WAP) (**G**), and strongly aggregation-prone (SAP) (**J**) client-proteins. **E, H, K.** Simulated association of clients within the Hsp70 cycle for non-aggregation-prone (NAP) (**E**), weakly aggregation-prone (WAP) (**H**), and strongly aggregation-prone (SAP) (**K**) client proteins.

**Fig. S18**

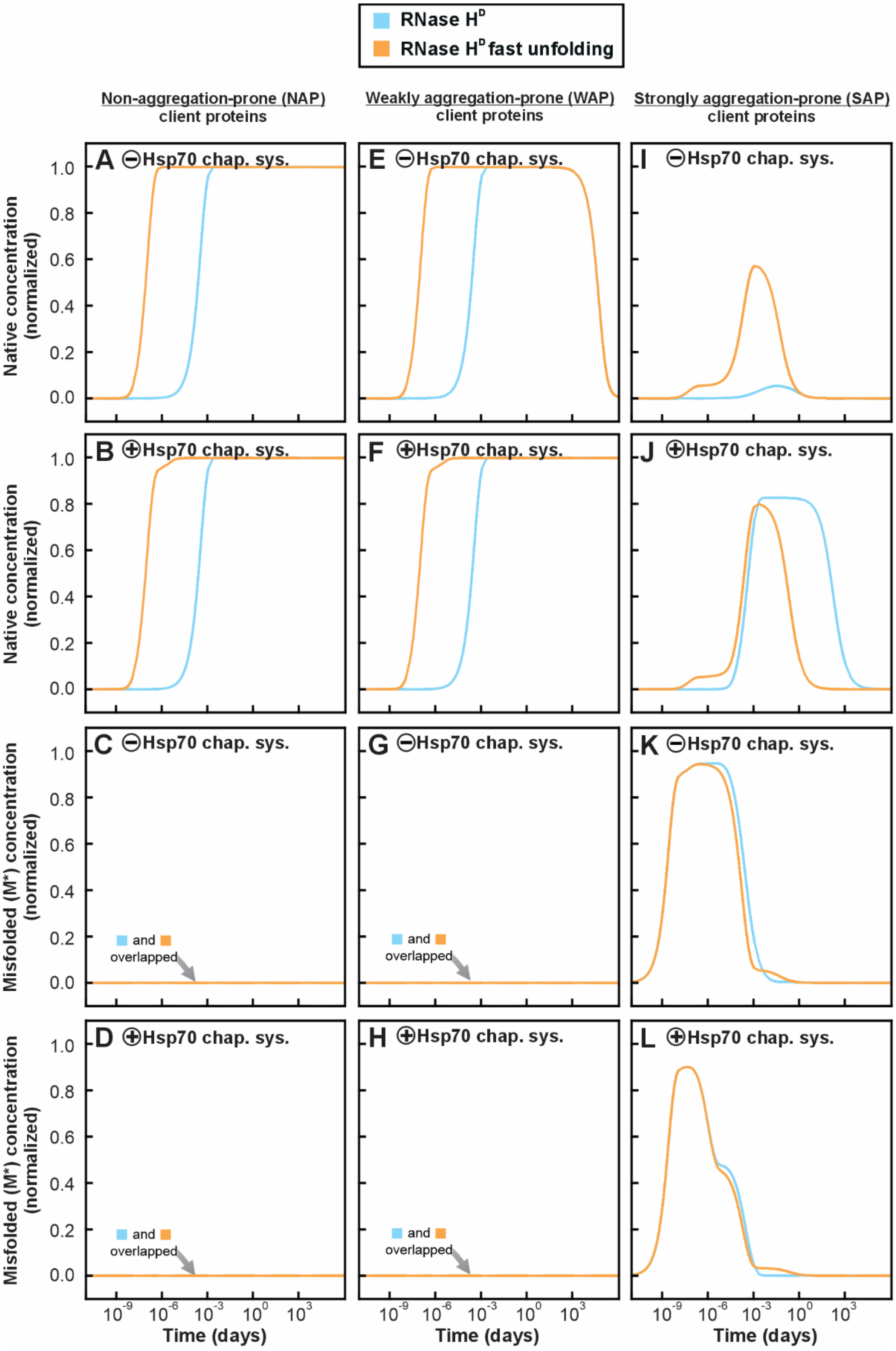

RNase H^D
RNase H^D fast unfolding
Non-aggregation-prone (NAP) client proteins
Weakly aggregation-prone (WAP) client proteins
Strongly aggregation-prone (SAP) client proteins
A ⊖Hsp70 chap. sys.
E ⊖Hsp70 chap. sys.
I ⊖Hsp70 chap. sys.
B ⊕Hsp70 chap. sys.
F ⊕Hsp70 chap. sys.
J ⊕Hsp70 chap. sys.
C ⊖Hsp70 chap. sys.
G ⊖Hsp70 chap. sys.
K ⊖Hsp70 chap. sys.
D ⊕Hsp70 chap. sys.
H ⊕Hsp70 chap. sys.
L ⊕Hsp70 chap. sys.
Native concentration (normalized)
Misfolded (M*) concentration (normalized)
and overlapped
1.0
0.8
0.6
0.4
0.2
0.0
10^-9 10^-6 10^-3 10^0 10^3
Time (days)

**Supplementary Fig. S18. Computationally predicted role of kinetic stability on native-state formation and misfolding timecourses.** All the CHAMPION70 simulations began with 15 μM client-protein in the conformationally unrelaxed denatured unfolded state (U'). Simulations in the presence of chaperones included 25 μM DnaK, 5 μM DnaJ, 10 μM GrpE, 3.5 mM ATP, 0.12 mM ADP, and 5 mM $P_i$. **A, E, .** Simulated formation of the native-state (N) in the absence of chaperones for clients with three aggregation propensities: non-aggregation-prone (NAP) (**A**), weakly aggregation-prone (WAP) (**E**) and strongly aggregation-prone (SAP) (**I**), shown for both the original RNase $H^D$ and the fast-fold/fast-unfold RNase $H^D$. **B, F, J.** Simulated formation of N in the presence of chaperones for non-aggregation-prone (NAP) (**B**), weakly aggregation-prone (WAP) (**F**), and strongly aggregation-prone (SAP) (**J**) client-proteins. **C, G, K.** Simulated formation of the misfolded-state (M*) in the absence of chaperones for non-aggregation-prone (NAP) (**C**), weakly aggregation-prone (WAP) (**G**), and strongly aggregation-prone (SAP) (**K**) client-proteins. **D, H, L.** Simulated formation of M* in the presence of chaperones for non-aggregation-prone (NAP) (), weakly aggregation-prone (WAP) (**H**), and strongly aggregation-prone (SAP) (**L**) client-proteins.
